# A Litmus Test for Confounding in Polygenic Scores

**DOI:** 10.1101/2025.02.01.635985

**Authors:** Olivia S. Smith, Samuel Pattillo Smith, Dandan Peng, Xinyi Miao, Hakhamanesh Mostafavi, Jeremy J. Berg, Michael D. Edge, Arbel Harpak

**Author notes:** Correspondence should be addressed to M.D.E and A.H. These authors contributed equally to this work (Olivia S. Smith, Samuel Pattillo Smith, Dandan Peng).

## Abstract

Polygenic scores (PGSs) are being rapidly adopted for trait prediction in the clinic and beyond. PGSs are often thought of as capturing the direct genetic effect of one’s genotype on one’s phenotype. However, because PGSs are constructed from population-level associations, they are influenced by factors other than direct genetic effects, including stratification, assortative mating, and dynastic effects (“SAD effects”). Our interpretation and application of PGSs may hinge on the relative influence of SAD effects, since they may often be environmentally or culturally mediated. We developed a method to measure these influences, Partitioning Genetic Scores Using Siblings (PGSUS, pron. “Pegasus”). PGSUS leverages a comparison of a PGS of interest based on a standard GWAS with a PGS based on a sibling GWAS—which is largely immune to SAD effects—to partition variance in a PGS (in a given sample) into components due to direct effects, SAD effects, and their covariance. Using PGSUS, we found that in many cases direct genetic effects contribute relatively little to PGS variation—most pronouncedly so in PGSs for social or behavioral traits, such as educational attainment or neuroticism. PGSUS further breaks down variance components by axes of genetic ancestry, allowing for a nuanced interpretation of SAD effects. In particular, PGSUS can detect stratification along major axes of ancestry as well as SAD variance that is “isotropic” with respect to axes of ancestry. Applying PGSUS, we found evidence of stratification in PGSs constructed using large meta-analyses of height as well as in multiple PGSs constructed using the UK Biobank. We show that a given PGS can suffer from stratification along a major axis of ancestry in one sample but not in another (for example, in comparisons of prediction in samples from contemporary vs. ancient DNA samples). We further show that when axes of stratification are shared between GWAS and prediction samples, stratification can both aid and impede phenotypic prediction accuracy. In summary, PGSUS offers advances in interpretation towards more informed application of polygenic scores.

## Introduction

Genome-wide association studies (GWASs) in humans have revealed that the genetic basis of variation for many health conditions and other traits is highly polygenic. That is, most variation derives from a large number of common genetic variants with effects that are individually small. GWASs further revealed that the joint effect of these variants is often well-captured by a simple linear combination, consistent with longstanding theoretical models ^1–3^. Following from these observations, attention has turned toward the construction of genomic predictors of traits, so-called “polygenic scores” (PGSs). A PGS for a trait is typically a sum over a set of loci of an individual’s alleles, each weighted by its GWAS-estimated effect.

PGSs have many potential uses, including in the clinic ^4,5^, as instruments to account for genetic variation when studying environmental effects ^6^, and for studying human evolution ^7,8^. However, GWAS and polygenic scores in humans suffer from a general weakness: a lack of experimental control. Neither environments nor genetic backgrounds can be randomized among individuals within GWAS samples. When a specific locus’s alleles are correlated with other causal factors (genetic, societal, or environmental), they become spuriously associated with the phenotype. This can bias allelic effect estimates with respect to the causal genetic effect of the focal locus (or, more precisely, of linked causal variants)^9–11^.

What factors contribute to standard-GWAS allelic effect estimates? It has long been established that, in addition to the causal effects of one’s alleles on one’s own phenotype (“direct” genetic effects), allelic effect estimates can be affected by confounding due to population stratification ^9,12^. More recently, other phenomena that affect GWAS allelic effect estimates have received increased attention ^13,14^, such as assortative mating ^14–18^ and “dynastic” (or indirect parental) genetic effects. Dynastic effects arise by virtue of the correlation between an individual’s genotype and that of their parent, when the parent’s genotype influences the focal individual’s phenotype ^19–22^. For example, if maternal genotypes affect the focal individual’s uterine environment during early embryonic development, then there will be an association between the focal individual’s genotype and traits that depend on the uterine environment, even if the genotype only operates via the mother. We henceforth refer to these influences as “SAD” (Stratification, Assortment, and Dynastic) effects, though there may be additional systematic influences on GWAS allelic effect estimates beyond these three factors (e.g., ascertainment biases such as “winner’s curse”). Although GWAS analysis pipelines include steps aimed at adjusting for environmental and genetic confounding, even residual SAD effects that are small on a per-SNP basis can combine to have large effects on the variance of a PGS across individuals ^11,23–26^.

When considering the application of a PGS to individual genomes, it may be important to know the extent to which variation among individuals’ PGSs is due to SAD effects. Yet, to our knowledge, there are no methods that address this question. To illustrate the importance of this task, consider an example in which variation among individuals in a PGS for cancer is largely due to correlates of SNPs included in the PGS (index SNPs) in the GWAS sample, such as exposure to pollutants. The PGS may nonetheless be predictive—this would depend on whether the genotype-exposure correlation in the GWAS sample exists in the prediction sample. If a PGS is predictive largely because of SAD effects, should we use the PGS as a predictor of genetic risk? The answer may vary across different applications of PGSs. Regardless, a diagnostic tool measuring the impact of SAD factors could provide grounds for an informed decision.

The extent to which variation in a given PGS is attributable to SAD effects may also differ among prediction samples. Therefore, understanding the relationship between SAD effects in the GWAS sample and characteristics of the prediction sample may help us understand PGS portability^25,27,28^.

One promising way to evaluate the impact of SAD effects on a PGS is family-based designs, such as sibling GWAS, in which trait differences among pairs of full siblings are regressed on differences in their genotypes. Sibling GWASs are largely robust to SAD effects ^25,29–31^, though they are not strictly immune ^18^ and complexities can arise in the presence of genetic interactions or other factors ^14^. Family-based designs are typically underpowered compared with population-based GWAS, and are therefore rarely used to build PGS for anything but research purposes. Nevertheless, family studies can help us distinguish direct genetic effects from SAD effects contributing to standard-GWAS estimates.

Here, we develop a method, Partitioning Genetic Scores Using Siblings (PGSUS, pron. “Pegasus”), that partitions the variance of a PGS derived from a population-based GWAS (“standard GWAS” below) on the basis of a comparison with the corresponding allelic effect estimates from a sibling-based GWAS (“sib-GWAS” below). PGSUS goes beyond previous approaches by quantifying the contribution of direct genetic effects versus SAD effects (and their covariance) to a PGS as applied in a given prediction sample. PGSUS can be viewed as a litmus test for the presence of confounding and can therefore inform specific applications and interpretations of PGSs.

## Results and Discussion

### Model

We consider a PGS applied to a “prediction sample”—a sample of genotypes for which the PGS is calculated. Our main aims are to quantify the extent to which PGS variance is driven by direct effects and to identify components of SAD variance aligning with specific axes of ancestry, suggestive of stratification along that axis (see **A litmus test for stratification**). We begin by outlining our model, which motivates the partitioning of PGS variance that PGSUS performs.

#### PCA partitioning of polygenic score variance

The variance of a PGS can be partitioned into orthogonal components attributable to principal components of the genotype matrix of the prediction sample. In **Text S1**, we show that

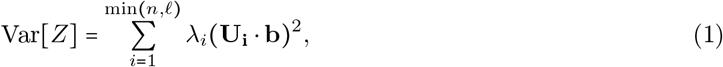

where *Z* is a random draw from a sample of *n* PGS values, *λ*_*i*_ is the eigenvalue for the *i*-th principal component of the prediction sample’s genotype matrix, **U**_**i**_ is a vector holding the loadings of index SNPs on principal component *i, b* is a vector of allelic effect estimates, and · represents the dot product. *n* is the number of individuals in the sample used to perform the PCA and *ℓ* is the number of index SNPs. This variance partitioning is equivalent to the partitioning that would be obtained by regressing individual PGS values on individual-level principal-component coordinates but calculated via a projection of the SNP allelic effect estimates onto the PCs. Thanks to this equivalence, PGSUS does not require any individual-level data, but only GWAS summary statistics and a PCA of the prediction sample (or a representative sample from the population to which the researcher wishes to apply the PGS; **Fig. 1B**).

**Figure 1.**
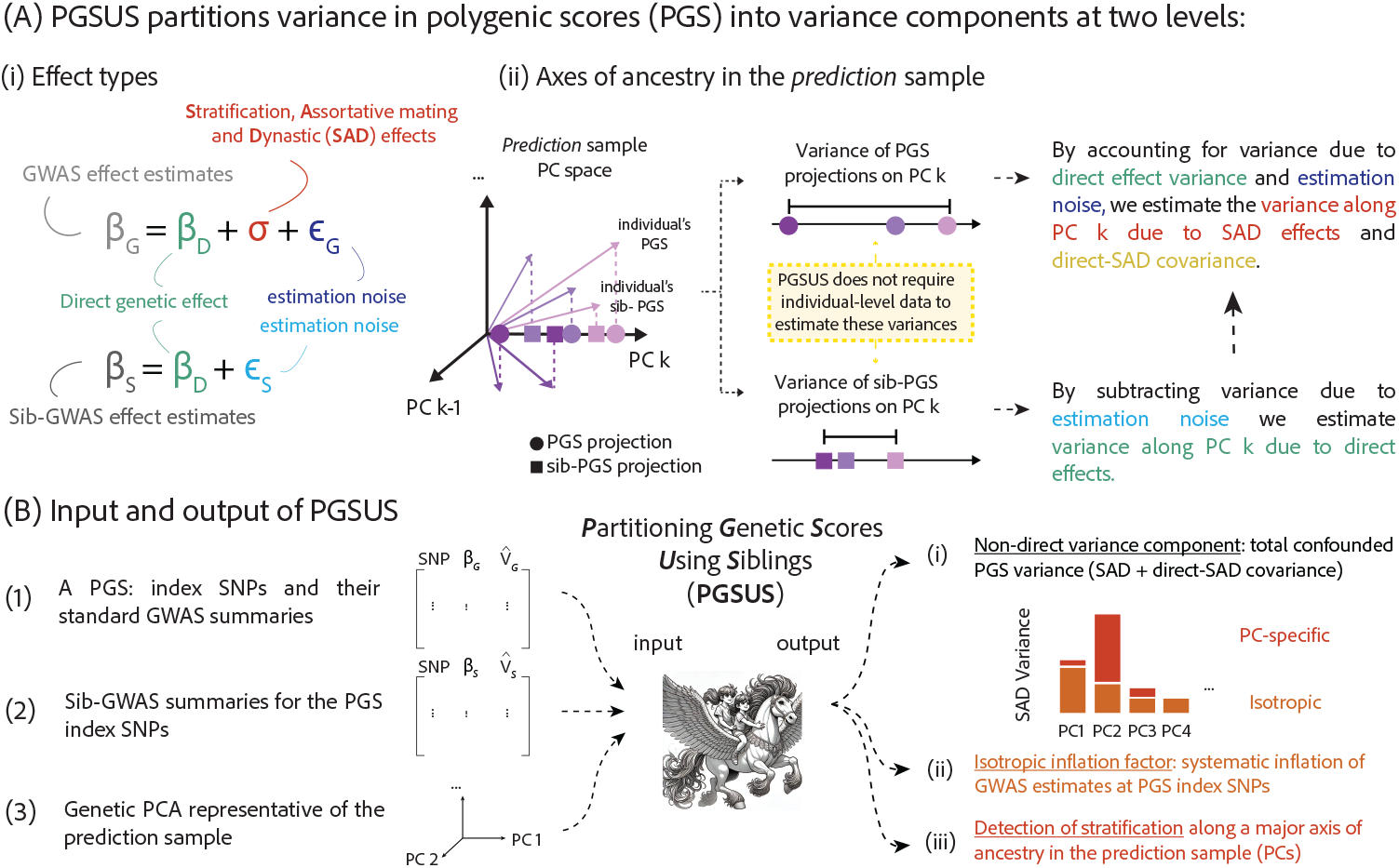
Partitioning Genetic Scores Using Siblings (PGSUS). **(A)** PGSUS partitions the variance of a PGS among individuals in a prediction sample at two levels. Namely, variance components are attributable to both (i) effect type: direct effects, SAD effects, and their covariance, as well as estimation noise; and (ii) a principal component of the genotype matrix of the prediction sample. The partitioning by effect types derives from the generative model shown in (i) for standard-GWAS effect estimates and the corresponding sib-based estimates. **(B)** PGSUS does not require individual-level data. It only requires an input of GWAS summary statistics (allelic effect estimates and associated standard errors), the corresponding sib-GWAS summary statistics, and a PCA (namely, eigenvalues and eigenvectors) of a genotype matrix representative of the sample to which the PGS is to be applied. The output is a set of variance components. These are also reported as three key summaries: (i) the summed contribution of all SAD variance and direct-SAD covariance divided by total PGS variance, a measure of relative PGS variance due to confounding, i.e. beyond direct effects alone, (ii) an estimate of “isotropic” inflation of allelic effect estimates at PGS index SNPs, due to SAD effects or other factors, e.g., ascertainment-related effects, that are not specific to a PC (as the cartoon shows, isotropic inflation is not assumed to result in equal SAD variance on each PC, but instead in variance proportional to the direct effect variance component of each PC, in expectation), and (iii) PCs with significantly large PC-specific SAD variance, suggestive of stratification along the corresponding PC.

In addition, expressing the variance partitioning in this way allows us to link it to a generative model for the allelic effects (see below). The quantity **U**_**i**_ ⋅ **b** is the scalar projection of the allelic effect estimates **b** on vector **U**_**i**_ and can be thought of as the effect estimated for principal component *i*. It is also closely related to the covariance of the vector of allelic effect estimates with **U**_**i**_.

#### Generative model of allelic effect estimates

We model the allelic effect estimates from a standard GWAS, *β*_*G*_, as the sum

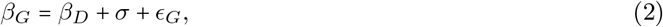

where *β*_*D*_ is the true direct genetic effect, *σ* represents SAD effects, and *ϵ*_*G*_ is a measurement error (**Fig. 1A**). At each locus, we assume that *ϵ*_*G*_ has expectation zero and standard deviation equal to the standard error of *β*_*G*_ as estimated in the GWAS. In a sib-GWAS, we model allelic effect estimates as

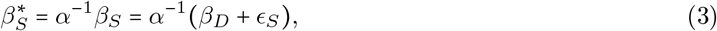

where *α*^−1^*ϵ*_*S*_ is the measurement error, again with expectation zero and standard deviation given by the estimated standard error of 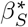 (**Text S6**). The parameter *α*, the isotropic inflation factor, captures systematic differences in the magnitude of allelic effect estimates between the standard GWAS and sib-GWAS. In the section **Isotropic inflation and non-direct variance**, we discuss both SAD factors and technical issues that may contribute to isotropic inflation and the estimation of *α*.

We highlight the assumption that the direct effect *β*_*D*_ is the same in standard- and sib-GWAS-based allelic effect estimates. This may not be true in general, especially in the presence of differences in ancestry, environment and social context between the standard and the sibling samples ^14,25^. (Even without such differences, and no SAD effects, there are differences in the expectations of *β*_*D*_ and *β*_*S*_ ^14^—but they are likely to be small, see **Text S2**).

#### Partitioning of PGS variance by both effect types and PCs

In combination, the expression for the variance of a PGS in **Eq. 1** and the models for the allelic effect estimates contributing to a PGS in **Eqs. 2,3** suggest a partitioning of variance in a PGS due to the projections of direct effects, SAD effects, and measurement error on each principal component. Specifically, suppose a vector of the allelic effect estimates from GWAS at index SNPs, 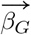, replaces **b** in **Eq. 1**, as in

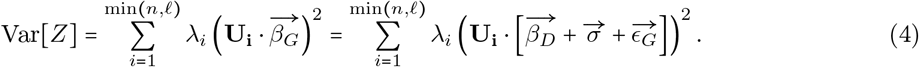

Distributing the **U**_**i**_ and expanding the expression on the right gives

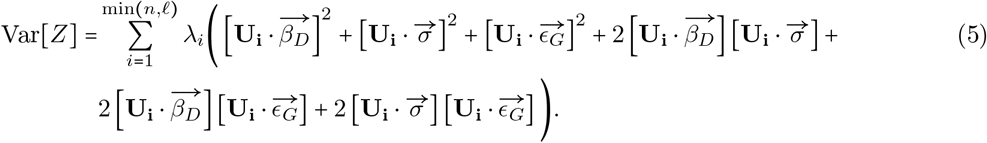

We are principally interested in estimating three of the terms that arise from the expansion in **Eq. 5**, which can be interpreted as variance components attributable to direct effects, to SAD effects, and to the covariance of direct effects and SAD effects. In particular,

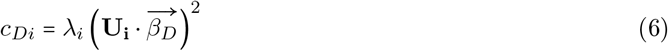

is the component of variance in the PGS due to direct effects specific to principal component *i* of the genotype matrix, and

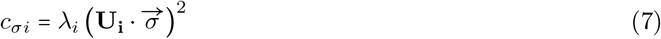

is the component of variance due to SAD effects specific to principal component *i*. We also consider

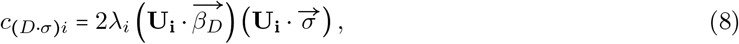

which is a contribution to variance due to covariance of direct and SAD effects specific to principal component *i*. This last term can be negative, unlike the previous two, which are constrained to be non-negative. The combination of SAD variance and direct-SAD covariance on PC *i* gives the PC-specific non-direct variance component,

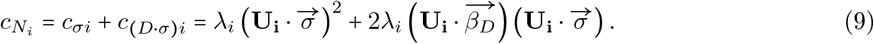

Summing over all PCs and dividing by the total variance in the PGS,

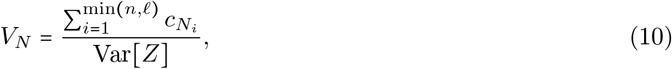

we get a measure of the relative contribution of confounded effects—more precisely, effects other than strictly direct effects—to PGS variance, which we henceforth refer to as the “non-direct variance component”.

In the section **PGSUS-based parameter estimation**, we derive estimators. Our basic strategy is to isolate 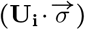 using the projection of the difference between allelic effect estimates from the standard GWAS and those from the sib-GWAS, and proceed with moment-based estimation.

We developed PGSUS as freely available software that performs this partitioning. The software takes three inputs (**Fig. 1B**). It reports all PGS variance components alongside three key summaries of interest (**Fig. 1B**), which we will focus on in our discussion:

i. the non-direct variance component,
ii. the isotropic inflation factor, *α*, and
iii. PC-specific, effect type-specific subcomponents.

In order to produce this output, PGSUS performs two consecutive partitionings for complementary purposes. First, we set *α* ≔ 1 in **Eq. 3** and perform the partitioning to estimate (i) and (ii). The estimate of the non-direct variance component therefore reflects the relative contribution of confounding due to both PC-specific and isotropic effects (see the sections **Overview of parameter estimation** and **Text S13**). The estimated isotropic inflation factor, 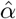(see the section **Estimation of the isotropic inflation factor**) is used to rescale sib-based allelic effect estimates and standard errors, multiplying each by 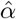,

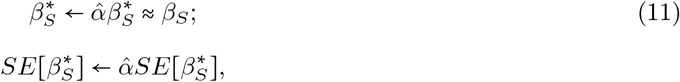

such that they are at the same scale as standard-GWAS-based estimates. We use these rescaled sibbased summaries as input for a second partitioning of the same PGS. The variance components from this second partitioning, and (iii) in particular, are reported. As a result, our estimates of PC-specific variance components are adjusted for isotropic inflation (in practice, results based on (iii) tended to be largely insensitive to this two-step strategy, which we elaborate on in **Text S13**; **Figs. S1-S3**). Below, we discuss how the outputs of PGSUS can be used as diagnostics of confounding.

### The relative contribution of confounding to variation in a PGS

We partitioned the variance of a suite of PGSs constructed using GWASs we performed in the UK Biobank ^32^ (UKB). For concreteness, we first focus on a set of GWASs performed with a standard adjustment for 20 principal components (PCs) of the GWAS sample genotype matrix and PGSs constructed through clumping and thresholding ^33–35^ for 12 highly heritable physiological traits and 5 additional social or behavioral traits (**Table S1**). All GWASs were performed using the self-identified White British subset of the UKB (UKB WB). We used PGSUS to partition variances in the PGSs with respect to either the entire 1000 Genomes Phase 3 sample (1KG) ^36^, the subsample of Europeans in 1KG (1KG Europeans), the complement of the White British subsample in UKB (UKB NWB), or to a Eurasian subsample of the Allen Ancient DNA Resource (AADR) ^37^ as the prediction sample.

We begin by examining the non-direct variance component as a measure of the relative contribution of confounded variance to PGS variation in a prediction sample. Focusing first on 1KG Europeans as the prediction sample with a GWAS ascertainment threshold of *p*-value < 10^−8^, the non-direct variance component for 16 PGSs ranged from 0.25 to 0.85 (averaged at 0.57), with the highest values observed for four social and behavioral traits—neuroticism score (0.85), years of schooling (0.84), pack years of smoking (0.82), and household income (0.81)—and the lowest for diastolic blood pressure (0.25) and hip circumference (0.25) (**Fig. 2A**). Other studies of social and behavioral traits reported inflation in the estimated magnitude of standard-GWAS-estimated allelic effects ^13,38,39^ or trait variance explained ^25^ compared with family studies; however, our estimates are, to our knowledge, the first estimates of the contribution of confounding to variance in these PGSs.

**Figure 2.**
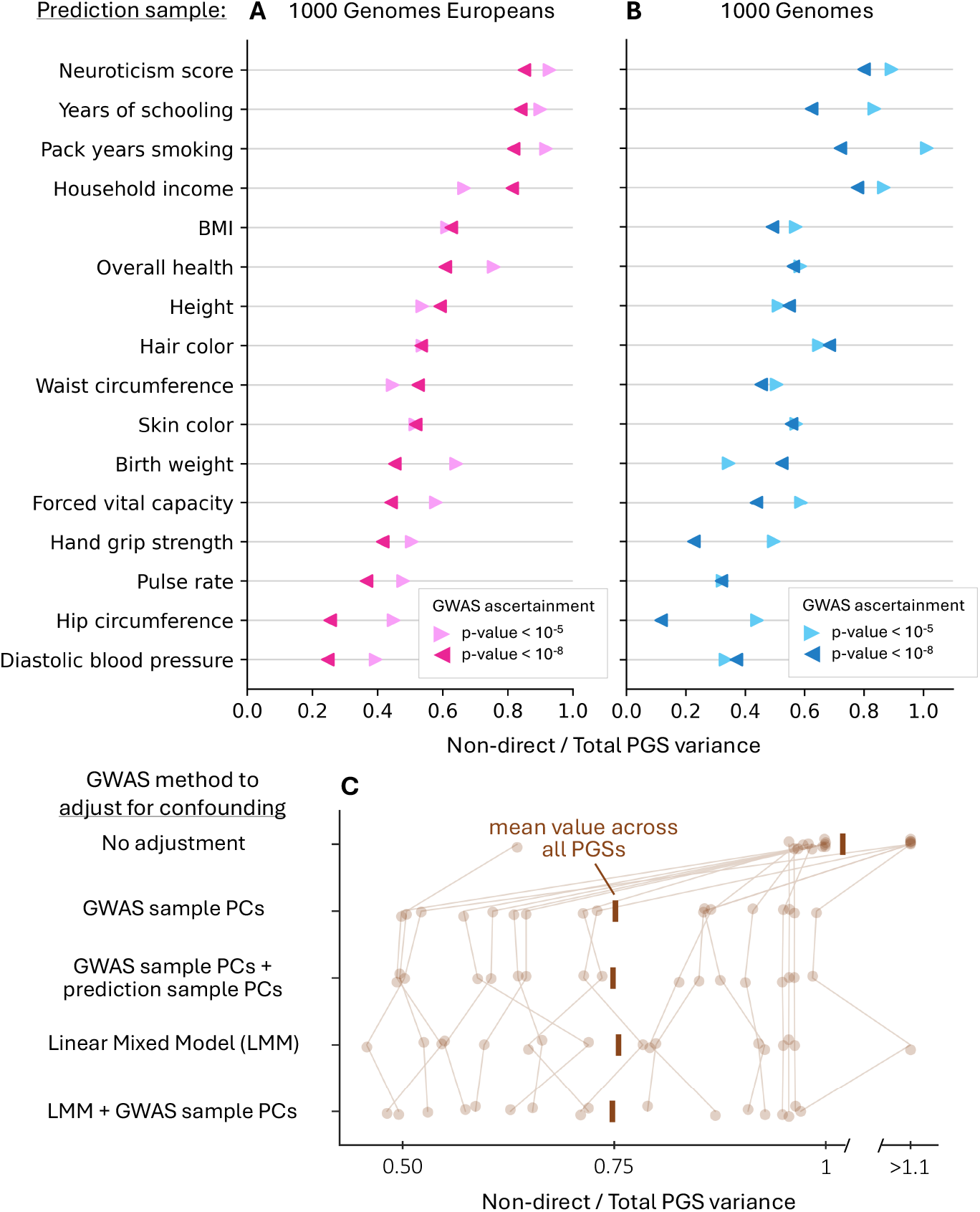
Contribution of non-direct variance to PGS variance. The “non-direct variance component” shown on the x-axis is the ratio of the sum of the SAD variance and direct-SAD covariance components and the total PGS variance. It is a measure of the relative contribution of confounding to PGS variance. Shown here are estimates for PGS variance in two distinct prediction samples, constructed from a GWAS in the White British subset of the UK Biobank using clumping and thresholding (two PGSs using different thresholding choices shown per trait). **(C)** We performed GWASs for each of 17 traits (ascertained at *p*-value < 10^−5^) in a random sample of UK Biobank individuals, using a different method to adjust for population structure in each GWAS. We then partitioned the PGS variance in the 1KG Europeans prediction sample and estimated the non-direct variance component. “GWAS sample PCs” refers to a GWAS adjustment for the first 20 principal components of the genotype matrix of the cohort in which the GWAS is performed. “GWAS sample PCs + prediction sample PCs” refers to adjustment for the first 20 principal components of the genotype matrix of both the GWAS cohort and the prediction sample. “LMM” refers to the *BOLT-LMM* implementation of a linear mixed model. Vertical lines connect PGSs for the same trait across adjustment methods. The mean value across all PGSs for a given GWAS adjustment method is shown as a large, opaque vertical bar.

As we will discuss in later sections, other diagnostics output by PGSUS depend heavily on GWAS and polygenic scoring methodology as well as the prediction sample. For the non-direct variance component, however, the magnitude and the order across traits was similar when we partitioned variance in the same PGSs but in a different prediction sample—1KG (Pearson *r =* 0.86, *p =* 1.9 × 10 ^−5^; **Fig. 2B**). The order was also highly similar (Pearson *r* = 0.86, *p =* 1.8 × 10 ^−5^), and the non-direct variance components were typically larger, when using a less stringent GWAS ascertainment for PGS index SNPs (GWAS *p*-value < 10 ^−5^; lighter colors in **Fig. 2A,B**).

Are GWAS adjustments for population structure successful in mitigating confounded variance? The PGSs discussed above were based on GWASs in which we adjusted allelic effect estimates by including the top 20 PCs of the GWAS sample’s genotype matrix as covariates in the GWAS—a widely used approach ^23,24,40,41^. We found that this approach and other GWAS adjustments, such as linear mixed models (LMMs), reduced non-direct variance to a similar extent (paired t-test *p >* 0.2 in pairwise comparisons across methods; **Figs. 2C, S4-S6; Text S9; Files S9, S10**). When the GWAS sample had relatively little genetic structure, i.e. in the same UKB WB sample, these adjustments did not significantly reduce non-direct variance components compared with a GWAS with no adjustment at all (**Fig. S6**). In a more structured sample (namely, a subset of the full UKB sample), all adjustment methods reduced non-direct variance on average by 23% compared with a GWAS with no adjustment (paired t-test *p*<0.004 for all adjustment method pairings, **Figs. 2C, S4**). Nonetheless, the lion’s share of PGS variance was still non-direct in most cases.

### A litmus test for stratification

If an axis of stratification largely overlaps with a PC of the prediction sample, we expect to see a large PC-specific SAD variance component. We confirmed this expectation with a simulation study (**Text S14.4, Fig. S7**). Here, we turn to an empirical case in which stratification is already known to contribute to PGS variance. In 2019, researchers discovered that the GIANT consortium’s height GWAS ^42^ was confounded along a north-south ancestry axis in Europe ^23,24^. We constructed a PGS using this GWAS and partitioned its variance in 1KG Europeans as the prediction sample. As expected, the SAD variance component on PC2 (which best tags the north-south ancestry axis ^43^; **Fig. 3A** inset) is large—28% of the size of the total variance of the PGS (**Fig. 3A**). This variance component is significantly larger than expected under a null model with no PC-specific SAD variance (*p =* 0.0005; 95% null non-rejection regions are shown in **Fig. 3**; see **Text S3** for details on the empirical null).

**Figure 3.**
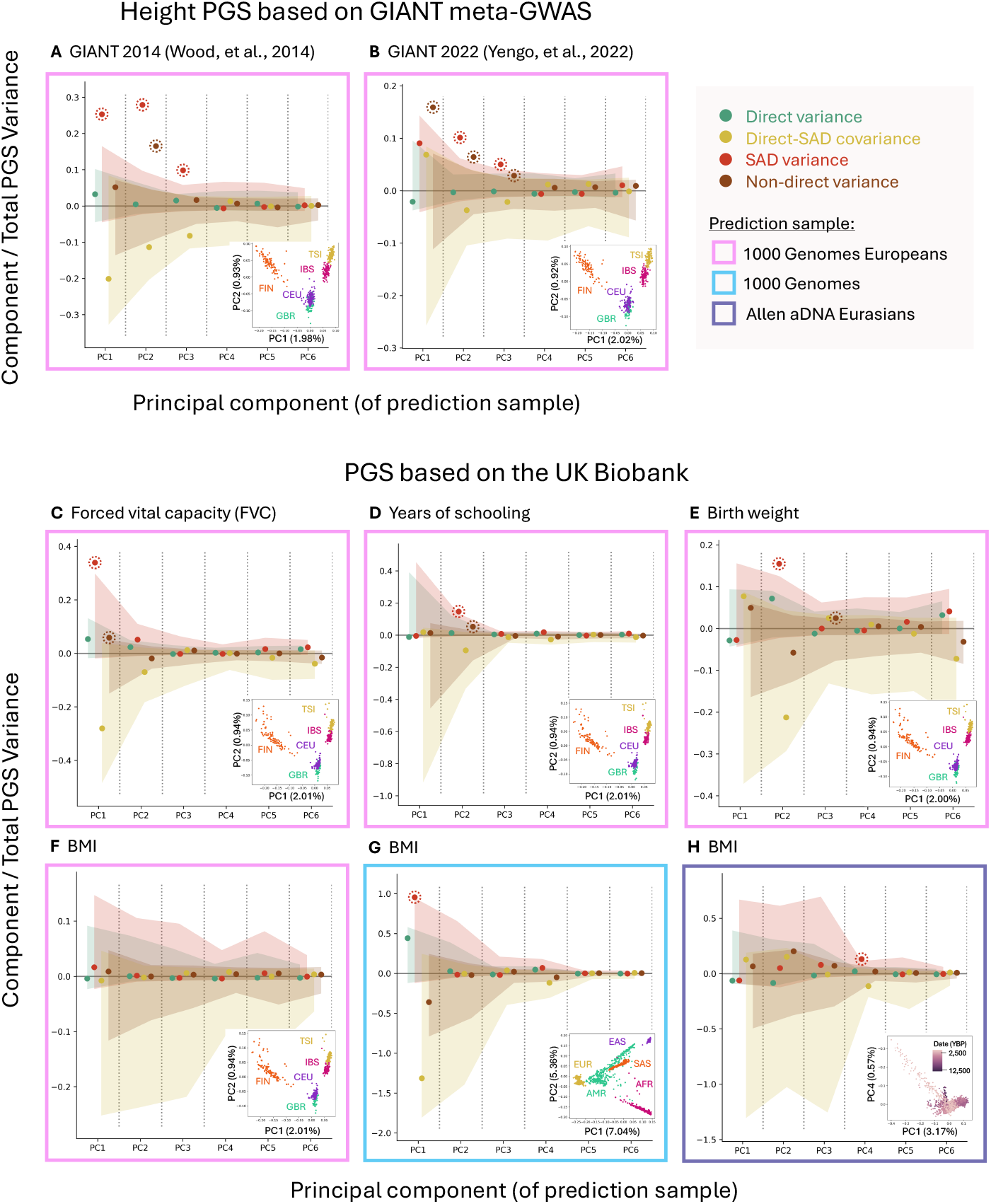
PGSUS partitioning uncovers stratification in polygenic scores. For eight examples, we show variance components along the first six PCs divided by the total variance in PGS in the prediction sample. Shaded regions show empirical permutation-based null non-rejection regions. Components deviating from this expectation are highlighted with dashed circles. Insets show the prediction-sample individual coordinates along some of the PCs on which we observed significantly large variance components; labels are shorthand for subsamples in the 1000 Genomes (1KG) data or the Allen Ancient DNA Resource Eurasian subset. Insets vary slightly between the same prediction cohort as they are constructed using the set of SNPs selected through clumping using trait-specific summary statistics. The 1KG subsample labels in panels **(A-F)** correspond to the five European subpopulations: Finnish in Finland (FIN), Utah residents with northern and western European ancestry (CEU), Iberian populations in Spain (IBS), British from England and Scotland (GBR), and Tuscan in Italy (TSI). The 1KG subsample labels in panel **(G)** correspond to the five 1KG superpopulations: European (EUR), admixed Americans (AMR), East Asian (EAS), South Asian (SAS), and African (AFR). Date (YBP) in panel **(H)** refers to the sample date estimate in years before present. Percentages of variance explained by each PC are given in the parentheses of each axis label in the insets. All PGSs shown are constructed using clumping and thresholding with marginal association thresholds as follows: *p* < 1 for panel **(D)**; *p* < 10^−5^ for panels **(A, C, E, F, H)**; *p* < 10^−8^ for panels **(B, G).** PGSs partitioned in panels **(C-H)** are based on GWASs with 20 GWAS sample PCs as covariates as the only adjustment for population structure.

This result confirms that PGSUS can detect a “positive control”, a known example of confounding along a major axis of genetic ancestry. Importantly, results produced by PGSUS tell us about variation in a specific PGS, not about a trait or even about a specific GWAS for that trait. To demonstrate this, we considered partitionings of other height PGSs with respect to the same prediction sample. For a similarly constructed PGS based on a GWAS conducted in the UK Biobank (UKB), the SAD variance component on PC2 was significantly large but only 16% of the size of the total variance of the PGS (*p =* 0.02, **Fig. S8B**). Using the most recent version (2022) of the GIANT height GWAS ^44^, we also observed large SAD variance on PC2 suggestive of stratification, this time with a size 0.10 of the size of the total PGS variance (*p =* 0.01; **Fig. 3B**). Furthermore, using the GIANT 2014 GWAS again, but constructing the PGS with a more stringent threshold for the significance of the index SNP, the SAD variance component on PC2 is only 3.3% of the total variance of the PGS (*p =* 0.4, **Fig. S9**; marginal GWAS *p*-value 10^−8^ instead of *p*-value 10^−5^ used in **Fig. 3A**).

The GIANT 2014 study ^42^ was, until recently, the largest GWAS of height, the complex trait most studied via GWAS, and so evidence of confounding sparked concern for complex trait research as a whole ^11,45^. Our partitionings of a PGS based on GIANT 2022 suggest the concern could extend to other recent GWAS meta-analyses as well. Some modern biobanks were designed to recruit large cohorts that are relatively ancestrally homogeneous with the intention of minimizing stratification ^32,46–48^. Nonetheless, the extent to which single biobank-based PGSs are affected by confounding remains an open question.

PGSUS can address this question. For concreteness, we first focus on the same set of UKB-based PGS built via clumping and thresholding at GWAS ascertainment *p*-value < 10^−5^ and their partitionings with respect to 1KG Europeans, as described in **The relative contribution of confounding to variation in a PGS**. In 12 of the 17 PGSs examined, we detected a significant (at level 0.05) SAD variance component in at least one of the top 6 PCs (**File S1**; *p* = 1.8 × 10^−4^ in a Bonferroni-corrected test of the hypothesis of no PC-specific SAD effects at play across all PGSs, see **Text S7**). For example, the SAD variance component on PC1 is approximately one third (33.9%) of the total variance in the PGS for forced vital capacity (FVC; *p =* 0.03; **Fig. 3C**). One possible contributor to PC-specific SAD variance is social stratification: lower FVC is associated with cigarette smoking^49^, which in turn is associated with educational attainment^50^. Among our GWAS-sample individuals, both pack-years of smoking and years of schooling are significantly associated with position on PC1 (**Fig. S10**). Other examples with significant SAD variance include a PGS for years of schooling with evidence of stratification along PC2 (*p =* 0.04; **Fig. 3D**) and a PGS for birth weight, a trait influenced by dynastic (maternal) effects linked to intrauterine environment^50–52^, (*p =* 0.0316; **Fig. 3E**). Results from other partitionings of a suite of PGS and prediction samples can be found in **Files S1-S8**.

### The relationship between confounding and PGS portability

PGS may be less applicable in samples that substantially differ, in some respect, from the GWAS samples in which they are estimated. This “portability” problem is often discussed in terms of variation in predictive accuracy across prediction samples that are more or less similar to the GWAS sample ^25,27,28,53,54^. (We return to this topic at the end of this section). Yet PGSUS can be informative about a broader sense of portability: regardless of its effect on prediction accuracy, the degree to which stratification explains variance in a PGS will depend on the specific prediction sample, possibly meriting additional caution in interpretation of the PGS with respect to that sample.

To illustrate this, we consider evidence of stratification we identified in two different prediction samples. We expected that, at least in some of these cases, the axes of ancestry with evidence of stratification will be aligned. Indeed, PCs with significantly large SAD variance were often correlated. For example, a UKB-based PGS of FVC had significant SAD variance along PC1 of 1KG Europeans and PC5 of 1KG; the SNP loadings for these two PCs have a Pearson correlation of |*r*| *=* 0.933 (*p <* 1 × 10^−250^; **Fig. S11A-C**). Similarly, the GIANT 2014-based PGS for height is stratified along PC2 in 1KG Europeans and PC1 of UKB NWB (Pearson correlation |*r*| *=* 0.096; *p* = 1 × 10^−258^; **Fig. S11D-F**).

In other cases, the relevant axes of stratification were not shared, and we detected significant PC-specific SAD variance only for some prediction samples. For example, a PGS constructed for body mass index (BMI) showed no evidence for PC-specific SAD variance when applied to 1KG Europeans (**Fig. 3F**). Yet in the full 1KG sample, we detected significant SAD variance on PC1, which largely tags differentiation between 1KG African and non-African individuals (*p =* 0.0484; **Fig. 3G**). We also detected significant SAD variance on PC4 in a prediction sample of ancient DNA from Eurasia, which partially distinguishes between Steppe and non-Steppe individuals (*p =* 0.040; **Figs. 3H, S12**).

The application of PGSs to ancient DNA samples is garnering interest, as several studies have used polygenic summaries based on a similar UKB GWAS to predict past phenotypes ^55–58^ or infer natural selection in recent European history ^8,59–63^. We examined partitionings of PGSs derived from UKB-based GWAS with respect to a sample of ancient West Eurasians who lived between 14,000 and 200 years ago (Allen Ancient DNA Resource ^37^). In addition to BMI (**Fig. 3H**), we found evidence of stratification among the top six PCs in PGSs for other traits reported to be under directional selection in ancient Eurasia during this time, including height, household income, skin color and hair color ^59,62,64^ (**File S1**). We note that the PCs with evidence of stratification are also significantly correlated with a number of sample variables, importantly date (as determined by conventional radiocarbon dating; **Fig. S13**). This raises the concern that confounding between major genome-wide changes in ancestry and UKB-based allelic effect estimates may spuriously contribute to polygenic signals of selection.

These examples help illustrate that PGSUS, as a diagnostic of confounding, can inform the application of a PGS to an intended prediction sample, but we now return to the traditional view of PGS portability in terms of prediction accuracy. We hypothesized that confounding can either aid or impede trait prediction, depending on whether SAD effects in the GWAS sample persist in the prediction sample. In a simulation study of stratification in the GWAS sample, we confirmed that prediction accuracy is higher when the confounding by population stratification in the prediction sample mimics that of the GWAS sample (**Text S14.10**; **Fig. S14**).

Finally, we turn to empirical case studies illustrating that confounded variance can indeed both increase and decrease trait prediction accuracy. We identified PGSs with significant PC-specific SAD variance components in the UKB NWB sample—a GIANT 2014 based PGS for height and a UKB-based PGS for household income (**Fig. 4A,C**). We randomly binned UKB NWB individuals such that bins represented variable degrees of variance in the PC scores among individuals with respect to the stratified PC (PC1 for GIANT height, PC3 for household income). We then predicted trait values and evaluated prediction accuracy (measured as the squared partial correlation between the PGS and trait within each bin) as a function of variance in the stratified PC. For GIANT height, prediction accuracy decreased with variation in the axis of stratification (Pearson *r =* 0.751, *p =* 1.3 × 10^−4^, **Fig. 4B**); whereas for household income, the trend was opposite (Pearson *r =* 0.715, *p =* 4.0 × 10^−4^; **Fig. 4D**; see **Figs. S15, S16** for similar trends obtained with other PGSs based on the same two GWASs).

**Figure 4.**
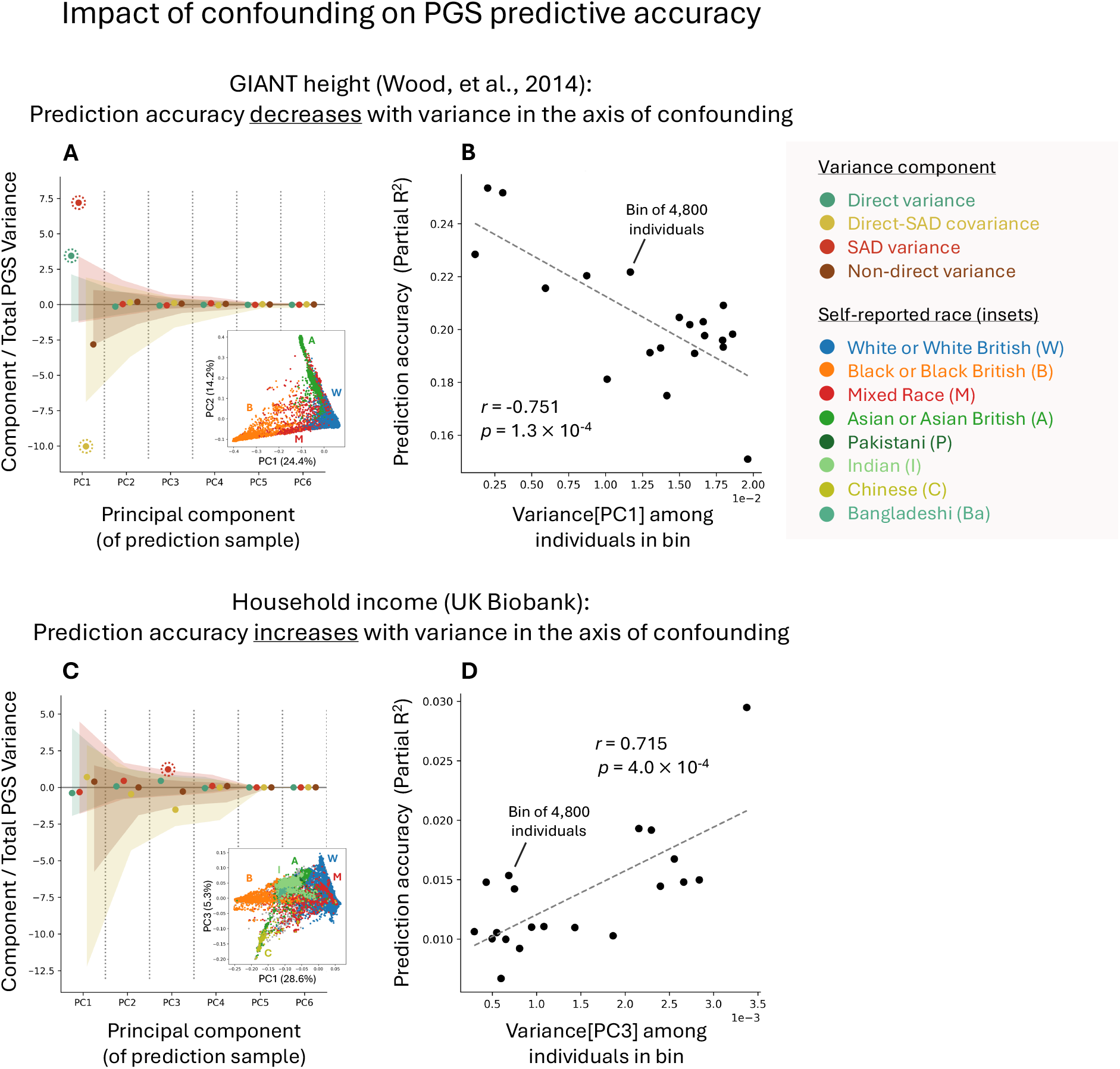
Confounding may either aid or impede PGS predictive accuracy. For two examples, we show variance components along the first six PCs divided by the total PGS variance in the prediction sample. Shaded regions show empirical permutation-based null non-rejection regions. Components deviating from this expectation are highlighted with dashed circles. Insets show the prediction-sample individual coordinates along the PCs on which we observed significantly large variance components. **(A)** A PGS for height based on GIANT 2014 with significant SAD variance along PC1 in a cohort of UK Biobank individuals not included in the White British subset used for GWAS (UKB NWB; n=96,493), which predominantly tags ancestry differences between self-identified Black individuals from the remainder of the UKB NWB sample. **(B)** Randomly binning individuals such that bins have variable degrees of variance in PC1 scores, we observed that prediction accuracy decreased with increasing variance in the axis of confounding. **(C)** A UKB-based PGS for household income with significant SAD variance along PC3 in the UKB NWB cohort, which in part tags ancestry differences between self-identified South and East Asians. **(D)** In contrast, our PGS of household income showed increased prediction accuracy as variance increased along the PC with significant SAD variance. Both PGSs are derived through clumping and thresholding with a GWAS ascertainment threshold *p <* 10^−3^.

### Isotropic inflation and non-direct variance

Often, standard-GWAS allelic effect estimates are larger than sib-GWAS effect estimates (after accounting for the degree of estimation error in each design) in a way that is not explained by any small set of principal components. Via a simulation study, we show that such “isotropic inflation”, or the systematic inflation of standard GWAS compared with corresponding sib-GWAS summary statistics, could be due to SAD effects (**Figs. S7, S17, S18**), as well as other factors, and importantly, ones related to the ascertainment of PGS index SNPs (**Fig. S19**). In the section **Estimation of the isotropic inflation factor** and **Text S10**, we expand this discussion.

This relationship is confirmed empirically. Across the set of UKB-based PGSs analyzed in this manuscript, nearly all PGSs show significant isotropic inflation (panels **(C)** in **Figs. S4-S6**), with more stringent GWAS ascertainment thresholds generally leading to more isotropic inflation (**Figs. S20-S22**).

We performed multiple simulations that suggest that factors that drive isotropic inflation tend to increase the non-direct variance component (**Fig. S23**; also see simulations confirming this as a plausible causal relationship, **Figs. S7, S17-S19, S24**). However, a theoretical model positing that all non-direct effects are isotropic provided a poor fit to these empirical data (**Fig. S25, Text S13**).

Given the many (non-exclusive) potential causes, we cannot make confident claims about the relative importance of drivers of isotropic inflation and non-direct variance. That said, our results do suggest that isotropic inflation, partly driven by index SNP ascertainment, is an important driver of non-direct variance.

## Conclusion

Through their uses in the clinic, research, and beyond, PGSs are sometimes assumed to represent direct, causal effects of an individual’s genotype on their phenotype. Most attempts to measure the extent to which this is true have been qualitative or heuristic in nature. The challenge of measuring the degree to which a PGS (and its variation in a prediction sample) instead reflects confounding factors—including environmentally and socially mediated factors—is critical to its interpretation and application. PGSUS is a proposed step toward addressing this challenge.

## Acknowledgements

We thank Graham Coop for his major contributions in conceptualizing PGSUS and guidance throughout its development. We thank Ipsita Agarwal, Yi Ding, Jonathan Pritchard, Molly Przeworski, Jeff Spence, Elliot Tucker-Drob, and members of the Harpak Lab for feedback on the manuscript. The work was funded by NIH grant RF1AG073593 to Elliot Tucker-Drob, NIH grants F32GM130050 and R35GM137758 to M.D.E., and NIH grant R35GM151108, a Pew Biomedical Scholarship, and a fellowship from the Simons Foundation’s Society of Fellows (#633313) to A.H. This study was conducted using the UK Biobank resource under application 92741, as approved by the University of Texas at Austin institutional review board (study protocol 00003287). The authors acknowledge the Texas Advanced Computing Center (TACC) at The University of Texas at Austin for providing computational resources that have contributed to the research results reported within this paper.

## Methods

### PGSUS-based parameter estimation

#### Overview of parameter estimation

To generate estimates underlying the output of PGSUS (**Fig. 1**), we perform two partitionings sequentially as outlined in **Results and Discussion**. For the first partitioning, we set 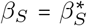 (i.e., setting *α* ≔ 1 in **Eq. 3**) estimating the non-direct variance component as

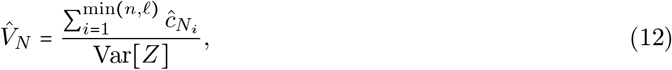

where

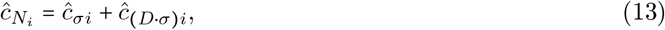

where the two terms in this sum are defined in **Eq. 18** and **Eq. 22**. The isotropic inflation factor is then estimated as described in the section **Estimation of the isotropic inflation factor** below. This estimate, 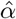, is the one we report. We then adjust for isotropic inflation by multiplying sib-based summary statistics—allelic effect estimates and standard errors—by 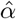 (**Eq. 11**) such that they are on the same scale as standard-GWAS summary statistics. These rescaled sib-GWAS summary statistics are then used as input for the second partitioning, yielding the estimates of (isotropic inflation-adjusted) PC-specific variance components we report.

#### Estimating PC-specific variance components given the isotropic inflation factor *α*

We describe a moment-based approach to estimating the variance components in **Eqs. 6-8** that applies if the isotropic inflation factor *α* is known. Our approach assumes that the difference between standard-GWAS-based and sib-GWAS-based allelic effect estimates, other than differences in estimation noise, is that the sib-based estimates are free of SAD effects (**Eqs. 2-3**). We note that, even in the absence of SAD effects, the estimated effects are expected to differ in the two study designs ^14^—but the difference is often small (**Text S2**). Applying **Eq. 1**, we then reason that the projection of the differences between standard-GWAS and sib-GWAS allelic effect estimates on the principal components will provide information about the proportion of variance in a PGS attributable to SAD.

Applying the model in **Eqs. 2-3** to index SNPs *j* ∈ {1, …, *ℓ*}, the variance component of the difference between standard-GWAS and sib-GWAS allelic effect estimates that is due to the *i*th principal component is

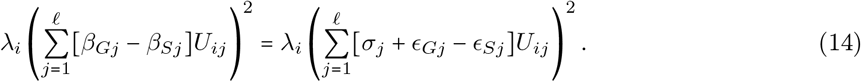

To derive the expectation of the expression in **Eq. 14**, we assume that the *λ* (eigenvalue), *σ* (SAD effect), and *U* (locus loadings on the eigenvector) terms are fixed and that the measurement error terms all have expectation 0 (i.e. 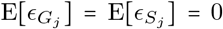 for all *j*). This assumption may be problematic for some traits as applied to some PCs when SNPs are ascertained for inclusion in the PGS according to *p*-value, which we discuss further in **Text S15.2**. We also assume that the error terms from the standard GWAS and the sib-GWAS are uncorrelated with each other (i.e. 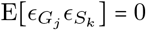 for all *j* and *k*, including *j = k*). The assumption that the measurement errors from the standard GWAS and sib-GWAS studies are uncorrelated might be problematic if, e.g., the sib-GWAS is performed in a subset of the standard-GWAS sample. Nonetheless, we adopt it here. Under these assumptions,

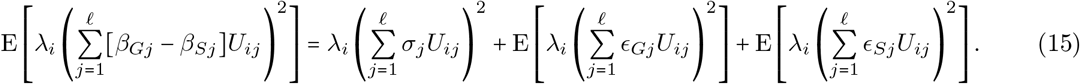

The first term on the right is 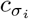, the variance component of the PGS attributable to SAD effects along principal component *i*, and one of our main estimands of interest. The latter two terms are variance components attributable to measurement errors in the standard-GWAS allelic effect estimates and the sib-GWAS allelic effect estimates, respectively. If we can assume that measurement errors at distinct loci are uncorrelated, or, slightly less restrictively, that

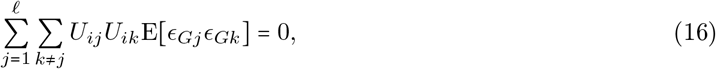

then the expected variance component due to GWAS errors is

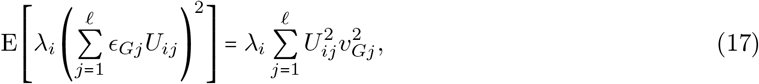

where 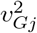 is the variance of the measurement error of the standard-GWAS allelic effect estimate at locus *j*. Making an analogous assumption for the measurement errors in the sib-GWAS allelic effect estimates, we arrive at an estimator for the variance component of the PGS due to SAD effects along principal component 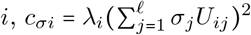,

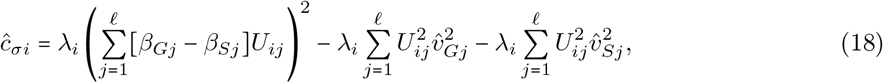

where 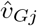 is an estimated standard error of the standard-GWAS allelic effect estimate and 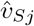 is an estimated standard error of the sib-GWAS allelic effect estimate at locus *j*.

Similarly, we can estimate the variance component of the PGS attributable to direct effects along the *i*th principal component of the genotype matrix of the prediction sample, 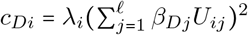, by starting with the projection of the sib-GWAS allelic effect estimates on principal component *i*. Under the same assumptions about the measurement errors in the sib-GWAS allelic effect estimates used immediately above (i.e. that the measurement errors have expectation 0 and 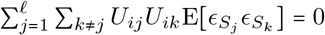), we have

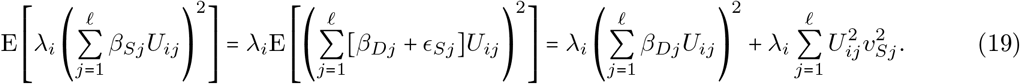

Thus, to estimate *c*_*Di*_, we use

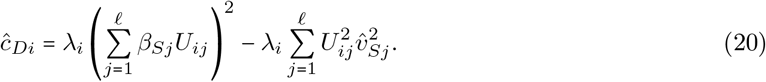

Finally, adapting **Eq. 5**, the variance along the *i*th principal component of the GWAS-based PGS can be written as

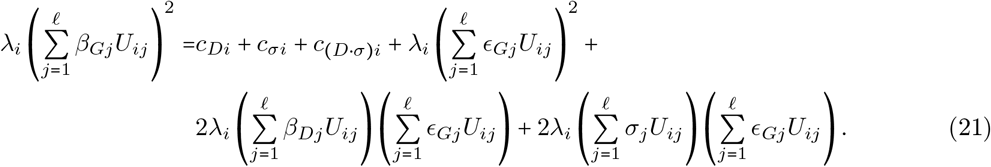

The final two terms have expectation zero because of the assumption that 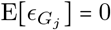 for all *j* and because all the other variables in the final two terms are treated as fixed. We thus estimate *c*_(*Dσ*) *i*_ by plugging in estimators for the other variance components and rearranging, giving

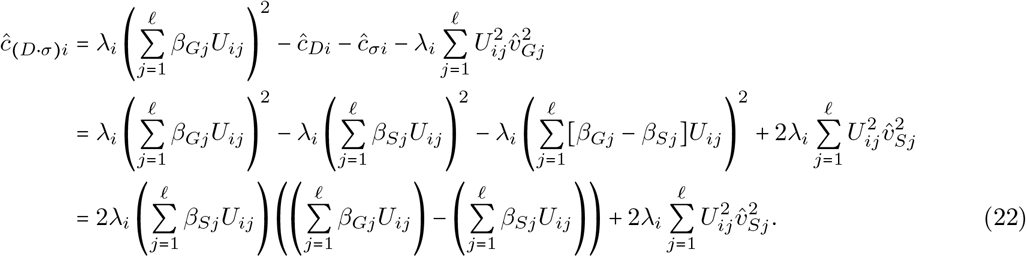

As mentioned previously, the estimands *c*_*Di*_ and *c*_*σi*_ are constrained to be non-negative, whereas *c* _*Dσi*_ may take positive or negative values. However, the estimators *ĉ*_*Di*_, *ĉ*_*σi*_, and *ĉ* _(*Dσ*) *i*_ presented here can all take positive or negative values. In **Text S3**, we describe our empirical permutation strategy for testing hypotheses about the variance components.

#### Estimation of the isotropic inflation factor

The presence of isotropic inflation, namely *α* ≠ 1, renders the approach above incomplete. With *α* as a free parameter, there are four estimands per principal component (*α* and the three variance components) and only three pieces of information per principal component (the projections of the sib-GWAS allelic effect estimates, standard-GWAS allelic effect estimates, and their difference on the principal component). However, *α* should be shared across all principal components, which gives other possibilities for estimation.

Because we are mostly interested in principal component-specific variance components for the “top” principal components—i.e. the principal components associated with the largest eigenvalues—and given evidence that other PCs often do not capture meaningful axes of population structure ^10^—we adopt a strategy in which we use the principal components with smaller eigenvalues to estimate *α*, meaning that the *σ* terms in the top principal components can be interpreted as departures from the relationship between sib-GWAS and standard-GWAS allelic effect estimates on the other components.

In particular, we assume that on the (*k +* 1)^*th*^ to final principal components (where PCs are ranked by eigenvalue), the PC-specific SAD effects *σ* are 0, yielding

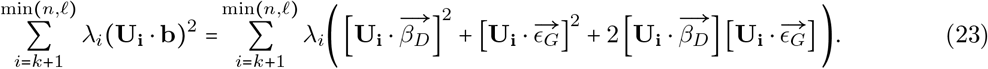

As before, the third bracketed term inside the parentheses is assumed to have expectation 0; the second one has an expectation that we can estimate using the standard error of the standard-GWAS allelic effect estimates. Analogously to the estimator of *c*_*Di*_ in **Eq. 20**, we can get an alternative estimate of the variance due to direct effects along PC *i* > *k*,

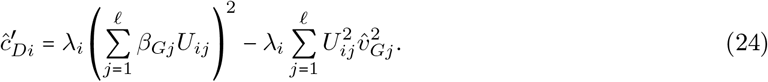

Setting the right hand side equal to the other estimator of *c*_*Di*_ (**Eq. 20**) gives

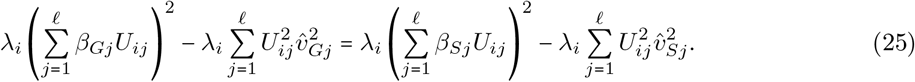

Since, instead of observing *β*_*Si*_, we observe 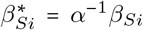, and obtain estimated squared standard errors for the observed sib-GWAS allelic effect estimates 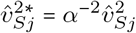, then we can write

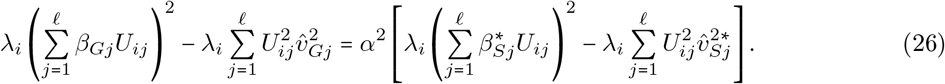

or

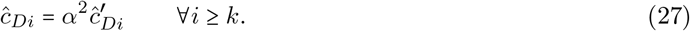

This expression suggests that we might estimate *α* as the square root of the coefficient relating 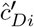 to 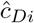 in a no-intercept regression across PCs *i* ≥ *k*. Note that both the estimators of *c*_*Di*_ appearing in **Eq. 27** have measurement error as given in **Eq. 42** for *ĉ*_*Di*_ and, equivalently, by

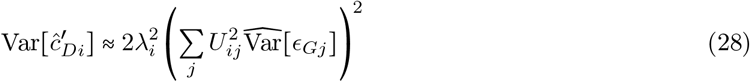

For 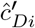, where 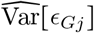 is the squared standard error estimate for the allelic effect at locus *j*.

We estimate *α*^2^ by setting *k* = 101 and using Deming Regression, an errors-in-variables method which accounts for uncertainty in each of the two axes that are considered ^65^. We obtain an estimate of the standard error of the isotropic inflation factor by performing 100 bootstraps of the paired estimates for each PC used in the empirical Deming regression and taking the standard error of the effect size estimates. We use the isotropic inflation factor to adjust for the systematic difference in standard and sib-GWAS statistics by multiplying the sib-GWAS statistics by 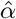 for a given PGS. We use this approach so that the total PGS variance before and after correction remains unchanged, since the standard-GWAS summary statistics remain unaltered.

### Large direct variance components as evidence of natural selection

Direct-effect variance components that deviate from the null non-rejection regions (green shaded regions in **Fig. 3**) are potentially suggestive of selection, as these regions include the center of the distribution of the component under a model in which trait-associated alleles (namely, PGS index SNPs) evolve neutrally. We observed several cases of significant direct variance components among top PCs in the PGSs we partitioned (**Files S1-S8**). For example, the significant direct variance components for PC2 and PC3 of 1KG Europeans in a PGS for waist circumference (**Fig. S26A,B**) are consistent with previous reports on selection for pelvic morphology to minimize obstetric obstruction ^66,67^ and its impact on ilium morphological development^68^, and hip-width proportions ^69^. In many instances, this signal was accompanied by significantly large SAD and/or non-direct variance along the same PC (**Figs. S26, S27**).

Such an observation might be expected if both selection and stratification are at play, or if selection acts on both direct and dynastic effects. However, a simulation study we performed suggested that such co-occurrence of large SAD and direct components on a PC may arise in the presence of stratification along that PC, even if there is no selection (**Text S15.1, Fig. S28**). (See further discussion in **Text S6** and Zaidi and Mathieson ^70^ and Blanc and Berg ^71^ for similar observations.) We therefore cannot interpret a large direct variance component as necessarily pointing to evidence of natural selection in these cases.

### Assessing predictive accuracy in confounded PGSs

In the section **The relationship between confounding and PGS portability**, we investigate the relationship between variance in a genetic ancestry axis along which a PGS is confounded and prediction accuracy. The prediction samples in this analysis are subsets of non-White British individuals in the UK Biobank—not included in either standard or sib-GWAS we performed in other analyses, who were further filtered using the same procedure as those included in standard-GWAS samples based on UKB (UKB NWB; see **Data**). We applied PGSUS to each of the 17 UKB-based PGSs as well as for the GIANT 2014 height PGS. We then identified two examples which showed significant evidence of PC-specific SAD variance along a top PC: along PC1 for height (GIANT 2014) and PC3 for household income (UKB WB-based GWAS).

We partitioned the UKB NWB cohort into 20 non-overlapping bins of 4,800 individuals, each with increasing variance along the PC of interest. To create a bin of individuals, we sampled 480 individuals from the most extreme scores along that PC and further sampled the remaining 4,320 individuals randomly. We removed the binned individuals from the sample pool and repeated this sampling procedure to generate 20 bins. We computed PGSs for all individuals using *PLINK 2*.*0* ‘s `--score` function.

We used partial *R*^2^ between the PGS and the trait value to evaluate predictive accuracy. For each bin, following Wang et al. ^28^, we first regressed the trait value on the covariates age, age^2^, sex, age × sex, and age^2^ × sex (phenotype ∼ covariates, i.e., the reduced model). Then we regressed the trait value on the PGS and covariates (phenotype ~ covariates + PGS, i.e., the full model). We then calculated partial *R*^2^ using

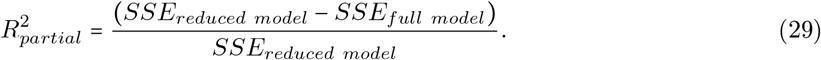

### Data

#### UK Biobank

The UK Biobank is a large database with extensive phenotypic and genetic information for over half a million individuals in the UK who were between 40 and 69 years of age at the time of recruitment^32^. The individuals included in this study were those who passed checks excluding individuals who: were identified by the UK Biobank to have sex chromosome aneuploidy, self-reported sex differing from sex determined by genotype data, and participants who withdrew from the study as of the last withdrawal notification delivery (April 24, 2024). We excluded individuals in the standard-GWAS cohort who were third-degree relatives or closer as identified by the UK Biobank in data field 22021. We further limited our sample to those individuals who were classified as “White British” (UKB WB; data field 22006) by the UKB, corresponding to individuals that self-identified as both White and British and were also found to be tightly clustered in the genetic principal component space ^32,72^. Finally, individuals who we identified as having full siblings in the cohort were removed to ensure no sample overlap between the standard-GWAS and sib-GWAS cohort. For each phenotype, we also omitted individuals who had missing data for the phenotype of interest, resulting in between 96,942 and 321,718 individuals across the 17 continuous traits we analyzed in the standard GWAS. In the sibling cohort, we only included pairs of siblings who both had phenotypic measurements, resulting in between 2,163 and 17,328 sibling pairs across phenotypes. A full account of the traits studied, corresponding UK Biobank data field identifiers, and GWAS sample sizes can be found in **Table S1**.

In our analysis of the relationship between prediction accuracy and confounding (**Fig. 4**, also see the section **The relationship between confounding and PGS portability**), we also refer to a UKB non-White British sample (UKB NWB). These are 96,493 individuals who meet all aforementioned filtering criteria except the WB classification. In our analysis of the utility of GWAS adjustments for population structure (**Fig. 2C, Text S9**), we also considered a random subset of 100,000 individuals from the full post-filtering sample, i.e. the union of UKB WB and UKB NWB.

A few specific phenotypes are worth noting for the transformations and filtering steps. We calculated the phenotype ‘years of schooling’ by converting the maximum educational attainment of the participants to years following Okbay et al. ^73^. For the ‘hand grip strength’ phenotype, we took the average of the measurement across both hands. Finally, for diastolic blood pressure levels, we adjusted the measurements of individuals who were taking blood pressure medication upward by 10 mmHg ^74^.

From the relatedness information provided by the UK Biobank in Resource 531 and data field 22021, we identified 17,353 White British full-sibling pairs using the following protocol. Groups of siblings were identified as those pairs of individuals who had a kinship coefficient between 0.1768 and 0.3536, as well as an IBS0 value greater than 0.0012, as estimated using the software KING ^75^. From each of the resulting family groups we then selected two individuals, resulting in the aforementioned 17,353 sibling pairs where every individual is only a part of a single sibling pair. Each individual identified as a part of a sibling pair was omitted from the standard-GWAS analysis of each trait.

We identified a set of 9,607,691 genetic variants that passed the following quality control filters using the *PLINK 2* software ^76^ in the standard cohort to include in both the standard GWASs and sib-GWASs for all traits following the protocol specified in Zhu et al. ^77^. Using the set of genotyped and imputed variants, we first filtered out all variants with an INFO score lower than 0.8. We then removed all SNPs with genotype missingness greater than 0.05 using the *PLINK 2* flag `--geno 0.05` and SNPs with an alternate allele frequency lower than 0.1%, *PLINK 2* flag `--maf 0.001`. Finally, we removed all variants that were not biallelic SNPs (`--snps-only`) and all variants which are far from Hardy-Weinberg equilibrium (`--hwe 1e-10`).

#### 1000 Genomes

In order to estimate the SAD variance in the 17 complex traits of interest in the UK Biobank, we downloaded the whole genome sequences from phase three of the 1000 Genomes (1KG) project at https://ftp.1000genomes.ebi.ac.uk/vol1/ftp/release/20130502/. From the initial set of roughly 80 million SNPs, we extracted the set of overlapping SNPs with a minor allele frequency greater than 1% in the 1000 Genomes and the SNPs passing QC steps in the UK Biobank. This resulted in 6,131,234 SNPs to be included in our analysis. We used these variants for all future analyses of both the entire 1KG cohort (*N =* 2, 504) and those individuals in the European superpopulation (*N =* 503, referenced as the 1KG Europeans).

### Software availability

All of the summary statistics generated in this study are available on the Harpak Lab website’s data tab https://www.harpaklab.com/data. The software for SAD variance estimation and a working example are available at https://github.com/harpak-lab/PGSUS. GIANT summary statistics were downloaded from https://portals.broadinstitute.org/collaboration/giant/index.php/GIANT_consortium_data_files. Finally, AADR data were downloaded from https://reich.hms.harvard.edu/datasets. See further details on external data processing in **Text S8**.

