## Supplementary Materials for "A Litmus Test for Confounding in Polygenic Scores"

**Supplemental File Descriptions** **92**

### Supplemental Text

#### S1 Mathematical details for the PCA-based variance decomposition

In the main text, we give an expression (**Eq. 1**) for a decomposition of the polygenic score (PGS) variance into terms attributable to distinct principal components, written in terms of locus-level additive allelic effect estimates and locus-level loadings on principal components. In this section, we justify this decomposition.

We begin with genotype data represented as a matrix $\mathbf{X}$. $\mathbf{X}$ has $n$ rows, one for each individual, and $\ell$ columns, one for each genetic locus; entry $ij$ contains the number of copies of a non-reference allele carried by individual $i$ at locus $j$. For some trait, we have a column vector $\mathbf{b}$ of length $\ell$ containing allelic effect estimates of the non-reference allele, which is used to compute a PGS $\mathbf{Z}$, a vector, as $\mathbf{Z} = \mathbf{X}\mathbf{b}$.

To perform principal components analysis (PCA) on $\mathbf{X}$, we first compute a covariance matrix $\mathbf{V_X} = \frac{1}{n-1}\mathbf{X^T C_n X}$, where $\mathbf{C_n}$ is the centering matrix of dimension $n$. Eigendecomposition then gives the diagonalization

$$\mathbf{V_X} = \frac{1}{n-1}\mathbf{X^T C_n X} = \mathbf{U \Lambda U^{-1}} = \mathbf{U \Lambda U^T}, \tag{24}$$

where $\mathbf{U}$ is an orthogonal $\ell \times \min(n,\ell)$ matrix of eigenvectors (orthogonality follows from the symmetry of the covariance matrix), and $\mathbf{\Lambda}$ is a diagonal $\min(n,\ell) \times \min(n,\ell)$ matrix of eigenvalues. (The last step follows because $\mathbf{U}$ is an orthogonal matrix and so $\mathbf{U^{-1}} = \mathbf{U^T}$.) The diagonal matrix of eigenvalues $\mathbf{\Lambda}$ is the covariance matrix of the data $\mathbf{X}$ expressed in principal-component coordinates, the axes of which are delineated by the columns of $\mathbf{U}$.

The variance of a random draw $Z$ from the $n$ PGS values in the vector $\mathbf{Z}$ can be computed as

$$\mathrm{Var}[Z] = \frac{1}{n-1}\mathbf{Z^T C_n Z}.$$

By the definition of $Z$,

$$\mathrm{Var}[Z] = \frac{1}{n-1}\mathbf{Z^T C_n Z} = \frac{1}{n-1}\mathbf{b^T X^T C_n X b} = \mathbf{b^T V_X b} = \mathbf{b^T U \Lambda U^T b}.$$

Because the columns of $\mathbf{U}$ are orthogonal, the last statement in the previous equation is equivalent to

$$\mathrm{Var}[Z] = \sum_{i=1}^{\min(n,\ell)} \lambda_i (\mathbf{U_i} \cdot \mathbf{b})^2, \tag{25}$$

where $\lambda_i$ is the $i$th eigenvalue from $\mathbf{\Lambda}$, $\mathbf{U_i}$ is the $i$th column of $\mathbf{U}$, and $\cdot$ represents the dot product. This is the source of **Eq. 1**. In words, this is a way of partitioning the variance of the PGS into components attributable to each principal component of the genotype matrix.

The variance partitioning in **Eq. 25** is equivalent to the one that would be obtained by regressing individual-level PGS on individual-level principal component scores. That is, for each principal component $i$,

$$r^2(\mathbf{XU_i}, \mathbf{Z})\mathrm{Var}[Z] = \lambda_i(\mathbf{U_i} \cdot \mathbf{b})^2, \tag{26}$$

where $r^2(\mathbf{XU_i}, \mathbf{Z})$, is the squared sample correlation coefficient between the individual-level scores on principal component $i$ and the individual-level PGS. To show this, we start by substituting in the definition of the squared correlation coefficient in terms of sample variances and covariances, centering the variables on the right using the centering matrix $\mathbf{C_n}$ (which does not affect the sample variances or covariances),

$$r^2(\mathbf{XU_i}, \mathbf{Z})\mathrm{Var}[Z] = \frac{\mathrm{Cov}(\mathbf{C_nXU_i}, \mathbf{C_nXb})^2}{\mathrm{Var}[\mathbf{C_nXU_i}]}. \tag{27}$$

The denominator $\mathrm{Var}[\mathbf{C_nXU_i}]$ is equal to the $i$th eigenvalue by the properties of the eigendecomposition. Making this substitution and expressing the numerator in terms of the dot product of the two vectors gives

$$r^2(\mathbf{XU_i}, \mathbf{Z})\mathrm{Var}[\mathbf{Xb}] = \frac{\left(\frac{1}{n-1}\right)^2 (\mathbf{C_nXU_i} \cdot \mathbf{C_nXb})^2}{\lambda_i}. \tag{28}$$

The dot product on the right side of **Eq. 28** can also be expressed as $\mathbf{C_nXU_i} \cdot \mathbf{C_nXb} = \mathbf{U_i^TX^TC_nXb}$, where the (symmetric) centering matrix $\mathbf{C_n}$ appears only once because centering is a projection, and so $\mathbf{C_n^TC_n} = \mathbf{C_n}$. Making this substitution and using **Eq. 24** gives

$$r^2(\mathbf{XU_i}, \mathbf{Z})\mathrm{Var}[Z] = \frac{(\mathbf{U_i^TU\Lambda U^Tb})^2}{\lambda_i} = \frac{(\lambda_\mathbf{i}\mathbf{U_i} \cdot \mathbf{b})^2}{\lambda_i} = \lambda_i(\mathbf{U_i} \cdot \mathbf{b})^2, \tag{29}$$

which is what we sought to prove. The penultimate step holds because the $\mathbf{U}$ matrix is orthonormal, so $\mathbf{U_i} \cdot \mathbf{U_i} = 1$ and $\mathbf{U_i} \cdot \mathbf{U_j} = 0$ for all $j \neq i$.

#### S2 The expectation of standard-GWAS and sib-GWAS allelic effect estimates

In this section, we study the expected values of standard-GWAS and sib-GWAS allelic effect estimators in a scenario with only direct effects and in the absence of drift, selection, and mutation acting over one generation. We derive the expectations of these allelic effects at a tag SNP that is genotyped in both the standard GWAS and sib-GWAS, tagging a causal SNP that is not genotyped. We show these expectations are not quite equal because of differences in the effect of linkage disequilibrium on the standard and sibling-based GWASs. This is contrary to what we assume in the main text, in which we treat the direct effects in sib-GWAS and standard GWAS as equal. However, we also show that differences are small if the recombination rate between the tag and causal SNPs is small. A more general and complete approach to similar questions appears in Veller & Coop[1]; we include the derivation here for completeness.

We consider an autosomal tag SNP with alleles A and a and an autosomal causal SNP with alleles B and b. The A and B alleles are in LD with strength $D_p$ in the parents of the generation we will study, and the recombination fraction between the loci is $\rho$ in both sexes. Each copy of the B allele increases the phenotype of interest, $Y$, by an amount $\gamma$. There are no SAD effects. The generation studied by a GWAS results from a generation of random mating in the parental generation. As a result, the LD between the tag SNP and causal SNP in the generation we study is $D = D_p(1-\rho)$, assuming that recombination is the only force affecting LD[2].

We model a standard GWAS with no adjustment for covariates. The association between the phenotype $Y$ and the tag SNP is due entirely to the causal SNP, and so the correlation between $Y$ and the causal SNP is

$$\mathrm{Cor}[Y, A_C] = \mathrm{Cor}[Y, B_C]\mathrm{Cor}[B_C, A_C] = \frac{\gamma D}{\sqrt{\mathrm{Var}[Y]\mathrm{Var}[A_C]}},$$

where $A_C$ and $B_C$ represent allele counts for the A and B alleles, respectively, and

$$\mathrm{Cov}[Y, A_C] = \gamma D.$$

In a simple linear regression of individual phenotype values on the number of $A$ alleles, the expected value of the additive allelic effect estimate will be

$$\mathrm{E}[\beta_G] = \frac{\gamma D}{p_\mathrm{A}(1-p_\mathrm{A})} = \frac{\gamma D_p(1-\rho)}{p_\mathrm{A}(1-p_\mathrm{A})}, \tag{30}$$

where $p_\mathrm{A}$ is the frequency of the A allele, since the expectation of the simple linear regression estimator is the covariance of the dependent and independent variables divided by the variance of the independent variable.

In the sib-GWAS, the estimator is equivalent to a linear regression with no intercept of the within-sibling-pair differences in phenotype regressed on the within-sibling-pair differences in genotype. Define $A_d$ the difference between siblings in the count of tag allele A (say sibling 1 minus sibling 2, where sibling 1 is chosen randomly), $B_d$ the difference between siblings in the count of causal allele B, and $Y_d$ the difference between siblings in the phenotype. To proceed, we need the covariance of $A_d$ and $B_d$, as well as their variances. Because sibling 1 is chosen at random, $A_d$, $B_d$, and $Y_d$ are all symmetric with expectation 0.

To compute the covariance of $A_d$ and $B_d$, consider the contribution to $A_d$ and $B_d$ of the haplotypes transmitted to each sibling by the biological mother, labeled $A_d(m)$ and $B_d(m)$. In principle, each sibling can receive one of four different haplotypes from the mother: AB, Ab, aB, and ab, giving 16 combinations. However, only four of these combinations give a nonzero value of $A_d(m) * B_d(m)$ and thus need to be considered when computing $\text{Cov}[A_d(m), B_d(m)] = \text{E}[A_d(m) * B_d(m)]$. When one sibling inherits AB and the other inherits ab, then $A_d(m) * B_d(m) = 1$. When one inherits Ab and the other inherits aB, then $A_d(m) * B_d(m) = -1$. Thus, we need to compute the probabilities of these two events.

There are two ways that one sibling can inherit AB and the other can inherit ab. First, the mother carries haplotypes AB and ab, and she passes unrecombined haplotypes to her children. This occurs with probability $p_{\text{AB}}p_{\text{ab}}(1-\rho)^2$, where $p_{\text{AB}}$ is the frequency of haplotype AB and $p_{\text{ab}}$ is the frequency of haplotype ab. The probability that the mother carries these two haplotypes is $2p_{\text{AB}}p_{\text{ab}}$, but the two cancels with the probability 1/2 that the two siblings inherit different haplotypes from their mother. The second way in which siblings with AB and ab haplotypes might be produced is if the mother carries haplotypes Ab and aB, and she passes a recombinant AB to one child and a recombinant ab to the other child. By a similar calculation, this occurs with probability $p_{\text{Ab}}p_{\text{aB}}\rho^2$. Thus, the probability that one sibling inherits AB and the other inherits ab is the sum

$$\text{P}(A_d(m) * B_d(m) = 1) = p_{\text{AB}}p_{\text{ab}}(1-\rho)^2 + p_{\text{Ab}}p_{\text{aB}}\rho^2. \tag{31}$$

The probability that one sibling inherits Ab and the other inherits aB, in which case $A_d(m) * B_d(m) = -1$, is computed similarly. The mother can either pass on unrecombined haplotypes Ab and aB, or she can carry haplotypes AB and ab and pass different recombinant haplotypes to each child. The probability that either one of these events happens is

$$\text{P}(A_d(m) * B_d(m) = -1) = p_{\text{Ab}}p_{\text{aB}}(1-\rho)^2 + p_{\text{AB}}p_{\text{ab}}\rho^2. \tag{32}$$

The covariance of $A_d(m)$ and $B_d(m)$ is the difference of the expressions in **Eqs. 31** and **32**, or

$$\begin{aligned}\text{Cov}[A_d(m), B_d(m)] &= p_{\text{AB}}p_{\text{ab}}(1-\rho)^2 + p_{\text{Ab}}p_{\text{aB}}\rho^2 - p_{\text{Ab}}p_{\text{aB}}(1-\rho)^2 - p_{\text{AB}}p_{\text{ab}}\rho^2 \\ &= [(1-\rho)^2 - \rho^2](p_{\text{AB}}p_{\text{ab}} - p_{\text{Ab}}p_{\text{aB}}) \\ &= D_p[(1-\rho)^2 - \rho^2] = D_p(1-2\rho). \end{aligned} \tag{33}$$

The simplification $p_{\text{AB}}p_{\text{ab}} - p_{\text{Ab}}p_{\text{aB}} = D_p$ comes from a standard identity for the linkage disequilibrium covariance of allele counts statistic[2]. Because the recombination rates are assumed to be the same in both sexes, the analogous covariance due to the father's contribution will be identical, and the father's and mother's contributions will be independent. Thus, the desired covariance of the within-sibling differences in genotype at the tag site and the causal site is

$$\mathrm{Cov}[A_d, B_d] = 2D_p(1-2\rho). \tag{34}$$

The variances of $A_d$ and $B_d$ are

$$\begin{aligned}\mathrm{Var}[A_d] &= 2p_{\mathrm{A}}(1-p_{\mathrm{A}})\\ \mathrm{Var}[B_d] &= 2p_{\mathrm{B}}(1-p_{\mathrm{B}}).\end{aligned} \tag{35}$$

For intuition, note that the variance of the signed difference in allele count between two unrelated people would be twice the variance of an individual's allele count, or $4p_{\mathrm{A}}(1-p_{\mathrm{A}})$. These variances are half as great because they are sibling pairs who may share 0, 1, or 2 alleles identical by descent.

**Eqs. 34** and **35**, along with an argument parallel to the one justifying **Eq. 30**, give the expectation of the sib-GWAS estimator. It is

$$\mathrm{E}[\beta_S] = \frac{\gamma D_p(1-2\rho)}{p_{\mathrm{A}}(1-p_{\mathrm{A}})}. \tag{36}$$

The sibling-based expectation in **Eq. 36** differs from the standard-GWAS expectation in **Eq. 30** only in that the parental linkage disequilibrium is multiplied by one minus twice the recombination fraction instead of one minus the recombination fraction. If the recombination fraction between the tag site and a causal site is very small, then the causal site's contribution to the phenotype association at the tag site will be similar when the GWAS is standard or sibling-based. However, if the recombination fraction is large, then the causal site's effects on the standard-GWAS and sib-GWAS allelic effect estimates might be expected to differ substantially. In practice, we typically expect causal alleles that have large recombination fractions with the focal SNP to be in very low or no LD with the focal SNP. Causal alleles in appreciable LD with the focal SNP will often be those with very small recombination fractions, held at relatively high LD in an equilibrium between genetic drift and recombination.

#### S3 Permutation tests for variance components

We use permutation to assess whether variance components corresponding to a given principal component differ significantly from a null prediction. If the loci were independent, we could permute the allelic effect estimate at each locus independently to generate a null of no association between allelic effects and loadings. However, because of linkage disequilibrium, loci cannot be considered independent. As such, we randomly permute the signs of the allelic effect estimates for

1,700 approximately independent linkage blocks[3]. That is, in a given permutation, for each block, all the allelic effect estimates are multiplied by either 1 or -1 with probability 1/2 each. For both effects and their principal component loading, such a strategy preserves any correlations between loci in the same block[4,5]. This strategy may be conservative if there are weakly correlated index SNPs within blocks.

In practice, we perform permutation tests using the estimated variance components as test statistics, applying the same blockwise permutation to both standard-GWAS and sib-GWAS allelic effect estimates. If sib-GWAS allelic effect estimates are associated with loadings on principal component $i$, then the estimated variance component for the direct effect $\hat{c}_{Di}$ (**Eq. 20**)—which depends principally on the covariance between PC loadings and sib-GWAS allelic effect estimates—will tend to be large compared with their permutation distribution. The null distribution formed is one in which the absolute values of sibling-based allelic effect estimates are preserved, but their sign is uniformly random with respect to principal component loading. Similarly, $\hat{c}_{\sigma i}$ (**Eq. 18**) depends mainly on the difference between standard-GWAS and sib-GWAS allelic effect estimates, and as such it is sensitive to, for example, stratification along principal component $i$. The null distribution generated by permutation is one in which, in each genomic block, the sign of the difference between the standard-GWAS and sib-GWAS allelic effect is randomized. In our significance testing for the PC-specific variance components, the direct and SAD variance components are tested with one-sided tests, since these components may only take non-negative values. $p$-values for direct-SAD covariance and non-direct components are evaluated using two-sided tests, since these components may take positive or negative values.

Whereas the SAD component is related to the projection of the difference between the standard-GWAS and sib-GWAS allelic effects on principal component $i$, the non-direct variance component, $\hat{c}_{N_i}$ (**Eq. 9**), is related to the difference in the projections of the standard-GWAS and sib-GWAS allelic effects on the principal component. We also perform permutation tests on $\hat{c}_{(D\cdot\sigma)i}$ (**Eq. 22**), though they are perhaps of less interest.

#### S4 Adaptive permutation procedure for calculating variance components

The permutation procedure described in the preceding section (**Permutation tests for variance components**) presumes a set number of permutations (default of 1,000) are performed for each variance component along each top PC of interest. However, for PC-specific variance components at (or close to) a significance level of 0.001, PGSUS users may want increased resolution on significance. Conversely, a smaller number of permutations may suffice once non-significance is clear for a given test. We therefore describe here an adaptive permutation procedure modeled after Che et al.[6].

The adaptive permutation procedure requires four user-specified quantities: a $p$-value significance threshold $\eta$, a relative precision parameter $c$, a minimal number of observed successes $R$, and a maximal number of permutation replicates $B$. By default, we set these parameters to $\eta = 0.05$, $c = 0.1$, $R = \frac{1}{c^2} = 100$, and $B = \frac{1-\eta}{c^2 \cdot \eta} = 1900$. With these default values, the standard error of the estimated $p$-value near $\eta = 0.05$ is approximately $c \cdot \eta = 0.005$.

The algorithm then proceeds as follows. As described in **Permutation tests for variance components**, we use the estimated variance components as test statistics, denoted as $T_{obs}$. In each permutation replicate $b$, the same test statistic is computed from the permuted allelic effects, giving $T_{(b)}$. We call a permutation replicate a "success" (following the convention in Che et al.[6]) if $T_{(b)}$ is at least as extreme as $T_{obs}$. After each permutation replicate, the algorithm updates the total number of permutations performed $b$ and the number of observed successes $r$. The procedure stops when either $R$ successes have been observed, or the maximum number of $B$ permutations has been reached. If $R$ successes are observed before reaching $B$ permutations, the test statistic is unlikely to be significant, and the procedure stops early because sufficient successes have been observed to estimate the $p$-value with the desired relative precision ($c \cdot \eta = 0.005$). If successes are rare, the procedure continues until $B$ permutations have been performed, giving greater resolution for small $p$-values while keeping computation bounded. In both cases, the permutation $p$-value is estimated as

$$\hat{p} = \frac{r}{b}.$$

As in the preceding section, the direct and SAD variance components are tested with one-sided tests while $p$-values for direct-SAD covariance and non-direct components are evaluated using two-sided tests. We compute lower- and upper-tail permutation probabilities ($\hat{p}_{lower}$ and $\hat{p}_{upper}$, respectively) separately and use twice the smaller probability as the two-sided $p$-value,

$$\hat{p} = 2 \cdot \min(\hat{p}_{lower}, \hat{p}_{upper}).$$

#### S5 Approximate sampling variances for the moment-based variance-component estimators

Here, we derive approximate sampling variances related to the variance component for direct effects along principal component $i$ (or $c_{Di}$) in some detail. At the end of the section, we give approximate sampling variances for the other variance components—the derivations for these are only sketched but are similar. The variances here are not used to compute the null non-rejection regions produced by PGSUS and shown in the main text. They are instead computed via permutations as in the previous section. We report the approximate standard errors here for completeness.

The variance component for direct effects along principal component $i$

$$c_{Di} = \lambda_i \left( \sum_j \beta_{Dj} U_{ij} \right)^2$$

is closely related to the squared projection of the sib-GWAS allelic effect estimates on the $i$th principal component, $s_i^2$ or

$$s_i^2 = \lambda_i \left( \sum_j (\beta_{Dj} + \epsilon_{Sj}) U_{ij} \right)^2.$$

The expectation of $s_i^2$ is

$$\mathrm{E}[s_i^2] = \lambda_i \left[ \mathrm{E}\left[ \left( \sum_j \beta_{Dj} U_{ij} \right)^2 \right] + \mathrm{E}\left[ \left( \sum_j \epsilon_{Sj} U_{ij} \right)^2 \right] + 2\mathrm{E}\left[ \left( \sum_j \beta_{Dj} U_{ij} \right) \left( \sum_j \epsilon_{Sj} U_{ij} \right) \right] \right]$$

Noticing that the first expectation inside the brackets is of a constant, $c_{Di}/\lambda_i$, and that $\mathrm{E}\left[\sum_j \epsilon_{Sj} U_{ij}\right] = 0$ gives the simplified form

$$\mathrm{E}[s_i^2] = c_{Di} + \mathrm{Var}\left[ \sum_j \epsilon_{Sj} U_{ij} \right].$$

An equivalent derivation is the basis of the proposed moment estimator for $c_{Di}$.

The variance of $s_i^2$ is

$$\mathrm{Var}[s_i^2] = \mathrm{Var}\left[ \lambda_i \left[ \left( \sum_j \beta_{Dj} U_{ij} \right)^2 + \left( \sum_j \epsilon_{Sj} U_{ij} \right)^2 + 2 \left( \sum_j \beta_{Dj} U_{ij} \right) \left( \sum_j \epsilon_{Sj} U_{ij} \right) \right] \right].$$

Because $\sum_j \beta_{Dj} U_{ij}$ is a constant, the variance becomes

$$\begin{aligned} \mathrm{Var}[s_i^2] = &\lambda_i^2 \left[ \mathrm{Var}\left[ \left( \sum_j \epsilon_{Sj} U_{ij} \right)^2 \right] + 4 \left( \sum_j \beta_{Dj} U_{ij} \right)^2 \mathrm{Var}\left[ \sum_j \epsilon_{Sj} U_{ij} \right] \right. \\ &\left. + 4 \left( \sum_j \beta_{Dj} U_{ij} \right) \mathrm{Cov}\left[ \left( \sum_j \epsilon_{Sj} U_{ij} \right)^2, \sum_j \epsilon_{Sj} U_{ij} \right] \right]. \end{aligned} \quad (37)$$

This is as far as the expression can be simplified using only the assumption that $\mathrm{E}[\epsilon_{Sj}] = 0$ for all $j$. Because we assume that only the $\epsilon$ terms are random, it is possible in principle to estimate the terms in **Eq. 37** by jackknifing or bootstrapping individuals (or, in this case, sibling pairs) from the original study used to estimate the allelic effect. We do not pursue this option here because it is demanding computationally and requires access to individual-level GWAS data that we do not have in every case we examine. As such, we proceed by making assumptions about the distribution of the error terms and their covariance.

If we assume further that $\mathrm{E}[(\sum_j \epsilon_{Sj} U_{ij})^3] = 0$—one sufficient but not necessary condition that guarantees this is that the error terms are symmetrically distributed around zero with a defined third moment— then the covariance term in **Eq. 37** becomes zero, and the third term inside the brackets vanishes. Next, if we adopt the assumption that $\sum_{j=1}^{\ell} \sum_{k \neq j} U_{ij} U_{ik} \mathrm{E}[\epsilon_{Sj}\epsilon_{Sk}] = 0$—already adopted in the main text, see **Eq. 16** for the analogous assumption for the standard-GWAS allelic effect estimates—then

$$\mathrm{Var}\left(\sum_j \epsilon_{Sj} U_{ij}\right) = \sum_j U_{ij}^2 \mathrm{Var}[\epsilon_{Sj}],$$

which simplifies the second term in the brackets of **Eq. 37**. The first term in brackets, $\mathrm{Var}\left(\left(\sum_j \epsilon_{Sj} U_{ij}\right)^2\right)$, depends on the kurtosis of the projection of the errors on the $i$th principal component, $\sum_j \epsilon_{Sj} U_{ij}$, and is thus sensitive to distributional assumptions. We assume that $\sum_j \epsilon_{Sj} U_{ij}$ has a Normal distribution. One way to defend a Normality assumption as approximately valid is via a central limit theorem argument, as $\sum_j \epsilon_{Sj} U_{ij}$ is a sum of many random variables. Though the variables in the sum are neither independent nor identically distributed, LD clumping should help ensure that their correlations are not too large, and their variances will typically not differ too drastically. We do not pursue a formal justification of the normality assumption.

The three assumptions in the previous paragraph specify that $\sum_j \epsilon_{Sj} U_{ij}$ has a Normal distribution with expectation 0 and variance $\sum_j U_{ij}^2 \mathrm{Var}[\epsilon_{Sj}]$. It follows that the projection of the sib-GWAS allelic effect estimates on principal component $i$, or $\sum_j (\beta_{Dj} + \epsilon_{Sj}) U_{ij}$, has a Normal distribution with the same variance and expectation $\sum_j \beta_{Dj} U_{ij}$. Thus, the quantity

$$\frac{\sum_j (\beta_{Dj} + \epsilon_{Sj}) U_{ij} - \sum_j \beta_{Dj} U_{ij}}{\sqrt{\sum_j U_{ij}^2 \mathrm{Var}[\epsilon_{Sj}]}}$$

has a standard normal distribution, and its square has a $\chi^2(1)$ distribution, meaning it has variance 2. It follows that under these assumptions, the variance of the squared projection of the sib-GWAS allelic effect estimate estimates on principal component $i$ is

$$\mathrm{Var}[s_i^2] = 2\lambda_i^2 \left[\sum_j U_{ij}^2 \mathrm{Var}[\epsilon_{Sj}]\right]^2. \tag{38}$$

Our estimator of the component of variance due to direct effects along principal component $i$, $c_{Di}$, is (**Eq. 20**)

$$\hat{c}_{Di} = \lambda_i \left( \sum_{j=1}^{\ell} \beta_{Sj} U_{ij} \right)^2 - \lambda_i \sum_{j=1}^{\ell} U_{ij}^2 \hat{v}_{Sj}^2.$$

The first term is $s_i^2$. The variance of $\hat{c}_{Di}$ is

$$\begin{aligned}\mathrm{Var}[\hat{c}_{Di}] &= \mathrm{Var}[s_i^2] + \lambda_i^2 \mathrm{Var}\left[ \sum_{j=1}^{\ell} U_{ij}^2 \hat{v}_{Sj}^2 \right] - 2\lambda_i^2 \mathrm{Cov}\left[ \left( \sum_{j=1}^{\ell} \beta_{Sj} U_{ij} \right)^2, \sum_{j=1}^{\ell} U_{ij}^2 \hat{v}_{Sj}^2 \right] \\ &= \mathrm{Var}[s_i^2] + \lambda_i^2 \sum_{j=1}^{\ell} U_{ij}^4 \mathrm{Var}[\hat{v}_{Sj}^2] - 2\lambda_i^2 \mathrm{Cov}\left[ \left( \sum_{j=1}^{\ell} \beta_{Sj} U_{ij} \right)^2, \sum_{j=1}^{\ell} U_{ij}^2 \hat{v}_{Sj}^2 \right], \end{aligned} \tag{39}$$

where the second step holds if the variance estimates at distinct loci are uncorrelated. In practice, one could plausibly ignore the second and third terms in **Eq. 39** because they might be expected to be much smaller than $\mathrm{Var}[s_i^2]$.

To justify ignoring the second term in **Eq. 39**, we need to argue that

$$2 \left( \sum_j U_{ij}^2 \mathrm{Var}[\epsilon_{Sj}] \right)^2 >> \sum_j U_{ij}^4 \mathrm{Var}[\hat{v}_{Sj}^2]. \tag{40}$$

First, we note that

$$\left( \sum_j U_{ij}^2 \mathrm{Var}[\epsilon_{Sj}] \right)^2 > \sum_j U_{ij}^4 \mathrm{Var}[\epsilon_{Sj}]^2,$$

and so the inequality in **Eq. 40** holds if

$$2 \sum_j U_{ij}^4 \mathrm{Var}[\epsilon_{Sj}]^2 >> \sum_j U_{ij}^4 \mathrm{Var}[\hat{v}_{Sj}^2].$$

In turn, this inequality holds if, for all loci indexed by $j$,

$$2\mathrm{Var}[\epsilon_{Sj}]^2 >> \mathrm{Var}[\hat{v}_{Sj}^2]. \tag{41}$$

In the case of the sib-GWAS allelic effect estimate estimates, we use resampling to estimate the variance of the allelic effect estimates, so we do not have a closed form for the variance estimator or its variance, $\mathrm{Var}[\hat{v}_{Sj}^2]$. However, we expect these quantities to be of roughly the same magnitude as they would be in standard linear regression. In a simple linear regression with $n$ observations, an independent variable $x$, and error variance $\eta^2$, the residual sum of squares (RSS) is distributed according to $\mathrm{RSS}/\eta^2 \sim \chi^2(n-2)$. It follows that the RSS has variance $2(n-2)\eta^4$. In the same setting, the variance of the slope estimator—analogous to $\mathrm{Var}[\epsilon_{Sj}]$ in

**Eq. 41**—is $\text{Var}[\epsilon_{Sj}] = \eta^2/\sum((x_i - \bar{x})^2)$, and this quantity is estimated by $\text{RSS}/((n-2)\sum(x_i - \bar{x})^2)$. The variance of the estimator of the variance of the slope estimator—analogous to $\text{Var}[\hat{v}_{Sj}^2]$ in Eq. 41—is $\text{Var}[\hat{v}_{Sj}^2] = 2\eta^4/((n-2)(\sum(x_i - \bar{x})^2)^2)$. Thus, using our notation, in the simple linear regression setting,

$$\frac{2\text{Var}[\epsilon_{Sj}]^2}{\text{Var}[\hat{v}_{Sj}^2]} = n - 2,$$

and therefore the inequality in **Eq. 41** holds for large $n$. Because all the allelic effect estimates we consider are based on large samples, if the linear regression setting is a valid approximate guide to the variability of the variance estimators obtained by resampling, then the inequality in **Eq. 40** will hold, and we are justified in ignoring the second term of **Eq. 39**. In turn, if the second term of **Eq. 39** is much smaller than the first term, then the third term is as well, as the magnitude of the covariance term is bounded from above by the geometric mean of its two terms.

Thus, to estimate the variance of our estimator of the variance component of the PGS due to direct effects along principal component $i$, we could use

$$\text{Var}[\hat{c}_{Di}] \approx \text{Var}[s_i^2] \approx 2\lambda_i^2 \left( \sum_j U_{ij}^2 \widehat{\text{Var}}[\epsilon_{Sj}] \right)^2 \tag{42}$$

where $\widehat{\text{Var}}[\epsilon_{Sj}]$ is an estimate of the variance of the sib-GWAS allelic effect estimate at locus $j$.

To estimate the variance of the estimator for the variance component of the PGS due to SAD effects along principal component $i$, we step through a similar set of approximations and assumptions for both sib-GWAS and standard-GWAS allelic effect estimates, including the assumption that measurement errors at distinct loci are uncorrelated, that the projection of the measurement errors along principal component $i$ are normal, and that variance due to uncertainty in estimating the per-locus sampling variances is small enough to ignore. We also add the assumption that measurement errors for sib-GWAS and standard GWAS are independent. Doing so gives the expression

$$\text{Var}[\hat{c}_{\sigma i}] \approx 2\lambda_i^2 \left( \sum_j U_{ij}^2 (\widehat{\text{Var}}[\epsilon_{Sj}] + \widehat{\text{Var}}[\epsilon_{Gj}]) \right)^2, \tag{43}$$

where $\widehat{\text{Var}}[\epsilon_{Gj}]$ is an estimate of the measurement error of the standard-GWAS allelic effect estimate at locus $j$.

To estimate the variance for the estimated variance component due to the covariance of direct effects and stratification along principal component $i$, we use the same assumptions of normality of the projections, uncorrelated measurement errors at distinct loci, and independence of sib-GWAS and standard-GWAS estimation errors. Defining $X = \sum_{j=1}^{\ell} \beta_{Gj} U_{ij}$ and $Y = \sum_{j=1}^{\ell} \beta_{Sj} U_{ij}$, and ignoring the final term of the third line of **Eq. 22**, we can write the approximate variance of $\hat{c}_{(D\cdot\sigma)i}$ as

$$\mathrm{Var}[\hat{c}_{(D\cdot\sigma)i}] \approx 4\lambda_i^2 \mathrm{Var}[XY - Y^2]. \tag{44}$$

$X$ and $Y$ are independent and each has expectation 0, so

$$\mathrm{Var}[XY] = \mathrm{E}[X^2Y^2] - (\mathrm{E}[XY])^2 = \mathrm{E}[X^2]\mathrm{E}[Y^2] = \mathrm{Var}[X]\mathrm{Var}[Y].$$

(The first step comes from the definition of the variance, the second step from noting that $\mathrm{E}[XY] = 0$ because $X$ and $Y$ are uncorrelated with expectation 0 and also that $\mathrm{E}[X^2Y^2] = \mathrm{E}[X^2]\mathrm{E}[Y^2]$ by the independence of $X$ and $Y$, and the third step from applying the definition of variance, remembering that $\mathrm{E}[X] = \mathrm{E}[Y] = 0$.) Further, and again by the independence of $X$ and $Y$ and by the fact that $\mathrm{E}[X] = \mathrm{E}[Y] = 0$, $\mathrm{Cov}[XY, Y^2] = 0$. Applying these facts and our other assumptions and substituting $X$ and $Y$ with their definitions gives

$$\mathrm{Var}[\hat{c}_{(D\cdot\sigma)i}] \approx 4\lambda_i^2 \left[ \left( \sum_j U_{ij}^2 \widehat{\mathrm{Var}}[\epsilon_{Sj}] \right) \left( \sum_j U_{ij}^2 \widehat{\mathrm{Var}}[\epsilon_{Gj}] \right) + 2 \left( \sum_j U_{ij}^2 \widehat{\mathrm{Var}}[\epsilon_{Sj}] \right)^2 \right]. \tag{45}$$

#### S6 GWAS and PGS construction

**Standard GWAS using a fixed effects model.** We performed six distinct GWAS for each trait, four of which were in a fixed effect model. We performed GWAS using *PLINK 2.0*'s generalized linear model flag (`--glm`) with age and sex as covariates. The four fixed effect model GWAS we ran used different sets of additional covariates:

(a) The first 20 PCs of the GWAS sample's genotype matrix

(b) The first 20 PCs of the prediction sample's genotype matrix (either the 1KG or the 1KG Europeans samples)

(c) Both sets of PCs as in (a) and (b).

(d) No additional covariates

For (b), we first computed the SNP loadings for the PC space defined by the prediction sample genotype matrix. We then used *FlashPCA2* [7] to project individuals from the GWAS sample into that PC space (with the `--project` flag) and used these individual PC-space coordinates as covariates in our GWAS.

**Standard GWAS using a linear mixed model**. We used the *BOLT-LMM* software[8] to perform two additional GWAS for each trait and prediction sample. We ran linear mixed model regression with age, sex, and allele count at the focal locus as fixed effects and a term with covariance proportional to the genome-wide relatedness matrix as a random effect. We also ran this regression where in addition we included the first 20 PCs of the standard-GWAS sample genotype matrix as fixed effects.

**PGS Construction and obtaining PCs**. As possible PGS index SNPs, we considered the autosomal variants that existed in both the UK Biobank imputed SNP set and in the 1KG data. We used a clumping and thresholding strategy[9–11]. In particular, for each trait, we used *PLINK 1.90*'s greedy clumping algorithm (flags `--clump-p1 1` `--clump-r2 0.1` `--clump-kb 250` `--clump-best`) to subset SNPs into clumps. Variants were ranked using their standard-GWAS marginal association $p$-values. Starting with the most significantly associated variant, all variants within 250 kilobases of that index variant and with an $r^2$ value greater than 0.1 were assigned to that index variant's clump. This process was repeated until no variants remained, resulting in roughly 350,000 clumps for each trait. The number of clumps with an index SNP $p$-value less than each of the four GWAS association $p$-value thresholds we used can be found in **Table S2** for the fixed effects model standard GWASs using age, sex, and the first 20 PCs of the GWAS. Additionally, summary statistics for all of the standard-GWAS models and corresponding sibling-GWAS models can be found on the Harpak Lab data page (`https://www.harpaklab.com/data`).

**sib-GWAS**. For each quantitative trait that we analyzed, we performed a sib-GWAS using *PLINK 1.9*'s sib-GWAS procedure implemented in the `--qfam` flag on individuals' phenotypes residualized for age and sex.

We estimated standard errors for each of the allelic effect estimates following the block-jackknife method described in Miao et al.[12]. This approach uses block-jackknife resampling to generate unbiased estimates of standard errors in sib-GWASs. Briefly, for each resampling $m$, a portion of the siblings is randomly dropped from the dataset and the allelic effect, $\beta^*_{S,m}$, is estimated for this subsample. For our analyses, we set the number of dropped sibling pairs per resampling at $d$=500 and the total number of resamplings was $M$=500. The standard error of the allelic effect is then estimated from the standard deviation across all replicates. Namely, per **Eq. 44** in Miao et al.[12], the standard error of the sib-GWAS allelic effect estimate for the focal SNP is estimated as:

$$\mathrm{SE}_{bjk}[\beta^*_S] = \sqrt{\frac{r}{d\cdot(M-1)}\sum_{m=1}^{M}\left(\beta^*_{S,m} - \bar{\beta}^*_{S,bjk}\right)^2},$$

where $\bar{\beta}^*_{S,bjk}$ is the average of $\beta^*_{S,m}$:

$$\bar{\beta}^*_{S,bjk} = \frac{1}{M}\sum_{m=1}^{M}\beta^*_{S,m},$$

and $r = n - d$ is the number of remaining sibling pairs.

**GWAS of seven binary health conditions from the UK Biobank.** In addition to our analysis of the 17 continuous traits in the UK Biobank, we analyzed seven binary health conditions (**Tables S3**, **S4**). We performed standard and sib-GWASs in the same White British cohorts that we analyzed for the 17 continuous traits. In addition to these seven traits, we also analyzed BMI but modeled the phenotype using the definition of BMI > 30 to label individuals as obese or non-obese. We treated the disease phenotypes as continuous variables in both GWASs in order to maintain the same statistical assumptions made in the PGSUS protocol and to illustrate its potential extension to binary health conditions. For the standard GWAS we used age, sex, and the first 20 PCs of the GWAS cohort as covariates and performed the same clumping and thresholding procedure described in the preceding section. For each trait, we used corresponding sib-GWAS summary statistics to decompose the variance in PGSs for both the 1KG and 1KG European prediction samples using the PGSUS framework. **Fig. S29** shows a selection of SAD and non-direct variance components that were significantly associated with top PCs of different target samples for a number of traits.

**Consequences of index SNPs ascertainment procedure.** In the clumping and thresholding approach on which we focus in the main text, genetic variants are selected as index SNPs (i.e., selected for inclusion in a polygenic score) on the basis of the significance of association with the phenotype. This significance is a function of estimated genetic variance contributed by a SNP,

$$\hat{\beta}^2 p(1 - p),$$

where $p$ is the frequency of the effect allele and $\hat{\beta}$ is the allelic effect estimate (assuming the index SNP is not in LD with other index SNPs). This process of choosing SNPs on the basis of their allelic effect estimates from a GWAS can affect the quantities we estimate and their interpretation. In **Text S11** and **Text S14** we discuss and explore with simulation studies how ascertainment of loci with large allelic effect estimates in GWAS affects estimates of the isotropic inflation factor $\alpha$, the non-direct variance component, and of PC-specific variance components. In our empirical tests, we found that 4 of 5 PGSs which showed significant SAD variance when index SNPs were ascertained using standard-GWAS summaries (**Fig. 4**) showed no significant SAD variance components when index SNPs were selected based on sib-GWAS summaries (**Fig. S30**).

#### S7 Multiple hypothesis testing for PC-specific SAD variance components

In the main text, we evaluate evidence for large PC-specific SAD variance components, suggestive of stratification. The analyses highlight tests for the top 6 PCs in 17 traits for PGSs derived from UKB WB-based GWAS applied to the 1KG prediction sample. Here, we describe a multiple hypothesis correction used for the omnibus null hypothesis of no PC-specific SAD effects at play for any of these PGSs and the resultant $p$-value for the empirical observation of 12 of the 17 PGSs analyzed having at least one significantly large SAD variance component in the top 6 PCs.

Assuming the tests along each PC are independent, the probability of obtaining significant SAD components on at least one of the first six PCs under this null hypothesis is $1 - 0.95^6$. Across all PGSs analyzed and assuming that tests of distinct traits are also independent, the probability of obtaining significant PC-specific SAD components on 12/17 or more traits under the null hypothesis is $p = 1.8 \times 10^{-4}$.

### S8 Procurement and processing of external data

#### S8.1 GIANT consortium GWAS data for height (Wood et al. 2014 and Yengo et al. 2022).

GWAS summary statistics for analyses of height using different versions of the GIANT meta-cohort were downloaded from `https://portals.broadinstitute.org/collaboration/giant/index.php/GIANT_consortium_data_files`. For Wood et al.[13] and Yengo et al.[14], we found the set of variants that overlapped with both the 1000 Genomes phase 3 data (1KG) and our array of 9.6 million SNPs from the UK Biobank. We then performed clumping (using the same set of parameters described in **Text S6**) using the CEU and GBR individuals from the 1KG European superpopulation.

#### S8.2 Ancient DNA sample selection and principal components analysis

In the main text, we discuss PGSUS results for PGSs for 10 of the continuous traits in the UK Biobank which we applied to a sample of ancient Eurasians from the Allen Ancient DNA Resource (AADR, v62.0)[15,16]. This analysis was performed using the 1240K + H0 files from the Allen Ancient DNA Resource. Files were converted from eigenstrat format to PED format and then to PLINK bfiles with EIGENSOFT v8.0.0[17,18]. Quality control steps for SNPs included in the analysis can be found in Mallick et al.[16]. We further restricted SNPs to those which overlapped the array used for sequencing of the UK Biobank participants and those with minor allele frequency of at least 0.001 using PLINK 2.0[19] (options `--maf 0.001`).

Selection of samples and SNPs followed that of Akbari et al.[20]. After filtering, we were left with 4,588 ancient individuals and 882,044 SNPs. Each individual was assigned to a population group: Paleolithic (P, n=16), Mesolithic (M, n=125), Neolithic (N, n=791), Chalcolithic (C, n=240), Bronze Age (BA, n=1,457), Iron Age (IA, n=746), Steppe Pastoralist (S, n=301), Medieval (Med, n=654), or Modern (Mod, n=221). 37 individuals did not have a population assigned. Groups were assigned using the method and notation described in the AADR release v62.0 documentation and metadata[16]. Briefly, population assignment for each sample is determined using geographic location, $f_4$-statistics, and sample date as determined by radiocarbon dating or well-known anthropological information. 37 individuals did not have a provided population assignment.

Because DNA damage is time-dependent and affected by environmental conditions that may be correlated with population structure, we computed the correlation between the first 10 principal components and various features of the sequencing data including coverage, sample date, and location (**Fig. S13**). For all of the variables we evaluated, we found significant correlations with at least two of the top ten PCs.

#### S9 The utility of population structure adjustments for GWAS in mitigating confounding

In the main text section **The relative contribution of confounding to variation in a PGS**, we present results showing that the addition of PCs or the use of linear mixed models (LMMs) to adjust GWAS allelic effect estimates can reduce non-direct variance in PGSs (**Fig. 2C**; **Text S6**). Here, in addition to evaluating the utility of the six GWAS adjustment methods in mitigating non-direct variance, we evaluate their utility in mitigating signals of stratification (significant PC-specific SAD variance) and isotropic inflation. In addition to the random subset of the UKB we analyze in the main text, we repeat the analysis using the UKB WB sample. We show that in this less structured sample, our three evaluation metrics generally suggest GWAS adjustments have little to no effect.

Our expectation was that no PC adjustment in the GWAS should lead to equal or larger confounding in all three respects: larger non-direct variance components, equal or more significant PC-specific SAD components, and larger isotropic inflation factors. Using a random sample of 100,000 individuals from the full UKB cohort, we performed GWAS using each adjustment method and computed the non-direct variance component, isotropic inflation factor, and variance partitionings for the corresponding PGSs. We used the UKB WB sib-GWAS summary statistics for our analysis (while this can introduce SAD effects, see **Section S14.8**, we note that our random sample is comprised of 87% White British individuals).

We found that both PC-based adjustments and LMMs provide significant reduction in the proportion of non-direct variance for a PGS as compared to either no adjustment or the use of prediction sample PCs as GWAS covariates ($p < 0.027$ for all comparisons, **Fig. S4A**, **File S9**). With respect to the presence of PC-specific SAD components in the top 6 PCs, only the addition of prediction sample PCs provided a significant reduction in SAD variance (McNemar $p = 0.008$ when compared to no adjustment, **Fig. S4B**, **File S9**). Results were approximately similar when looking at the impact of GWAS adjustments on isotropic inflation, with all methods significantly reducing isotropic inflation factors as compared to no adjustment or to prediction sample PCs (paired t-test $p < 0.05$ for pairwise comparisons, **Fig. S4C**, **File S9**).

For GWASs based on the UKB WB cohort, we find that GWAS adjustment methods did little to reduce non-direct variance components ($p > 0.05$ for all pairwise comparisons except between prediction sample PCs and LMM-based GWAS; **Fig. S6A**; **File S10**; also see **Fig. S31** for the correlations between GWAS and prediction sample PCs). Surprisingly, the GWAS sample PC adjustment method resulted in the highest number of PGSs with PC-specific SAD components among the top 6 PCs (McNemar $p < 0.05$ compared to either no adjustment or prediction sample PCs; **Fig. S6B**; **File S10**). This result may be due to the relatively homogeneous UKB WB sample. In the absence of PC-specific SAD effects, and assuming independence among tests for PC-specific tests on the first 6 PCs, we expect that of the 17 traits we examined, $17(1-.95^6) \approx 4.5$ PGSs should have significant PC-specific SAD variance on the first six PCs. With no adjustment for population structure, we observe such significant effects on only 6 of the 17 PGSs, which is not significantly different from the null expectation by a one-tailed binomial test ($p = 0.282$; **Text S7**). Finally, GWAS adjustment methods did not consistently perform better than no adjustment in reducing isotropic inflation (**Fig. S6C**; **File S10**).

#### S10 Isotropic inflation factor

In the main text, we present "isotropic inflation", as the systematic inflation of standard GWAS summary statistics compared with corresponding sibling GWAS summary statistics. Here, we discuss—and provide support via simulations for—various possible drivers of isotropic inflation, including SAD effects as well as technical considerations.

Empirically, among a set of PGSs (UKB-based, with clumping and thresholding on a GWAS association $p$-value $< 10^{-5}$) applied to the 1KG prediction sample, nearly all PGSs show significant isotropic inflation (**Fig. S20B**). When we constructed PGSs differently, using a GWAS association $p$-value $< 10^{-8}$ for index SNPs, the rankings of isotropic inflation factors across traits changed substantially, illustrating again that PGSUS characterizes the variance of a specific PGS in a given prediction sample, and not a trait or a GWAS (**Text S6; Figs. S20-S22**).

**What drives isotropic inflation?** In principle, isotropic inflation could result from each of the SAD effects we explore in the main text: assortative mating[21–26], indirect parental effects[24,26–31], or some modes of population stratification[1,26,32–36].

Confounding due to stratification can act isotropically when the environmental resemblance among individuals is proportional to their genetic relatedness, i.e. the genetic relatedness matrix. Such covariance between environmental and genetic features is unlikely to act along the axis of stratification captured by a single PC, instead leading to inflation of standard-GWAS summary statistics with respect to many PCs. In simulations, we examined case studies wherein we found that isotropic inflation increased with increased stratification (**Fig. S7G,I**).

Assortative mating plays a role in complex trait variation in human populations[25,37–42] and, similar to stratification, its effects on genetic variance may not localize to an individual principal component. In fact, some forms of assortative mating produce patterns that conform to the isotropic inflation factor assumed here, in which the allelic effect estimate is inflated by a scalar with size dependent on the strength of assortative mating[30,43]. In simulations, we show an increase in isotropic inflation with increasing degrees of assortative mating (**Fig. S17A**).

Indirect parental, or dynastic, effects are also often thought of as acting isotropically, at least in part[24,28,31,44]. In our simulations of dynastic effects, we found that, as expected, dynastic effects which were positively correlated with direct effects resulted in isotropic inflation factors above 1, dynastic effects which were negatively correlated with direct effects resulted in isotropic inflation values less than 1, and dynastic effects with no correlation to direct effects resulted in approximately null ($\alpha = 1$) isotropic inflation (**Fig. S18**).

There are also technical drivers of isotropic inflation, for example, differences in units of measurement among studies (e.g., centimeters as opposed to standard deviations of height). Another technical factor driving isotropic inflation may be in index-SNP ascertainment biases. Winner's curse[45,46] may lead allelic effect estimates to be systematically larger in the sample in which the effects were ascertained as significant, as we show in simulations (**Fig. S19A**; **Text S14.3**). Throughout our analyses here, ascertainment is based on significance of marginal SNP association in the standard GWAS, and so intuition dictates that allelic effect estimates would be larger in the standard GWAS and the isotropic inflation factor would be driven to be larger than 1. In our winner's curse simulation, when we instead ascertained on sib-GWAS effects, we saw isotropic inflation factor estimates which were lower than 1 and became smaller with increased ascertainment stringency (**Fig. S19B**). Empirically, our estimates of $\alpha$ were highly sensitive to the choice of a marginal GWAS association $p$-value (**Figs. S20-S22, S32**).

There are likely additional ascertainment biasing effects at play[1,34]. One example is clumping, a strategy to account for LD in the ascertainment of index SNPs[10,19]. Typically, in the clumping and thresholding technique, the clumps are chosen, through a greedy algorithm, to be centered around the most significant SNPs that are not yet included in other clumps. We hypothesized that because SNPs in the same clump are in LD, using even random SNPs as index SNPs would induce ascertainment bias and increase isotropic inflation, as long as these SNPs were chosen from the "best" clumps, i.e. clumps centered around a highly significant SNP. Indeed, choosing index SNPs in this manner still resulted in isotropic inflation factors greater than 1 for most PGSs (second row in **Fig. S32**). (We did not observe a consistent effect of the stringency of ascertainment on the rate of significant PC-specific components when comparing the same selection approaches (**Fig. S33**)).

We also considered that isotropic inflation may be due to ancestry-related participation bias in GWAS or sib-GWAS cohorts. We performed a standard GWAS with a sample containing individuals from two subpopulations (split 100 generations prior from a single population) in equal proportions and a sib-GWAS with individuals drawn from only one of the subpopulations. As expected, ancestry-related participation bias results in increased isotropic inflation ($\bar{\alpha}$ = 3.52 across 100 simulations; **Text S14.8**).

Finally, we note that, given that allelic effect estimates and their standard errors are used to estimate $\alpha$ (**Eq. 23**), its estimation may be affected by GWAS sample size or any other factors (e.g., trait architecture) that affect the power to detect associations in the GWAS.

We were not able to identify a strong predictor for variation in isotropic inflation factors across traits. Isotropic inflation estimates in 1KG Europeans were not significantly correlated with LD score regression[47]-based SNP heritability estimates across traits (**Fig. S34**). Further, we found that the correlation coefficient between individual trait values and Townsend deprivation index, a composite metric of socioeconomic status, were not correlated with the isotropic inflation factor estimates of the same traits (**Fig. S35**). Estimates of the isotropic inflation factor also depended on the prediction sample (e.g., when comparing **Fig. S20** and **Fig. S21**).

#### S11 Index SNP ascertainment in standard GWAS (sib-GWAS) increases (decreases) isotropic inflation

In **Text S10** and **Text S14.3** we show that more stringent SNP ascertainment leads to larger isotropic inflation factor estimates, primarily as a result of the winner's curse[48,49]. Here, we further solidify this conclusion with simulations using real UKB genotypes for the UKB White British (UKB WB) cohort. We then show that ascertainment based on sib-GWAS, which results in the systematic selection of SNPs with larger allelic effects in the *sib-GWAS* sample, results in smaller isotropic inflation factor estimates.

For each iteration of our simulations, we computed a phenotype value for each individual as the sum of a genetic and an environmental contribution to the phenotype. For the genetic contribution, 1% of genotyped and imputed variants were randomly selected to be causal—that is, to directly contribute to the variation in the phenotype. Allelic effects were selected at random from a standard normal distribution for these variants and were scaled such that they explained 50% of the total variance in the phenotype. An environmental component of the phenotype was then constructed by randomly selecting an environmental contribution for each individual from a standard normal distribution, such that the variance in those effects was equal to 50% of total phenotypic variance. That is, the environmental contributions were scaled to explain 50% of the total phenotypic variation. For the sibling cohort, the correlation in environmental contributions to the phenotype between siblings was set to be 0.5. We then performed standard and sib-GWASs using the real genotypes and simulated phenotypes, and built polygenic scores using clumping and thresholding as described in **Text S6** using one of four different standard-GWAS ascertainment thresholds: $p$-value < 1, $p$-value < $10^{-3}$, $p$-value < $10^{-5}$, or $p$-value < $10^{-8}$. We then performed PGS partitioning with PGSUS using the 1KG Europeans as the prediction sample and estimated the isotropic inflation factor. We performed 100 iterations for each test.

Our results show a similar pattern to our empirical analyses (**Figs. S20-S22**; **Text S10**) and simulations using simulated genotypes (**Fig. S19A; Text S14.3**). Namely, more stringent ascertainment leads to an increase in isotropic inflation (**Fig. S36**).

We repeated the analysis with ascertainment based on the sib-GWAS summaries. To account for differences in statistical power between the standard and sibling approaches, we did not use nominal $p$-value thresholds in the manner described in our standard-GWAS ascertainment approach. Instead, we matched each of the four standard-GWAS-based PGSs with a PGS with the same number of SNPs, ascertained by applying the clumping and thresholding approach to the sib-GWAS summary statistics. Here, more stringent thresholding based on sib-GWAS statistics leads to smaller estimates of isotropic inflation (**Fig. S36**), matching the pattern we observed in our simulations with simulated genotypes (**Fig. S19B; Text S14.3**).

#### S12 Comparison of isotropic inflation factor with measures of heritability and confounding

Across the 17 traits in UKB analyzed in the main text, we evaluated the relationship between our estimates of the isotropic inflation factor with several field-standard measures of SNP heritability and confounding, applied to the corresponding GWAS summary statistics.

First, we examined the slope and intercept in LD score regression (LDSC)[47]—measures of SNP heritability and inflation in GWAS significance due to population structure confounding, respectively (**Fig. S34**). We observed that both LDSC slopes and intercepts are only correlated with estimates of isotropic inflation at a GWAS ascertainment threshold of $p$-value < 1 (**Fig. S34G,H**; we did not observe a significant correlation between LDSC-based estimates and the non-direct variance component, see **Fig. S37**). We hypothesize that this might be in part due to the fact that LDSC uses all variants to estimate heritability parameters, thus estimates of isotropic inflation using more stringent thresholds do not use the same set of variants as LDSC. It is additionally important to note that isotropic inflation is a parameter of a variance in a given PGS in a given prediction sample, so its estimate will depend on the target population.

Next, we compared our estimates of the isotropic inflation factor to another estimate of SNP heritability due to assortative mating (which we expect to act, at least in part, isotropically). Zhang et al.[50] estimated the contribution of "gametic phase disequilibrium" (GPD), or the occurrence of high rates of linkage disequilibrium among physically unlinked variants that result from assortative mating, to SNP heritability. The contribution of this phenomenon to trait variance is captured by what Zhang et al.[50] label the disequilibrium genetic relatedness matrix (DGRM), which explicitly models the difference in the expected covariance among genotypes versus what is observed at variants with high GPD. The DGRM is used in conjunction with the GRM to then decompose the sources of genetic contribution to phenotypic variation into that explained by standard additive contribution of variants and that explained by GPD (i.e. explained by the effects of assortative mating). Per the recommendation of the authors of Zhang et al.[50], we obtained estimates of the proportion of phenotypic variance explained by the DGRM (or DGRM heritability) by taking the mean estimate from seven non-overlapping cohorts of the UKB White British individuals included in our standard GWAS. The primary motivation for this approach is the memory constraint imposed by estimating and modeling the GRM and DGRM for large cohorts. We use the DGRM heritability estimate as an external estimator of the contribution of assortative mating and evaluate if it correlates with isotropic inflation as we hypothesize it should.

We only observe a significant correlation between the mean DGREML heritability and isotropic inflation estimates for PGS using clumping with less stringent thresholding for the 17 traits we analyzed applied to either the 1KG or the 1KG Europeans prediction sample (**Fig. S38F-H**). As in our comparison of the isotropic inflation factor to estimates from LDSC, this may partially result from the fact that the index SNPs for these PGSs are more similar to the genome-wide set of SNPs underlying DGREML estimates.

#### S13 Relationships between PGSUS summaries

##### S13.1 Isotropic inflation and non-direct variance

Here, we consider empirical observations and theoretical expectations for the relationship between the isotropic inflation estimates and the non-direct variance component.

We first analyzed raw Pearson correlations in a set of PGSs based on the UKB WB GWAS for the 17 traits in **Table S1** at two different GWAS ascertainment thresholds ($p$-value < $10^{-5}$ and $p$-value < $10^{-8}$) applied to two prediction samples (1KG and 1KG Europeans). The two summaries were not significantly correlated in our tests (**Fig. S23**).

The lack of correlation made us revisit our expectation. In particular, we asked whether we should expect a linear — or even monotonic — relationship between the two. We considered the expectation under a theoretical model where isotropic inflation is the sole driver of non-direct variance. In the absence of PC-specific SAD effects,

$$c_{\sigma i} = c_{(D\cdot\sigma)i} = 0$$

for each PC $i$. As described in **PGSUS-based parameter estimation**, the non-direct variance component, $V_N$, is estimated before the sib-GWAS effects are adjusted for isotropic inflation by setting $\beta_S^* = \beta_S$. In this partitioning, the direct variance component of the $i$th PC estimated by unadjusted sib-GWAS allelic effects is:

$$c_{Di}^* = \lambda_i \sum_{j=1}^{\min(n,\ell)} \left(U_{ij} \cdot \beta_{Sj}^*\right)^2 .$$

After estimating the isotropic inflation factor, we rescale the sib-GWAS allelic effect estimates so that they are on the same scale as the standard-GWAS allelic effect estimates (**Eq. 11**). The corresponding direct variance component after rescaling is:

$$\begin{aligned} c_{Di}^* &= \lambda_i \sum_{j=1}^{\min(n,\ell)} \left(U_{ij} \cdot \alpha^{-1} \beta_{Sj}\right)^2 \\ &= \alpha^{-2} \lambda_i \sum_{j=1}^{\min(n,\ell)} \left(U_{ij} \cdot \beta_{Sj}\right)^2 \\ &= \alpha^{-2} c_{Di}. \end{aligned} \tag{46}$$

We define the sum over all PCs of the direct variance component as $V_D^*$:

$$V_D^* := \sum_{i=1}^{\min(n,\ell)} c_{Di}^* .$$

In the absence of PC-specific SAD effects, the total PGS variance is attributable only to direct variance and isotropic inflation:

$$\mathrm{Var}[Z] = \sum_{i=1}^{\min(n,\ell)} c_{Di} = \alpha^2 \sum_{i=1}^{\min(n,\ell)} c_{Di}^* = \alpha^2 \cdot V_D^* . \tag{47}$$

Additionally, because the non-direct variance component is estimated with unadjusted sib-GWAS allelic effects, the direct variance component in this partitioning is $c_{Di}^*$ in **Eq. 46**. When no PC-specific SAD effects are at play, the numerator of the non-direct variance component is therefore the difference between total variance and direct variance component with unadjusted sib-GWAS allelic effects:

$$V_N = \frac{\mathrm{Var}[Z] - V_D^*}{\mathrm{Var[Z]}} . \tag{48}$$

Then we can combine **Eqs. 47** and **48** as follows:

$$\begin{aligned} V_N &= \frac{\text{Var}[Z] - V_D^*}{\text{Var}[Z]} \\ &= \frac{\alpha^2 V_D^* - V_D^*}{\alpha^2 V_D^*} \\ &= \frac{\alpha^2 - 1}{\alpha^2} \end{aligned}$$

Thus, under a theoretical model where all non-direct variance components are due to isotropic inflation, the relationship between the non-direct variance component and the isotropic inflation factor is monotonic, and in particular fractional,

$$V_N = 1 - \frac{1}{\alpha^2}. \tag{49}$$

This model does not explain variation across PGSs well ($R^2$ = -1.41; **Fig. S25A**). When considering only PGSs with the same GWAS ascertainment threshold, the fit of this model is slightly better among PGSs with more stringent GWAS ascertainment ($R^2 = 0.11$ and $R^2 = 0.009$ for ascertainment at $p < 1 \times 10^{-8}$ and $p < 1 \times 10^{-5}$, respectively; **Fig. S25B,C**).

#### S13.2 Significant PC-specific SAD variance components and non-direct variance

We next investigated the relationship between the non-direct variance component and significant PC-specific SAD variance components. For the same 17 phenotypes from **Table S1** and using the UKB WB as our GWAS cohort, we tested for a mean difference in estimates of the non-direct variance components between PGSs with and without significant PC-specific SAD variance components among the top six PCs using a t-test. In four tests — across two $p$-value thresholds ($p$-value< $10^{-8}$ and $p$-value< $10^{-5}$) and two prediction samples (1KG and 1KG Europeans) we only find a marginally significant difference for 1KG Europeans when using the ascertainment threshold $p$-value < $10^{-5}$ ($p$=0.018; **Fig. S39**).

#### S13.3 Sensitivity of PC-specific SAD variance component estimation to the isotropic inflation factor

In our two-partitionings-based estimation procedure (described in the **Model** section), the estimation of PC-specific variance components is based on sib-GWAS summary statistics that are adjusted by an isotropic inflation factor estimated based on the first partitioning. Here, we ask, using both theory and an empirical case study, whether the estimates of PC-specific components and their significance testing are sensitive to the value of the isotropic inflation factor used as an input for the second partitioning.

**Empirical case study.** We considered PGSs for height based on a UKB WB GWAS, constructed via clumping and thresholding at either $p < 1 \times 10^{-5}$ or $p < 1 \times 10^{-8}$) applied to the 1KG Europeans prediction sample. We use a first partitioning to estimate the isotropic inflation factor (as usual). Then, before performing the second partitioning, we multiplied the sib-GWAS allelic effect estimates and standard errors by an isotropic inflation factor which is larger or smaller than the estimate—$\alpha := 1.314 \times \hat{\alpha}$ or $\alpha := 0.867 \times \hat{\alpha}$, respectively. We found that the significance of SAD variance components, and their nominal values (when standardized by total PGS variance), was largely unaffected by misspecification of the isotropic inflation factor (**Figs. S1, S2**). However, the same was not true for PC-specific non-direct variance components, for which there was one component that was significant with a correctly-specified isotropic inflation factor (PC3 in **Fig. S2B**) but insignificant when using a factor that was too large (**Fig. S2D**).

To examine this in more depth, we compared nominal $p$-values (generated at increased precision using the adaptive $p$-value procedure described in **Text S4**) for variance components for the top 6 PCs corresponding to all four effect types. This examination confirms that, for both too large and too small isotropic inflation factors, $p$-values of PC-specific SAD variance and direct variance components appear unaffected by misspecification (**Fig. S3A,B,E,F**). Conversely, direct-SAD covariance components (and consequently, non-direct variance components) were biased upward. The bias appeared to be roughly proportional to the variance component (**Fig. S3C,D,G,H**).

**Theoretical explanation.** Here, we explain why, in the real-data sensitivity analysis to misspecification of the isotropic inflation factor, the direct variance component $p$-values do not change, the SAD $p$-values remain close to those seen when using the true isotropic inflation factor, and the direct-SAD covariance $p$-values are larger than those seen when using the true isotropic inflation factor.

We begin with the $p$-value estimation for the direct variance component. Let $\alpha$ denote the true isotropic inflation factor. Suppose that the value used in the second partitioning, $\tilde{\alpha}$ is

$$\tilde{\alpha} = \kappa\alpha \tag{50}$$

for some $\kappa > 0$. By **Eq. 3**, the adjusted sib-GWAS allelic effect is

$$\beta_{Sj} = \alpha\beta^{*}_{Sj},$$

but we are instead using

$$\tilde{\beta}_{Sj} = \tilde{\alpha}\beta^*_{Sj} = \kappa\alpha\beta^*_{Sj} = \kappa\beta_{Sj}, \tag{51}$$

and the corresponding standard errors

$$\tilde{v}_{Sj} = \kappa\hat{v}_{Sj} \tag{52}$$

as inputs. By **Eq. 20**, the direct variance component along PC $i$ is

$$\hat{c}_{Di} = \lambda_i \left( \sum_{j=1}^{\ell} \beta_{Sj} U_{ij} \right)^2 - \lambda_i \sum_{j=1}^{\ell} U_{ij}^2 \hat{v}_{Sj}^2.$$

When the isotropic inflation factor is misestimated (**Eq. 50**), the sib-GWAS allelic effect estimates are affected (**Eqs. 51, 52**). Therefore, the direct variance component is estimated as

$$\begin{aligned}
\tilde{c}_{Di} &= \lambda_i \left( \sum_{j=1}^{\ell} \kappa\beta_{Sj} U_{ij} \right)^2 - \lambda_i \sum_{j=1}^{\ell} U_{ij}^2 \left(\kappa\hat{v}_{Sj}\right)^2 \\
&= \kappa^2\lambda_i \left( \sum_{j=1}^{\ell} \beta_{Sj} U_{ij} \right)^2 - \kappa^2\lambda_i \sum_{j=1}^{\ell} U_{ij}^2 \hat{v}_{Sj}^2 \\
&= \kappa^2 \left[ \lambda_i \left( \sum_{j=1}^{\ell} \beta_{Sj} U_{ij} \right)^2 - \lambda_i \sum_{j=1}^{\ell} U_{ij}^2 \hat{v}_{Sj}^2 \right] \\
&= \kappa^2 \hat{c}_{Di}.
\end{aligned}$$

Similarly, each permutation of allelic effect signs (**Text S3**) leads to a different value of $\hat{c}_{Di}$, as before. But its rescaling to $\tilde{c}_{Di}$ is by the same positive constant. As a result, the relative rank of the observed $\tilde{c}_{Di}$ among permutation-based values is expected to be the same, and thus the $p$-value is expected to be the same.

Similarly, the SAD variance component along PC $i$ is

$$\hat{c}_{\sigma i} = \lambda_i \left( \sum_{j=1}^{\ell} [\beta_{Gj} - \beta_{Sj}] U_{ij} \right)^2 - \lambda_i \sum_{j=1}^{\ell} U_{ij}^2 \hat{v}_{Gj}^2 - \lambda_i \sum_{j=1}^{\ell} U_{ij}^2 \hat{v}_{Sj}^2,$$

and using a misspecified isotropic inflation factor we get

$$\tilde{c}_{\sigma i} = \lambda_i \left( \sum_{j=1}^{\ell} [\beta_{Gj} - \kappa\beta_{Sj}] U_{ij} \right)^2 - \lambda_i \sum_{j=1}^{\ell} U_{ij}^2 \hat{v}_{Gj}^2 - \lambda_i \sum_{j=1}^{\ell} U_{ij}^2 \left(\kappa\hat{v}_{Sj}\right)^2. \tag{53}$$

Expanding the squared term gives

$$\begin{aligned}\tilde{c}_{\sigma i} &= \lambda_i \left(\sum_{j=1}^{\ell} \beta_{Gj} U_{ij}\right)^2 - 2\kappa\lambda_i \left(\sum_{j=1}^{\ell} \beta_{Gj} U_{ij}\right)\left(\sum_{j=1}^{\ell} \beta_{Sj} U_{ij}\right) \\ &+ \kappa^2 \lambda_i \left(\sum_{j=1}^{\ell} \beta_{Sj} U_{ij}\right)^2 - \lambda_i \sum_{j=1}^{\ell} U_{ij}^2 \hat{v}_{Gj}^2 - \kappa^2 \lambda_i \sum_{j=1}^{\ell} U_{ij}^2 \hat{v}_{Sj}^2. \end{aligned} \tag{54}$$

Subtracting the true SAD variance component in **Eq. 18** from the estimated component of **Eq. 54** gives

$$\begin{aligned}\tilde{c}_{\sigma i} - \hat{c}_{\sigma i} &= 2(1-\kappa)\lambda_i \left(\sum_{j=1}^{\ell} \beta_{Gj} U_{ij}\right)\left(\sum_{j=1}^{\ell} \beta_{Sj} U_{ij}\right) \\ &+ (\kappa^2 - 1)\left[\lambda_i \left(\sum_{j=1}^{\ell} \beta_{Sj} U_{ij}\right)^2 - \lambda_i \sum_{j=1}^{\ell} U_{ij}^2 \hat{v}_{Sj}^2\right] \\ &= 2(1-\kappa)\lambda_i \left(\sum_{j=1}^{\ell} \beta_{Gj} U_{ij}\right)\left(\sum_{j=1}^{\ell} \beta_{Sj} U_{ij}\right) + (\kappa^2 - 1)\hat{c}_{Di}. \end{aligned} \tag{55}$$

To express the first term using the variance components, note that by **Eq. 22**, the direct-SAD covariance component is

$$\begin{aligned}\hat{c}_{(D\cdot\sigma)i} &= 2\lambda_i \left(\sum_{j=1}^{\ell} \beta_{Sj} U_{ij}\right)\left(\sum_{j=1}^{\ell} [\beta_{Gj} - \beta_{Sj}] U_{ij}\right) + 2\lambda_i \sum_{j=1}^{\ell} U_{ij}^2 \hat{v}_{Sj}^2 \\ &= 2\lambda_i \left(\sum_{j=1}^{\ell} \beta_{Sj} U_{ij}\right)\left(\sum_{j=1}^{\ell} \beta_{Gj} U_{ij}\right) - 2\lambda_i \left(\sum_{j=1}^{\ell} \beta_{Sj} U_{ij}\right)^2 + 2\lambda_i \sum_{j=1}^{\ell} U_{ij}^2 \hat{v}_{Sj}^2 \\ &= 2\lambda_i \left(\sum_{j=1}^{\ell} \beta_{Sj} U_{ij}\right)\left(\sum_{j=1}^{\ell} \beta_{Gj} U_{ij}\right) - 2\left[\lambda_i \left(\sum_{j=1}^{\ell} \beta_{Sj} U_{ij}\right)^2 - \lambda_i \sum_{j=1}^{\ell} U_{ij}^2 \hat{v}_{Sj}^2\right] \\ &= 2\lambda_i \left(\sum_{j=1}^{\ell} \beta_{Sj} U_{ij}\right)\left(\sum_{j=1}^{\ell} \beta_{Gj} U_{ij}\right) - 2\hat{c}_{Di}. \end{aligned}$$

Therefore,

$$2\lambda_i \left(\sum_{j=1}^{\ell} \beta_{Gj} U_{ij}\right)\left(\sum_{j=1}^{\ell} \beta_{Sj} U_{ij}\right) = \hat{c}_{(D\cdot\sigma)i} + 2\hat{c}_{Di}. \tag{56}$$

Substituting **Eq. 56** into **Eq. 55** gives

$$\begin{aligned}\tilde{c}_{\sigma i} - \hat{c}_{\sigma i} &= (1-\kappa)\left[\hat{c}_{(D\cdot\sigma)i} + 2\hat{c}_{Di}\right] + (\kappa^2 - 1)\hat{c}_{Di} \\ &= (1-\kappa)\hat{c}_{(D\cdot\sigma)i} + \left[2(1-\kappa) + (\kappa^2-1)\right]\hat{c}_{Di} \\ &= (1-\kappa)\hat{c}_{(D\cdot\sigma)i} + (\kappa-1)^2\hat{c}_{Di}.\end{aligned}$$

Thus,

$$\tilde{c}_{\sigma i} = \hat{c}_{\sigma i} + (1-\kappa)\hat{c}_{(D\cdot\sigma)i} + (\kappa-1)^2\hat{c}_{Di}. \tag{57}$$

When the isotropic inflation factor is misestimated, the SAD variance component estimate equals the correctly estimated SAD component plus a first-order direct-SAD covariance component and a second-order direct variance component. The same form of leakage applies to both the observed statistic and the permutation null distribution. As a result, the permutation distribution tends to move together with the observed statistic, so the $p$-value resulting from misspecification often remains close to the value obtained using the true isotropic inflation factor (**Fig. S3B,F**).

The case is different for direct-SAD covariance components. By **Eq. 22**,

$$\hat{c}_{(D\cdot\sigma)i} = 2\lambda_i\left(\sum_{j=1}^{\ell}\beta_{Sj}U_{ij}\right)\left(\sum_{j=1}^{\ell}[\beta_{Gj} - \beta_{Sj}]U_{ij}\right) + 2\lambda_i\sum_{j=1}^{\ell}U_{ij}^2\hat{v}_{Sj}^2.$$

When the isotropic inflation factor is misestimated (**Eq. 50**), the sib-GWAS summaries are affected (**Eqs. 51, 52**). Therefore, the direct-SAD covariance component is misestimated as

$$\tilde{c}_{(D\cdot\sigma)i} = 2\lambda_i\left(\sum_{j=1}^{\ell}\kappa\beta_{Sj}U_{ij}\right)\left(\sum_{j=1}^{\ell}[\beta_{Gj} - \kappa\beta_{Sj}]U_{ij}\right) + 2\lambda_i\sum_{j=1}^{\ell}U_{ij}^2\left(\kappa\hat{v}_{Sj}\right)^2.$$

Expanding this expression gives

$$
\begin{aligned}
\tilde{c}_{(D\cdot\sigma)i} &= 2\kappa\lambda_i\left(\sum_{j=1}^{\ell}\beta_{Sj}U_{ij}\right)\left(\sum_{j=1}^{\ell}\beta_{Gj}U_{ij}\right) - 2\kappa^2\lambda_i\left(\sum_{j=1}^{\ell}\beta_{Sj}U_{ij}\right)^2 + 2\kappa^2\lambda_i\sum_{j=1}^{\ell}U_{ij}^2\hat{v}_{Sj}^2 \\
&= \kappa\left[2\lambda_i\left(\sum_{j=1}^{\ell}\beta_{Sj}U_{ij}\right)\left(\sum_{j=1}^{\ell}\beta_{Gj}U_{ij}\right)\right] - 2\kappa^2\left[\lambda_i\left(\sum_{j=1}^{\ell}\beta_{Sj}U_{ij}\right)^2 - \lambda_i\sum_{j=1}^{\ell}U_{ij}^2\hat{v}_{Sj}^2\right] \\
&= \kappa\left[\hat{c}_{(D\cdot\sigma)i} + 2\hat{c}_{Di}\right] - 2\kappa^2\hat{c}_{Di} \\
&= \kappa\hat{c}_{(D\cdot\sigma)i} + 2\kappa(1-\kappa)\hat{c}_{Di} \\
&= \kappa\left[\hat{c}_{(D\cdot\sigma)i} + 2(1-\kappa)\hat{c}_{Di}\right].
\end{aligned}
$$

The direct-SAD covariance component is not only rescaled by $\kappa$, but also receives an additional contribution from the direct variance component. This extra term can increase the spread of the permutation null distribution, especially when the direct component is large. Consequently, the $p$-values for covariance terms can become larger under misestimation of the isotropic inflation factor, consistent with what we observe in the real-data sensitivity analysis (**Fig. S3C,G**).

#### S14 Simulation study

We designed a simulation study to evaluate the performance of PGSUS under a range of conditions. We first established a baseline model: a single, randomly mating population with direct genetic effects and independent environmental effects. We then introduced a series of modifications to this scenario to explore their effects on PGSUS outputs. These extended scenarios include population stratification arising from genetic drift and environmental divergence, assortative mating on the basis of phenotypic similarity, dynastic effects in which parental genotypes influence offspring phenotype regardless of their transmission, environmental effects correlated with genetic relatedness, biased sampling of individuals, and scenarios exemplifying possible effects of SAD variance on PGS portability.

##### S14.1 Baseline simulation procedures

The simulation pipeline consists of three stages: first, a library of linked marker SNP-causal SNP pairs was constructed. Second, marker-causal variant pairs were randomly and independently drawn from the library to assemble into genomes for individuals in the parental population, which was split into three cohorts. Third, individuals within each cohort were mated to produce offspring (**Fig. S40**).

**Marker-causal variant library.** We simulated the coalescent history of a 500 kb genomic region with *msprime*[51]. This window size was selected to capture the typical physical distances between marker and causal variants in humans[52]. We assumed an effective population size ($N_e$) of 10,000, a mutation rate of $1 \times 10^{-8}$ per base pair per generation, and a recombination rate of $1 \times 10^{-8}$ per base pair per generation. To ensure diverse genealogical histories, we performed 500 independent replicates using distinct random seeds. Within each replicate, we enumerated every pair of SNPs in the genomic region and recorded both the physical distance between the two variants and the four observed haplotype frequencies ($f_{00}$, $f_{01}$, $f_{10}$, $f_{11}$). Here we focus on common variants, so we restricted the library to SNPs with a minor allele frequency greater than 0.01. To ensure strong LD between marker and causal SNPs, we only retained pairs with an $r^2 \geq 0.5$.

**Parental population generation.** To create a population of 15,000 diploid individuals, we generated 30,000 independent haploid "genomes." Each haploid genome carried 1,000 marker-causal pairs randomly drawn from the library. Every draw of a marker-causal pair of alleles was independent of all other such draws. For each pair, the haplotype state (00, 01, 10, or 11) was sampled according to the frequencies $f_{00}$, $f_{01}$, $f_{10}$, $f_{11}$. We then randomly paired these 30,000 chromosomes to form the genomes of 15,000 individuals. The parental population was partitioned into three groups (5,000 individuals per group) to generate offspring for downstream analysis: a standard-GWAS cohort (one offspring per mating pair), a sibling GWAS cohort (two offspring per mating pair), and a prediction cohort (one offspring per mating pair).

**Offspring generation and phenotype simulation.** The 5,000 individuals within each cohort were randomly paired to form 2,500 mating pairs. Offspring populations were generated by simulating meiosis in these mating pairs. To construct each offspring haplotype, the simulation traversed the parental marker and causal variants. The probability of a crossover event occurring between a marker and its paired causal variant was calculated by multiplying their physical distance (bp) by a uniform recombination rate of $1 \times 10^{-8}$ per base pair. After simulating recombination, one of the two parental haplotypes was passed to an offspring, with probability $\frac{1}{2}$ each. The offspring's other haplotype was sampled from the other parent in a similar manner. Phenotypes ($y$) were modeled as the sum of a genetic component ($G$) and an independent environmental component ($\epsilon$) such that $y = G + \epsilon$. The genetic component was calculated as the weighted sum of alleles carried: $G = \sum_{i=1}^{n} \beta_i X_i$, where $\beta_i$ is the true allelic effect of causal variant $i$ drawn from a standard Normal distribution ($\beta_i \sim \mathcal{N}(0,1)$), and $X_i$ denotes the allele dosage ($X_i \in \{0,1,2\}$). The environmental component was drawn independently from a Normal distribution, $\epsilon \sim \mathcal{N}(0, V_G)$ where the variance was set to equal the empirical variance of the genetic component across all individuals in the cohort, $V_G$, such that genetic and environmental variance contributed roughly equally.

##### S14.2 Performance of PGSUS under baseline (null) conditions

We first evaluated PGSUS's performance under the baseline model—that is, with random mating, no population structure, no indirect effects, and no ascertainment of SNPs. Since no SAD effects were introduced, we expected to see few significant SAD loadings on any PCs, with the type I error rate remaining near the nominal level (0.05). Additionally, we expected estimated isotropic inflation factors close to one and non-direct variance components near zero.

Consistent with our expectations, across 100 replicates, PC-specific SAD components were near zero, and the great majority of estimated variance components fall within the empirical 95% null non-rejection region generated from permutations. The false positive rates (proportion of replicates with $p$-value $< 0.05$) for both direct and SAD-related variance components are close to the nominal rate (**Fig. S41A**), suggesting that PGSUS does not falsely attribute variance to SAD effects under this null model. In addition, the estimated isotropic inflation factors from 100 replicates are centered near 1 (mean = 1.02, standard error = 0.018) (**Fig. S41B**), and the estimated non-direct variance components are centered near zero (mean = -0.016, standard deviation = 0.099) (**Fig. S41C**).

##### S14.3 Effects of SNP ascertainment on estimated isotropic inflation factor and non-direct variance component under baseline conditions

In realistic applications, SNPs will be incorporated into a PGS on the basis of a small $p$-value for the null hypothesis of no association, or some statistic closely related to the $p$-value. We examined the effect of such SNP ascertainment on PGSUS outputs under the baseline model. First, we ascertained the $m$ most significantly associated marker SNPs from the standard GWAS ($m \in \{200, 500, 800\}$) as index SNPs included in the PGS. We expected the standard-GWAS allelic effects estimates' magnitude to be upwardly biased as a result of the winner's curse[48,49]. The bias should be more pronounced under more stringent ascertainment thresholds, leading to an isotropic inflation factor greater than one and a non-direct variance component greater than zero.

In addition, we considered another case in which the most significant SNPs in the sib-GWAS were ascertained for inclusion in the PGS. In this case, we expected winners' curse to be reversed and ascertainment to be reflected in isotropic deflation.

These predictions are borne out in the results. When SNPs are ascertained on the basis of standard-GWAS $p$-values, isotropic inflation factor estimates are generally higher than one (**Fig. S19A**). The degree of inflation is greater when ascertainment is more stringent. Conversely, when SNPs are ascertained on the basis of sib-GWAS $p$-values, it produces the opposite effect: isotropic inflation factors are consistently below one, with stricter thresholds leading to lower values (**Fig. S19B**). Similarly, more stringent ascertainment on the basis of standard-GWAS $p$-values leads to larger estimated non-direct variance components (**Fig. S19C**), and more stringent ascertainment on the basis of sib-GWAS $p$-values leads to smaller estimated non-direct variance components (**Fig. S19D**).

##### S14.4 Simulations of population stratification

One of the main intended uses of PGSUS is to study the effects of population stratification on the application of a PGS. Under neutral evolution, populations diverging from a common ancestor accumulate phenotypic differences over time as genetic drift leads to allele-frequency divergence between isolated subpopulations. The expected magnitude of phenotypic difference scales with split time and environmental difference[53–55]. We thus systematically evaluated two sources of stratification: genetic drift, represented by the split time between subpopulations; and environmental stratification, represented by a shift in the mean of environmental component distribution.

To simulate population stratification, we modified marker-causal variant library construction by specifying a two-population demographic model in *msprime*[51]. In this model, a single ancestral population of 20,000 individuals split into two subpopulations (A and B) at a fixed number of generations ago $t$. Each subpopulation evolved with a constant size of 10,000 individuals. In different simulations, we used $t \in \{100, 500, 1000\}$. Each cohort (standard-GWAS, sib-GWAS, and prediction) consisted of 5,000 individuals (2,500 mating pairs), with 2,500 drawn from each subpopulation. Within each subpopulation, individuals were randomly paired for mating. In some simulations, the environmental effects did not differ on average between populations. In others, we added an additional component of environmental divergence to two populations diverged t=500 generations ago. The environmental component was drawn from $\mathcal{N}(-\delta\sqrt{V_G}, 1)$ for population A and $\mathcal{N}(\delta\sqrt{V_G}, 1)$ for population B, where $V_G$ represents the empirical variance of the genetic component across all individuals in the cohort, and $\delta \in \{0, 0.2, 0.6, 1\}$ controls the extent of environmental divergence between subpopulations. In some cases, we ran GWAS without an adjustment for population structure. In others, we adjusted either for the true binary subpopulation identity (A or B) or for individual coordinates on the first principal component of the genotype matrix as a GWAS covariate (via the `--covar` flag in PLINK[19]). All other simulation settings were identical to the baseline scenario.

When standard GWAS and sib-GWAS were conducted with these stratified samples without adjustment, we observe significantly elevated SAD variance and non-direct variance along PC1 (**Fig. S7A-F**). This is not surprising, as PC1 captures the primary axis of genetic difference between two groups[56]. In many iterations, both estimated SAD variance and non-direct variance fall outside the null non-rejection region. We also noticed that although the rejection rate is elevated with uncorrected structure, the relationships between rejection rate on one hand and environmental divergence and split time on the other hand is not monotonic. This is possibly caused in part by an expansion of the null non-rejection region. Another non-exclusive explanation relates to the fact that the mean difference in the genetic component evolves neutrally. As such, the direction of mean genetic difference between subpopulations will oppose the direction of the environmental difference half the time, partially canceling the total phenotypic difference between populations when the genetic and environmental effects are of similar magnitude.

We also examined the isotropic inflation factor and non-direct variance component (**Fig. S7G-J**). The estimated isotropic inflation factor tends to increase as both split time and environmental divergence grow (**Fig. S7G,I**). Adjusting for population stratification by including either population label or individual PC1 coordinates as covariates in the regression brought PC-specific SAD variance near zero and the average estimated isotropic inflation factor close to one. Similarly, the estimated non-direct variance component is large when population structure is not adjusted for and near zero when it is adjusted (**Fig. S7H,J**).

##### S14.5 Simulations of assortative mating

We next examined the impact that assortative mating on phenotypic value had on PGSUS outputs. The variant library and parental population were generated just as in the baseline model. To model assortative mating, individuals in the parental population were paired on the basis of phenotypic similarity to achieve a predefined spousal correlation—the Pearson correlation coefficient between mates' phenotypes. The matching algorithm takes the array of parental phenotypes and reorganizes the individuals into pairs such that the Pearson correlation coefficient between the two partners across all pairs approximates target spousal correlation. This is implemented by rank-matching the simulated phenotypes to a target bivariate Normal distribution generated with the desired correlation. The algorithm, originally developed for two correlated traits simultaneously[57], was simplified to the single-trait case here and returns a set of re-ordered indices indicating the matings. We evaluated three scenarios by varying spousal correlation, $\rho \in \{0.2, 0.5, 0.8\}$.

Under positive assortative mating, standard-GWAS allelic effect estimates are expected to be biased to be larger in magnitude as if multiplied by a constant factor. In our simulation framework, we expected this to translate to an increase in the estimated isotropic inflation factor without systematically elevating SAD variance estimates along any specific PC. Consistent with the expectation, we observed that the isotropic inflation factor values increased with higher spousal correlation (**Fig. S17A**). Higher spousal correlations also increased the estimated non-direct variance component (**Fig. S17B**). We do not observe any elevated SAD variance on specific PCs (**Fig. S17C**).

##### S14.6 Simulations of dynastic effects

To simulate dynastic effects, we augmented the phenotype values resulting from the baseline generative model. Given the causal SNP's direct genetic effect, $\beta_{direct}$, we assigned its corresponding marker variant a dynastic allelic effect, $\beta_{dynastic} = \rho \cdot \beta_{direct} + \sqrt{1-\rho^2} \cdot \eta$ ($\eta \sim \mathcal{N}(0,1)$), where $\rho$ controls the correlation between direct and dynastic effects. The contribution of dynastic effect to the offspring phenotype was calculated as the weighted sum of the average of the parents' allele dosages multiplied by $\beta_{dynastic}$. The total offspring phenotype was then reset to be $y = \beta_{\mathrm{direct}} G_{\mathrm{offspring}} + \beta_{\mathrm{dynastic}} G_{\mathrm{avg(parent)}} + \epsilon$, where $G_{\mathrm{offspring}}$ is the offspring genotype and $G_{\mathrm{avg(parent)}}$ is the average parents' genotypes. All other simulation settings were identical to the baseline model.

To evaluate how dynastic effects affect PGSUS outputs, we studied three cases. Specifically, we simulated dynastic allelic effects negatively correlated ($\rho = -0.6$), uncorrelated ($\rho = 0$), or positively correlated ($\rho = 0.6$) with direct allelic effects. In an unstructured population, none of these cases led to elevated PC-specific variance components (across the top six PCs, all variance components were inside the null non-rejection region at least 91% of the time). However, the estimated isotropic inflation factor was influenced by dynastic effects, becoming larger than one if the dynastic and direct effects were positively correlated and smaller than one when they were negatively correlated (**Fig. S18A**). The estimated non-direct variance component responded similarly, becoming larger than 0 when dynastic and direct effects were positively correlated and smaller than 0 when dynastic and direct effects were negatively correlated (**Fig. S18B**).

##### S14.7 Simulations of confounding by an environmental component proportional to relatedness.

The simulations in the previous sections focus on the core targets of PGSUS, examining how PGSUS responds in the presence (or absence) of stratification, assortative mating, and dynastic effects. However, there may be other ways in which GWAS-estimated allelic effects could be biased that do not fall neatly into this framework. As one example, we simulated a structured environment in which environmental similarity correlates with genetic relatedness. We used the same two-population demographic model as in **Text S14.4** with a split time of $t = 1,000$ generations. We drew the environmental component from a multivariate normal distribution $\mathcal{N}(0, \lambda^2 \mathbf{G})$, where $\mathbf{G}$ is a genetic relatedness matrix (GRM) calculated from all SNPs across individuals in the GWAS and sib-GWAS samples. Specifically, let $\mathbf{X}$ be an $n \times m$ genotype matrix for $n$ individuals and $m$ SNPs, where genotypes are coded as {0, 1, 2} per the number of effect alleles. $\mathbf{G}$ was calculated following the method of VanRaden[58]: $\mathbf{G} = \frac{ZZ^T}{\sum_{j=1}^{m} 2p_j(1-p_j)}$ where $Z$ is the centered genotype matrix and $p_j$ is the observed allele frequency of the $j$-th SNP. The scaling factor $\lambda$ controls the magnitude of environmental effects. We evaluated $\lambda \in \{1, 3, 5\}$.

Under this model, we found that the effect of the scaling factor on the isotropic inflation factor estimate depended on the alignment of genetic and environmental effects. When the environmental difference between the two populations aligns with the direction of genetic difference, the estimated isotropic inflation factor increases monotonically with the environmental scaling factor (**Fig. S42A**). In contrast, when the environmental difference opposed the genetic difference, increasing the scaling factor diminished the magnitude of the estimated isotropic inflation factor (**Fig. S42B**). Notably, when we included an adjustment for PC1 in the standard GWAS, the isotropic inflation estimates centered approximately at one (**Fig. S42C**).

##### S14.8 Simulations of biased sampling of individuals by ancestry

We also examined the effects of non-uniform sampling of individuals. To simulate biased sampling by ancestry, we used the same two-population demographic model as in **Text S14.4** with a split time of $t = 1,000$ generations without environmental difference. The cohort composition was modified to introduce ancestry-biased sampling: the standard-GWAS sample and prediction sample comprised 1,250 individuals from each of populations A and B (2,500 total), and the sib-GWAS sample was restricted to 2,500 individuals drawn exclusively from population A.

Since standard GWAS captures both the direct genetic effect and correlation between genotype and phenotype due to stratification, we anticipated this sampling strategy should lead to inflation in standard-GWAS estimates. At the variant level, we observed mismatches between standard and sib-GWAS samples in allele frequencies, local LD, and estimated allelic effects (**Fig. S43A-C**). Our simulation yielded an average value of 3.52 (standard error = 0.25) for the estimated isotropic inflation factor across 100 iterations. After including population labels as standard-GWAS covariates, the average value drops to 1.01 (standard error = 0.012). However, we still observed elevated PC1-SAD variance driven by population structure (**Fig. S43D**). Adjusting for population structure led to a reduction in PC1 variance components and their rejection rate, but they remained slightly elevated above baseline levels (**Fig. S43E**), suggesting that the effect is not fully explained by population stratification biasing GWAS effect estimates.

##### S14.9 Simulations of phenotype-based ascertainment

In addition to biased sampling by ancestry, we considered biased sampling into the sib-GWAS by phenotype. Here, we considered phenotypes affected by both direct and SAD effects—specifically, dynastic effects—and participation bias into the sib-GWAS sample. We used the simulated data generated in **Text S14.6** (with the correlation between direct and dynastic effects, $\rho = 0$). Within the sib-GWAS samples, we retained a fraction (20% and 50%) of sibling pairs with the highest sib-averaged phenotype value. Because the top 20% subset was the smallest, we randomly downsampled the samples containing all individuals or the top 50% of individuals to the same number of sibling pairs as were present in the sample of siblings with the top 20% of phenotype values. Sib-GWAS estimation was then rerun on each set of ascertained sibling pairs.

We assessed whether this phenotype-based ascertainment changed the relationship between estimated direct effects and simulated dynastic effects by calculating their correlation across ascertainment levels. The results indicate that the ascertainment of phenotypic extremes does not systematically change the magnitude of the correlation, the estimated isotropic inflation factor, or the estimated non-direct variance component (**Fig. S24**). However, it did seem to alter the distribution of the estimated isotropic inflation factor and the estimated non-direct variance component (**Fig S24B,C**), increasing their spread.

##### S14.10 A cautionary tale regarding PGS portability

The relationship between SAD variance in a PGS and its portability is not straightforward. Importantly, it depends on whether the SAD effects lead standard-GWAS allelic effect estimates to be predictive in the set of individuals of interest. To illustrate this, we simulated a three-population split model in which two closely related populations experience very different environments.

Specifically, a single ancestral population first split into population A and the common ancestor of population B and C 500 generations ago; populations B and C then diverged from each other 50 generations ago (**Fig. S14A**). Each subpopulation had a constant size of 10,000 individuals. In contrast to the baseline model that incorporated 1,000 pairs of SNPs, we restricted this simulation to 100 pairs of SNPs, drawing true direct allelic effects from $\mathcal{N}(0, 0.1^2)$. Environmental divergence across subpopulations was introduced by drawing environmental components for individuals in population A from $\mathcal{N}(0, 1)$, for individuals in population B from $\mathcal{N}(\sqrt{V_G}, 1)$, and for individuals in population C from $\mathcal{N}(-\sqrt{V_G}, 1)$, where $V_G$ represents the empirical variance of the genetic component calculated across all individuals in the cohort. Standard GWAS were conducted on a training sample of 2,000 individuals, with 1,000 individuals each from populations A and B, both with and without population-label adjustment as a covariate. The standard-GWAS allelic effect estimates were then applied to two independent test sets of 2,000 individuals each: one matching the training sample composition (1,000 each from populations A and B) and one comprising individuals from populations A and C (1,000 from each, **Fig. S14A**). We evaluated prediction accuracy as the Pearson correlation between the phenotype value and the PGS in the test set.

When the test sample had the same population composition as the training GWAS samples (populations A and B), standard-GWAS adjustment for population structure decreased PGS prediction accuracy (**Fig. S14B**). But when the composition of the test set (A and C) differed from that of the training set (A and B), population-label adjustment increased prediction accuracy (**Fig. S14B**). Thus, SAD variance in the GWAS allelic effect estimates – present when population labels are not adjusted for and absent otherwise — helps prediction in one scenario we considered here and hurts prediction in another. This mirrors the results of our empirical study of predictive accuracy in the presence of SAD effects (**Fig. 5**).

#### S15 The effects of index SNP ascertainment

It is typical to select SNPs for inclusion in a PGS on the basis of the allelic effects estimated in standard GWAS (or functions of them, such as $p$-values). This process affects the interpretation of PGSUS outputs in two ways. First, it changes the estimands of PGSUS—PGSUS aims to decompose variance not in a trait but rather in a specific PGS in a specific prediction sample, so changing the PGS by ascertaining different SNPs changes the estimands. Second, ascertainment can, in some cases, lead to violations of some of the assumptions employed in deriving the PGSUS estimators, which can cause bias. Here, we discuss how SNP ascertainment in standard GWAS can a) change the interpretation of an estimated quantity even in the absence of bias, and b) cause bias in PC-specific estimated SAD and covariance terms.

##### S15.1 SNP ascertainment as a collider

In the majority of cases we examined, the presence of significant direct variance components were accompanied by large (most often also significant) SAD and/or non-direct variance components. We highlighted several such examples in **Fig. S27**. We hypothesized that the observed significant direct components may be partly or fully due to an artifact driven by the combination of substantial SAD variance at the population level—possibly in the absence of natural selection—and index SNP ascertainment.

For example, consider a two-population demographic model in which subpopulations differ in allele frequency and stratification is not adjusted (or incompletely adjusted). In this case, standard-GWAS allelic effect estimates are inherently confounded by stratification. Specifically, alleles at higher frequencies in the subpopulation with a larger mean phenotype will exhibit upwardly biased allelic effect estimates. If this stratification is not negligible and loci are ascertained by their absolute allelic effect, the selection process favors variants where the true direct effect and the stratification bias directionally align. Consequently, even if sibling-based allelic effect estimates are unbiased at individual loci, a sibling-based PGS computed from these GWAS-ascertained SNPs will be biased toward alignment with the axis of stratification. This phenomenon arises from collider bias: the ascertainment of index SNPs induces a correlation between the direct effects and the stratification effects. Zaidi and Mathieson[59] observe this phenomenon (see their Figure 6), and Blanc and Berg[60] also consider this process in detail (see their supplementary text section S8), though neither reference discusses the process in terms of collider bias.

This mechanism suggests that ascertainment based on GWAS allelic effect estimates can generate a large direct-effect variance component along a given PC, provided there is substantial SAD variance along that same PC.

To show that this intuition holds, we used the genetic data simulated under the two-population demographic model described in **Text S14.4** ($t = 100$ generations, $\delta = 0.6$), in which the two subpopulations differ in allele frequencies due to neutral genetic drift and in mean phenotype due to an environmental shift, with no difference in direct genetic effects between populations. We ascertained SNPs across varying GWAS ascertainment $p$-value thresholds (1, 0.3, and 0.03). We observed that as the threshold becomes more stringent, a positive correlation between the PC1-direct variance and PC1-allelic effect variance components emerges across ascertained levels (**Fig. S28A-C**). Correspondingly, the permutation-based rejection rate for the PC1-specific direct variance components increases with a stringent threshold, while the rejection rate for SAD variance components decreases (**Fig. S28D**). One consequence is that a significantly large PC-specific direct variance component should not be taken as evidence of natural selection when it is accompanied by a large SAD variance component on the same PC.

##### S15.2 SNP ascertainment can bias PC-specific SAD and covariance component estimators

In the main text (**Eqs. 14-22**), we derive estimators based on a partitioning of PGS variance in a prediction sample into four components per PC: direct variance, SAD variance, direct-SAD covariance, and error variance. These components arise because we model standard-GWAS-estimated allelic effects as arising from a direct effect, a SAD effect, and estimation error. However, the full decomposition (adapting **Eq. 21**) is

$$\lambda_i \left( \sum_{j=1}^{\ell} \beta_{Gj} U_{ij} \right)^2 = \lambda_i \left( \sum_{j=1}^{\ell} \beta_{Dj} U_{ij} \right)^2 + \lambda_i \left( \sum_{j=1}^{\ell} \sigma_j U_{ij} \right)^2 + 2\lambda_i \left( \sum_{j=1}^{\ell} \beta_{Dj} U_{ij} \right) \left( \sum_{j=1}^{\ell} \sigma_j U_{ij} \right) + \lambda_i \left( \sum_{j=1}^{\ell} \epsilon_{Gj} U_{ij} \right)^2 + 2\lambda_i \left( \sum_{j=1}^{\ell} \beta_{Dj} U_{ij} \right) \left( \sum_{j=1}^{\ell} \epsilon_{Gj} U_{ij} \right) + 2\lambda_i \left( \sum_{j=1}^{\ell} \sigma_j U_{ij} \right) \left( \sum_{j=1}^{\ell} \epsilon_{Gj} U_{ij} \right). \quad (58)$$

The first four terms are, respectively, the direct component ($c_{Di}$), the SAD component ($c_{\sigma i}$), the direct-SAD covariance component ($c_{(D\cdot\sigma)i}$), and the error component. The subsequent coterms are structurally analogous to $c_{(D\cdot\sigma)i}$ and can be labeled a direct-error covariance term and a SAD-error covariance term. Although these final two terms are always present in the PGS variance along a given PC, in the main text, they are assumed to be zero in expectation because $\mathrm{E}[\epsilon_{G_j}] = \mathrm{E}[\epsilon_{S_j}] = 0$ for all $j$. This assumption is standard and sensible for estimation errors in general, but it may not always apply to ascertained SNPs.

We consider a heuristic approximation of the SNP ascertainment approach: we specify whether a given SNP will be included in a PGS by whether the magnitude of its allelic effect estimate $(\beta_D + \sigma + \epsilon_G)$ is greater than some cutoff. (In practice, the cutoff would depend on allele frequency.) For SNPs whose values of $|\beta_D + \sigma|$ are near the cutoff, their inclusion can depend on the size and sign of $\epsilon_G$, which can lead to a positive correlation between the error term and the sum $\beta_D + \sigma$. If, in turn, $\beta_D + \sigma$ is correlated with SNP loadings on principal component $i$—e.g., due to stratification along that axis—then ascertainment may induce a correlation between SNP PC loadings and estimation errors among selected SNPs, potentially invalidating the assumption that $\mathrm{E}[\sum_{j=1}^{\ell} \epsilon_{Gj} U_{ij}] = 0$ among ascertained SNPs, even if it holds when considering a uniformly randomly chosen set of SNPs.

Along similar lines, in the main text, we assume that estimation errors are isotropic in PC space (i.e. that errors are not systematically correlated with loadings on any one PC). This assumption (along with uncorrelated errors among loci) lets us write the expectation of the error component as $\mathrm{E}[(\sum_{j=1}^{\ell} \epsilon_{Gj} U_{ij})^2] = \sum_{j=1}^{\ell} U_{ij}^2 v_{Gj}^2$ where $v_{Gj}^2$ represents the SNP-level variance of the standard-GWAS estimation error. Thus, subtracting an estimate of this expectation (plugging in estimates of $v_{Gj}^2$) allows one to estimate the error variance along a PC and isolate it from the other variance components. But under ascertainment, the isotropy assumption for estimation errors may not hold—there may be systematic correlations between errors and PC loadings along certain PCs, as discussed above.

Thus, motivated by the setting of ascertainment in GWAS, we might ask the expectation of the estimators of PC-specific variance components if we relax assumptions about standard-GWAS error projections along a PC of interest. (Because we are imagining ascertainment in standard GWAS, we will assume that sib-GWAS errors behave as in the main text, as well as retaining the assumption that sib-GWAS estimation errors are independent of standard-GWAS estimation errors.)

First, the estimated direct component $\hat{c}_{Di}$ does not depend on standard-GWAS estimation errors (**Eq. 20**), so it remains approximately unbiased provided that the estimates of sib-GWAS estimation error variance are nearly accurate. To emphasize, this lack of bias is with respect to the PGS—as discussed above, the estimands of PGSUS are variance components of PGSs, not of the traits on which they are based.

In contrast, the estimated SAD component has expectation

$$\mathrm{E}[\hat{c}_{\sigma i}] = c_{\sigma i} + 2\lambda_i \left( \sum_{j=1}^{\ell} \sigma_j U_{ij} \right) \mathrm{E}\left[ \sum_{j=1}^{\ell} \epsilon_{Gj} U_{ij} \right] + \lambda_i \mathrm{E}\left[ \left( \sum_{j=1}^{\ell} \epsilon_{Gj} U_{ij} \right)^2 - \sum_{j=1}^{\ell} U_{ij}^2 \hat{v}_{Gj}^2 \right]. \tag{59}$$

There are two bias terms here—any SAD-error covariance caused by ascertainment is lumped into the estimated SAD term, and any systematic difference between the true and estimated error variance projection on PC also biases the estimated SAD term.

The estimated direct-SAD covariance component has expectation

$$\mathrm{E}[\hat{c}_{(D\cdot\sigma)i}] = c_{(D\cdot\sigma)i} + 2\lambda_i \left( \sum_{j=1}^{\ell} \beta_{Dj} U_{ij} \right) \mathrm{E}\left[ \sum_{j=1}^{\ell} \epsilon_{Gj} U_{ij} \right]. \tag{60}$$

In words, the estimated direct-SAD covariance term absorbs the direct-error covariance and is biased by its expectation for any PCs on which that expectation is nonzero due to ascertainment.

The biasing terms in **Eqs. 59, 60** can in principle be either positive or negative. Ignoring the misestimation of the error component (the final square-bracketed statement in **Eq. 59**), the remaining bias terms share $\mathrm{E}\left[\sum_{j=1}^{\ell} \epsilon_{Gj} U_{ij}\right]$. Thus, these contributions to variance will be of the same sign when $\left(\sum_{j=1}^{\ell} \sigma_j U_{ij}\right)$ and $\left(\sum_{j=1}^{\ell} \beta_{Dj} U_{ij}\right)$ are of the same sign, which is to say when the true direct-SAD covariance term is positive. The biasing terms are of opposite signs when $\left(\sum_{j=1}^{\ell} \sigma_j U_{ij}\right)$ and $\left(\sum_{j=1}^{\ell} \beta_{Dj} U_{ij}\right)$ are of opposite signs, meaning that the true direct-SAD covariance term is negative. This observation may partially explain empirical examples in which large, PC-specific estimated SAD variance components are accompanied by large negative estimated covariance components. Some of these may be cases in which the true direct-SAD covariance components are negative, and SNP ascertainment on a PC with a large SAD components drives the estimated direct-SAD covariance component to be negative with an even larger magnitude.

### Supplemental Figures

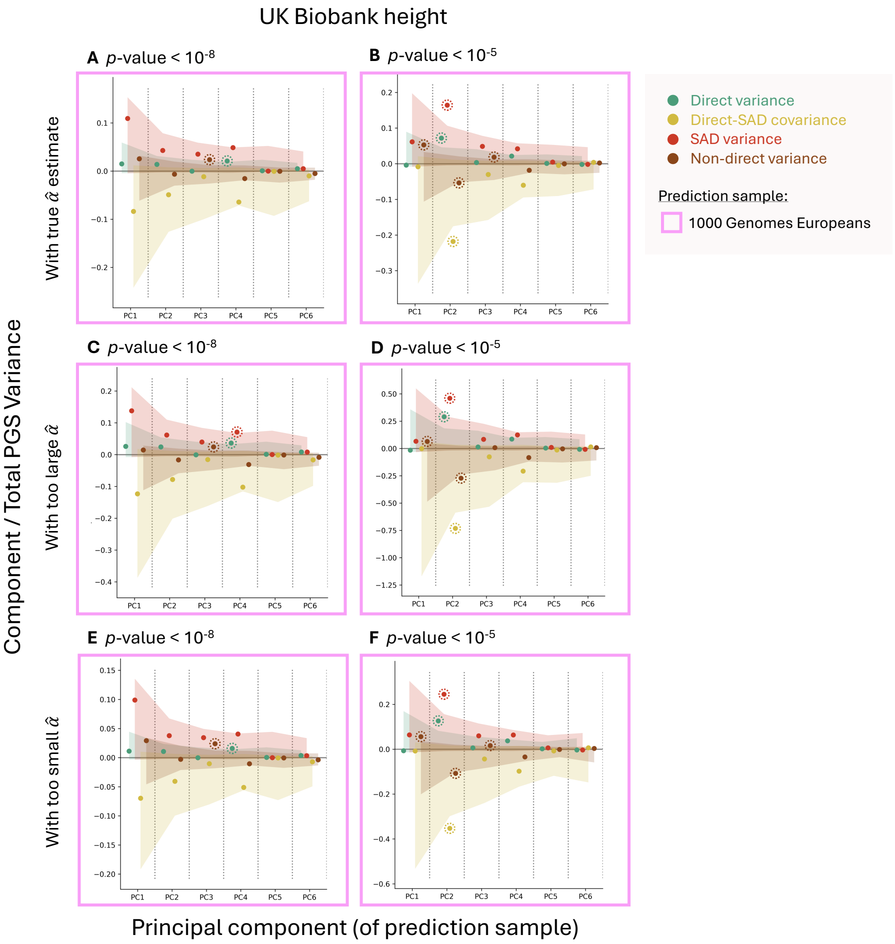

**Figure S1. Variance partitionings are largely insensitive to misspecification of the isotropic inflation factor.** PGSUS partitions the variance of a PGS among individuals in a prediction sample. Variance components are attributable to an effect type—direct effects, SAD effects and their covariance—and a principal component of the genotype matrix of the prediction sample. Shown are variance components (divided by the total variance in the PGS) along the first six PCs for PGSs constructed for the UK Biobank standard-GWAS summary statistics for height at two different GWAS ascertainment thresholds ($p$-value $< 10^{-8}$ and $p$-value $< 10^{-5}$). To test the effect of misspecification of the isotropic inflation factor, we adjusted the sib-GWAS allelic effect estimates and standard errors by an isotropic inflation factor which is larger or smaller than the true estimate ($\hat{\alpha}$)—$\alpha := 1.314 \times \hat{\alpha}$ or $\alpha := 0.867 \times \hat{\alpha}$, respectively—prior to performing variance partitioning. Partitionings in panels **(A)** and **(B)** were performed with the true isotropic inflation factor, while panels **(C)** and **(D)** used the larger isotropic inflation factor and panels **(E)** and **(F)** used the smaller isotropic inflation factor. Variance component estimates and their significance were mostly unaffected.

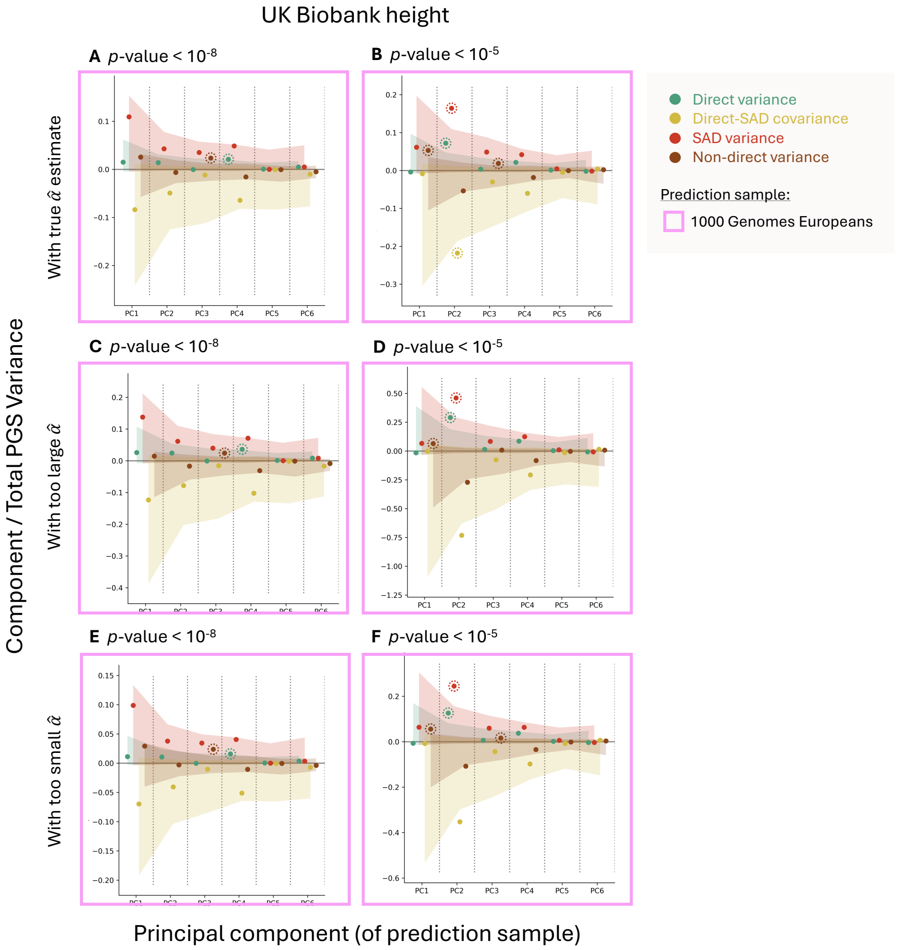

**Figure S2. Variance partitionings, tested for significance using the adaptive $p$-value procedure, are largely insensitive to misestimation of the isotropic inflation factor.** PGSUS partitions the variance of a PGS among individuals in a prediction sample. Variance components are attributable to an effect type—direct effects, SAD effects and their covariance—and a principal component of the genotype matrix of the prediction sample. Shown are variance components (divided by the total variance in the PGS) along the first six PCs for PGSs constructed for the UK Biobank standard-GWAS summary statistics for height at two different GWAS ascertainment thresholds ($p$-value < $10^{-8}$ and $p$-value < $10^{-5}$). To test the effect of misspecification of the isotropic inflation factor, we adjusted the sib-GWAS allelic effect estimates and standard errors by an isotropic inflation factor which is larger or smaller than the true estimate ($\hat{\alpha}$)—$\alpha := 1.314 \times \hat{\alpha}$ or $\alpha := 0.867 \times \hat{\alpha}$, respectively—prior to performing variance partitioning. Partitionings in panels **(A)** and **(B)** were performed with the true isotropic inflation factor, while panels **(C)** and **(D)** used the larger isotropic inflation factor and panels **(E)** and **(F)** used the smaller isotropic inflation factor. Variance component estimates and their significance were mostly unaffected. This figure is the same as **Fig. S1**, but $p$-values were estimated using the adaptive $p$-value estimation procedure outlined in **Text S4**.

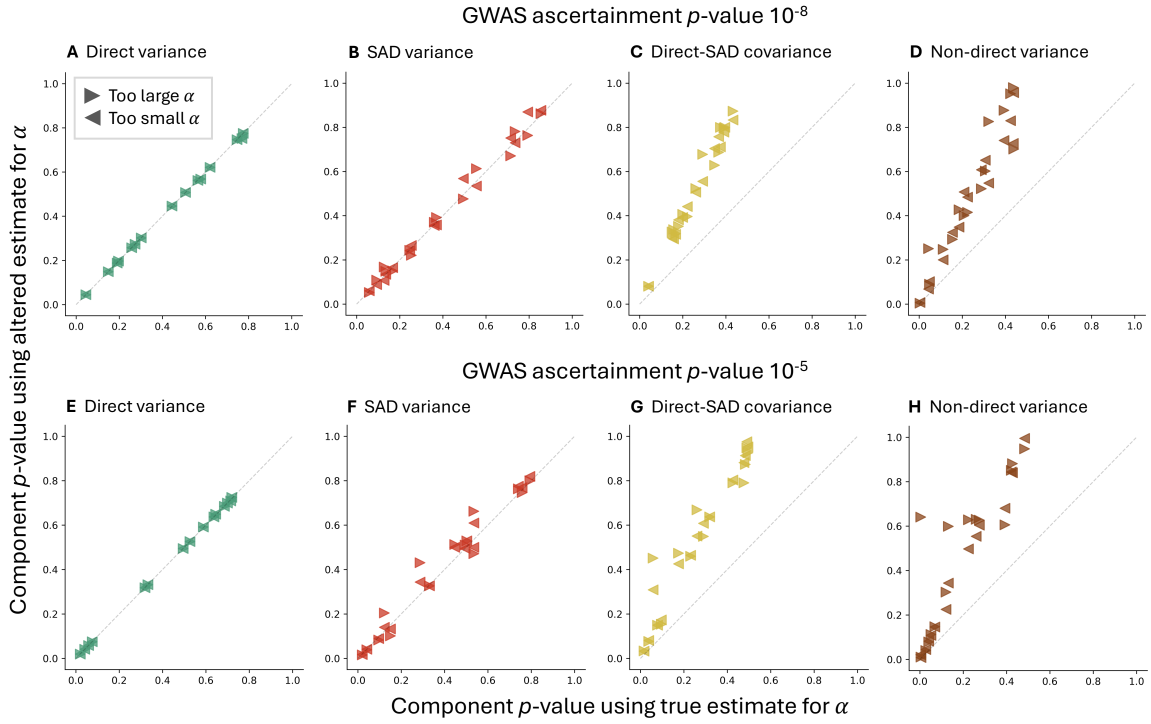

**Figure S3. Misestimation of the isotropic inflation factor results in equal or larger $p$-value estimates for variance components.** PGSUS partitions the variance of a PGS among individuals in a prediction sample. Variance components are attributable to an effect type—direct effects, SAD effects and their covariance—and a principal component of the genotype matrix of the prediction sample. For PGSs of height using the UKB White British GWAS summaries at two GWAS ascertainment thresholds, we tested the effect of misestimation of the isotropic inflation factor ($\alpha$) on $p$-values for partitioned variance components. Shown are the variance component $p$-values when the true estimate of the isotropic inflation factor is artificially increased or decreased to result in an estimate which is too large (right-facing triangles) or too small (left-facing triangles) as compared to the true estimate. Dashed gray lines indicate variance component $p$-values where the altered isotropic inflation estimate is the same as the true estimate. Direct variance component $p$-values are unchanged **(A,E)** and SAD variance component $p$-values are mostly unaffected **(B,F)**; in contrast, $p$-values for direct-SAD covariance components **(C,G)** and non-direct variance components **(D,H)** are larger than those computed using the true estimate of the isotropic inflation factor.

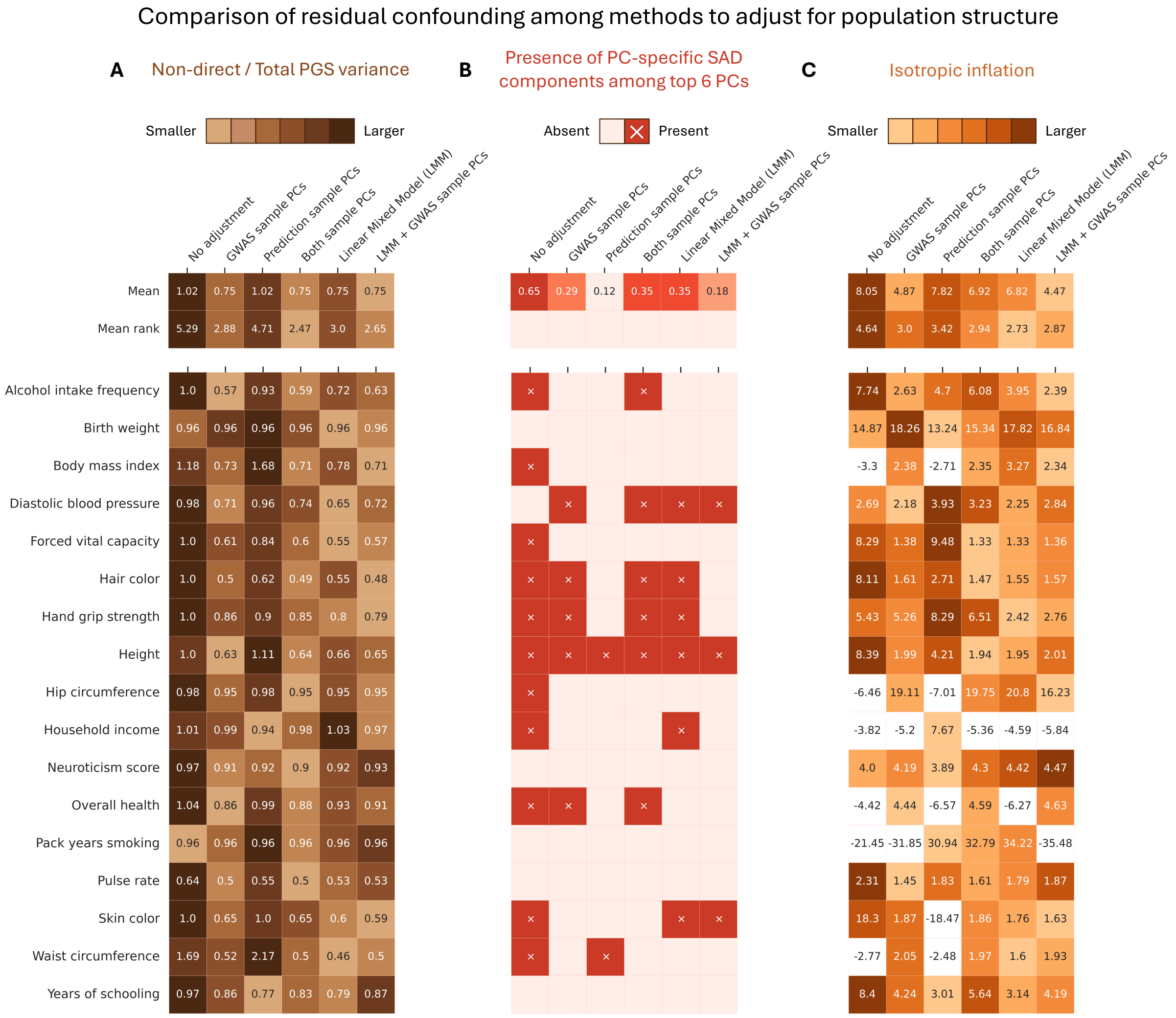

**Figure S4. The utility of GWAS population structure adjustments in mitigating confounding.** This figure is based on the same data as **Fig. 2C** of the main text but is more elaborate and contains details of the PGS-specific performance of each trait examined. We constructed PGSs using clumping and thresholding in a UKB-based GWASs with an ascertainment threshold of $p$-value $< 10^{-5}$. We then partitioned the PGS variance in the 1KG Europeans prediction sample. The PGSs were based on GWASs with various methods to adjust for population structure. "GWAS sample PCs" refers to a GWAS adjustment for the first 20 principal components of the genotype matrix of the UKB cohort in which the GWAS is performed. "Prediction sample PCs" refers to the inclusion of 20 1KG Europeans genotype matrix PCs. "LMM" refers to the *BOLT-LMM* [8] implementation of a linear mixed model. For each trait, **(A)** shows the non-direct variance component for each method. **(B)** shows the presence or absence of significantly large PC-specific SAD components ($p$-value $< 0.05$). **(C)** shows the isotropic inflation factor. For each trait, rank 1 is given to the GWAS adjustment method(s) that yielded the smallest non-direct variance component in **(A)**, the fewest significant PC-specific SAD variance components in **(B)**, and the smallest isotropic inflation factor in **(C)**. Rank 2 is given to the second fewest/smallest and so forth. The rank is reflected in the hue of each cell. The uppermost (second) row summarizes the mean value (rank) across traits. White cells represent trait-method pairs whose estimates for either the non-direct variance component or isotropic inflation are negative; these PGSs are not considered in the trait-wise ranking or in the calculation of method-wide values and ranks.

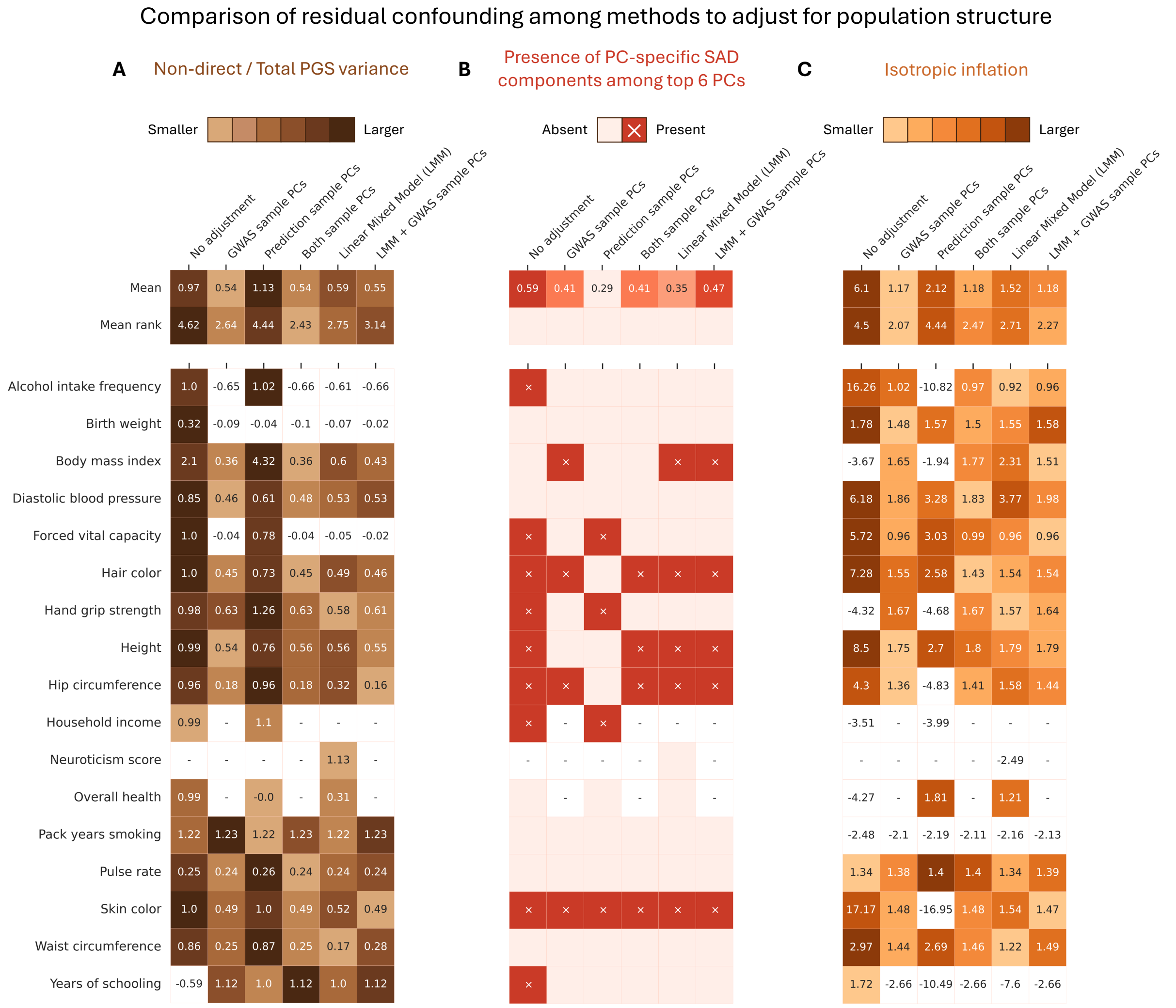

**Figure S5. The utility of GWAS population structure adjustments in mitigating confounding—with more stringent PGS index SNP ascertainment.** This figure is the same as **Fig. S4**, except that we constructed PGSs using a stricter ascertainment threshold ($p$-value $< 1 \times 10^{-8}$) on SNPs' marginal GWAS $p$-value. We then partitioned the PGS variance in the 1KG Europeans prediction sample. The PGSs were based on GWASs with various methods to adjust for population structure. "GWAS sample PCs" refers to a GWAS adjustment for the first 20 principal components of the genotype matrix of the UKB cohort in which the GWAS is performed. "Prediction sample PCs" refers to the inclusion of 20 1KG Europeans genotype matrix PCs. "LMM" refers to the *BOLT-LMM* [8] implementation of a linear mixed model. For each trait, **(A)** shows the non-direct variance component for each method. **(B)** shows the presence or absence of significantly large PC-specific SAD components ($p$-value$< 0.05$). **(C)** shows the isotropic inflation factor. For each trait, rank 1 is given to the GWAS adjustment method(s) that yielded the smallest non-direct variance component in **(A)**, the fewest significant PC-specific SAD variance components in **(B)**, and the smallest isotropic inflation factor in **(C)**. Rank 2 is given to the second fewest/smallest and so forth. The rank is reflected in the hue of each cell. The uppermost (second) row summarizes the mean value (rank) across traits. White cells represent trait-method pairs whose estimates for either the non-direct variance component or isotropic inflation are negative; these PGSs are not considered in the trait-wise ranking or in the calculation of method-wide values and ranks. Dashes in white cells indicate trait-method pairs which had an insufficient number of SNPs below the GWAS ascertainment threshold to construct a PGS with.

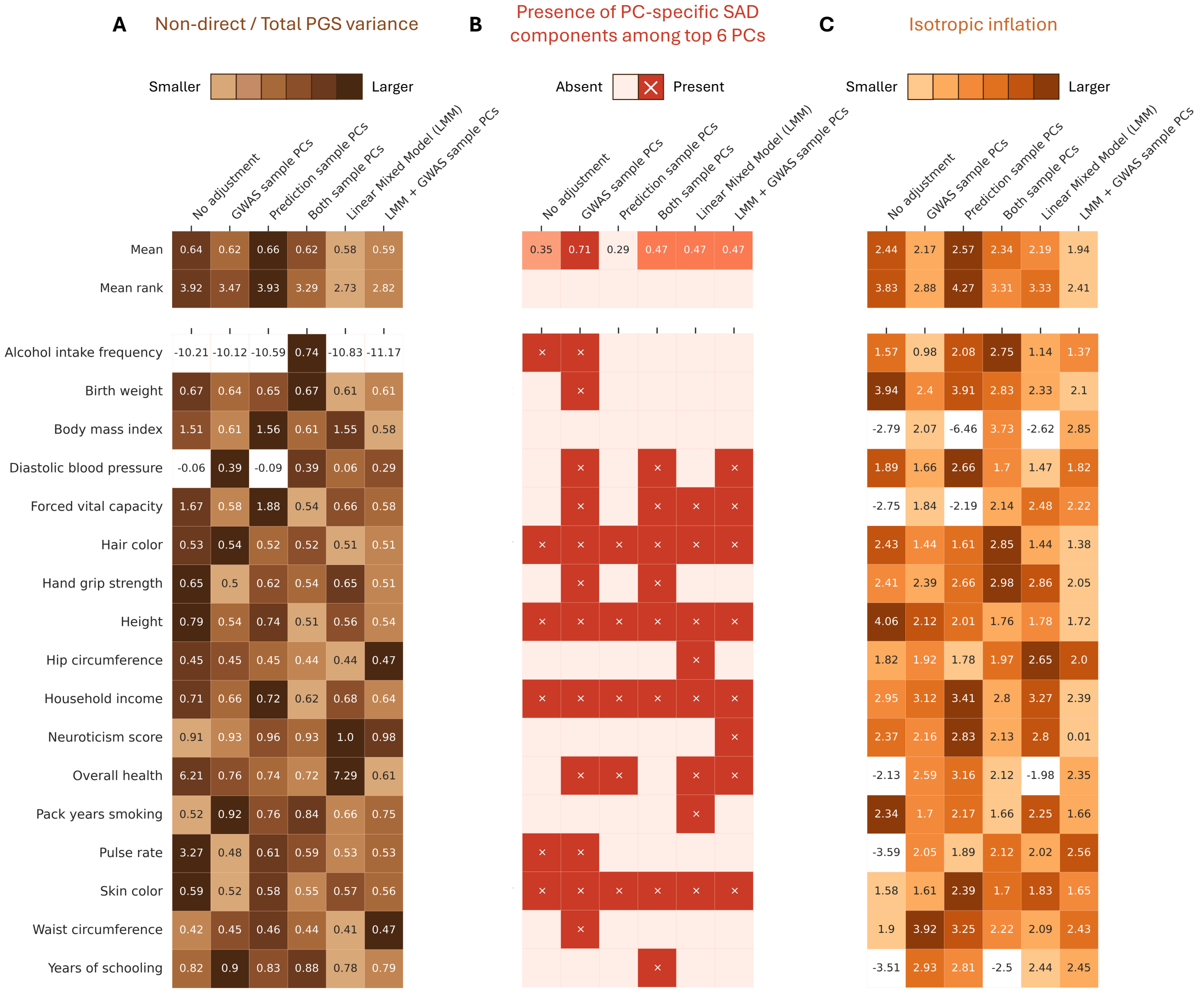

**Figure S6. The utility of population structure adjustments to UKB WB-based GWAS in mitigating confounding.** Using the UKB White British (UKB WB) cohort as our GWAS sample, we constructed PGSs using clumping in a GWAS with an ascertainment threshold of $p$-value $< 1\times10^{-5}$. We then partitioned the PGS variance in the 1KG Europeans prediction sample. The PGSs were based on GWASs with various methods to adjust for population structure. “GWAS sample PCs” refers to a GWAS adjustment for the first 20 principal components of the genotype matrix of the UKB White British cohort in which the GWAS is performed. “Prediction sample PCs” refers to the inclusion of 20 1KG Europeans genotype matrix PCs. “LMM” refers to the *BOLT-LMM*[8] implementation of a linear mixed model. For each trait, **(A)** shows the non-direct variance component for each method. **(B)** shows the presence or absence of significantly large PC-specific SAD components ($p$-value$< 0.05$). **(C)** shows the isotropic inflation factor. For each trait, rank 1 is given to the GWAS adjustment method(s) that yielded the smallest non-direct variance component in **(A)**, the fewest significant PC-specific SAD variance components in **(B)**, and the smallest isotropic inflation factor in **(C)**. Rank 2 is given to the second fewest/smallest and so forth. The rank is reflected in the hue of each cell. The uppermost (second) row summarizes the mean value (rank) across traits. White cells represent trait-method pairs whose estimates for either the non-direct variance component or isotropic inflation are negative; these PGSs are not considered in the trait-wise ranking or in the calculation of method-wide values and ranks.

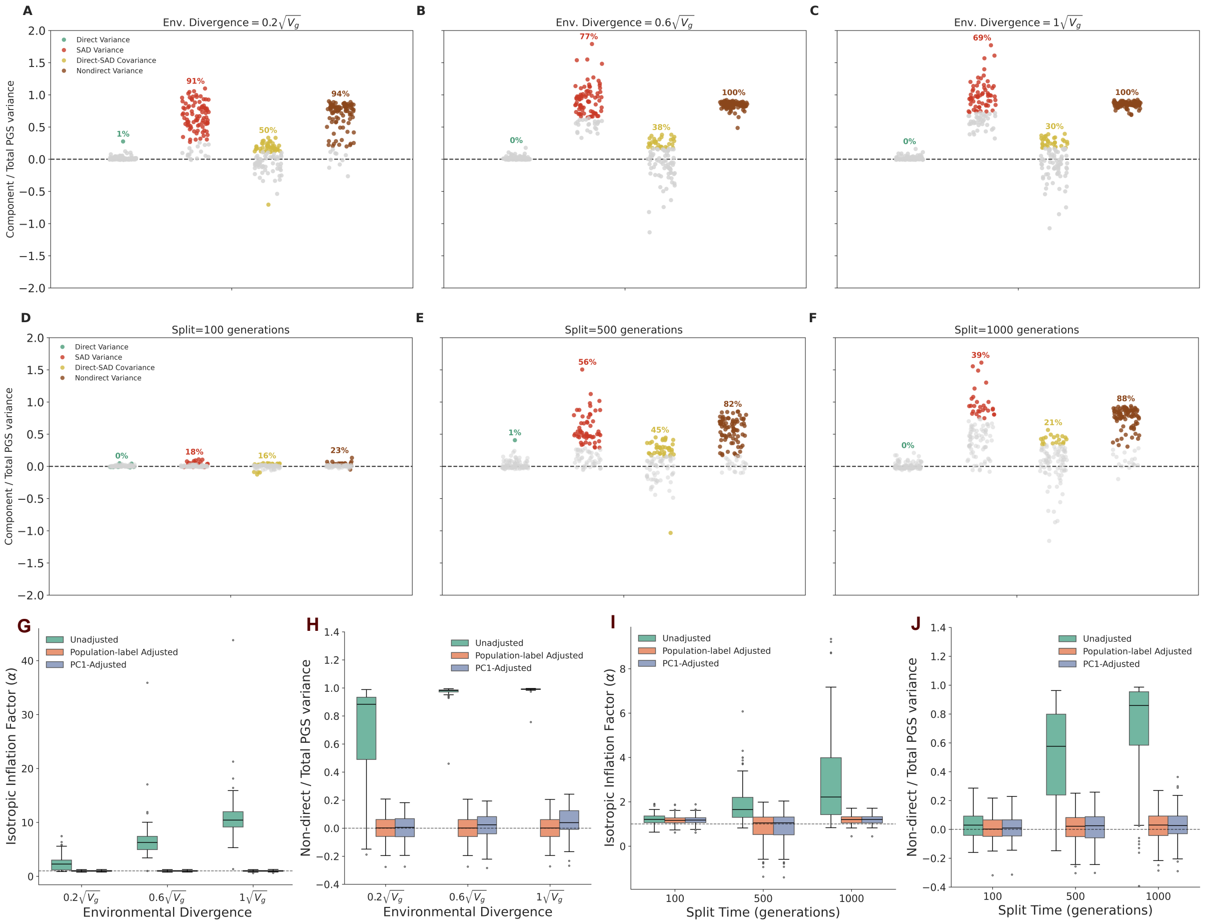

**Figure S7. Population stratification in a two-population split model.** We performed variance decomposition on PGS constructed with GWAS performed on simulated populations that are stratified due to either genetic drift alone or drift + environmental divergence (n=2,500 for each of the standard-GWAS, sib-GWAS, and prediction sample cohorts, see **Text S14.4**). **(A-C)** Variance components along the first PC of the prediction sample in simulations with varying levels of environmental divergence (which scales with genetic variance, $V_g$) between two subpopulations split 500 generations ago with effective population sizes of 10,000. **(D-F)** Variance components along the first PC under varying levels of split time without environmental divergence. In panels **(A-F)**, the percentage above each cluster of points indicates the proportion of simulation iterations in which the estimated component falls outside the null non-rejection region; gray points indicate iterations in which the estimated components fall inside the null non-rejection region. **(G)** Distribution of the isotropic inflation factor under varying environmental divergence and with three different strategies for stratification correction (green = no correction, orange = adjustment for population labels, purple = adjustment for individual position on PC1). **(H)** Distribution of the non-direct variance component under varying environmental divergence. **(I)** Distribution of the isotropic inflation factor under varying split times. **(J)** Distribution of the non-direct variance component under varying split times.

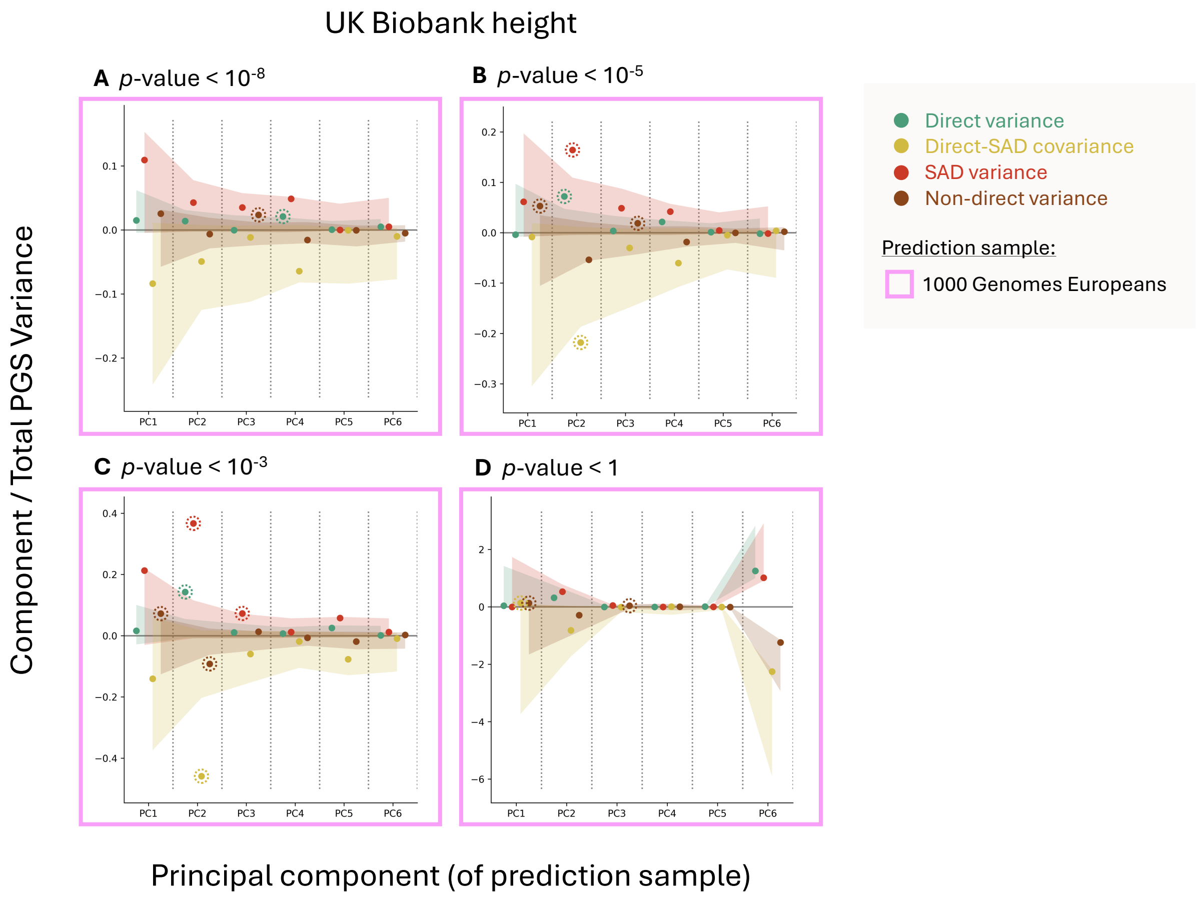

**Figure S8. Variance partitionings for PGSs for height using UKB summary statistics.** PGSUS partitions the variance of a PGS among individuals in a prediction sample. Variance components are attributable to an effect type—direct effects, SAD effects and their covariance—and a principal component of the genotype matrix of the prediction sample. Shown are variance components (divided by the total variance in the PGS) along the first six PCs for four PGSs constructed for the UK Biobank standard-GWAS summary statistics for height using different ascertainment thresholds. Shaded regions show empirical permutation-based null non-rejection regions. Components deviating from this expectation are highlighted with dashed circles.

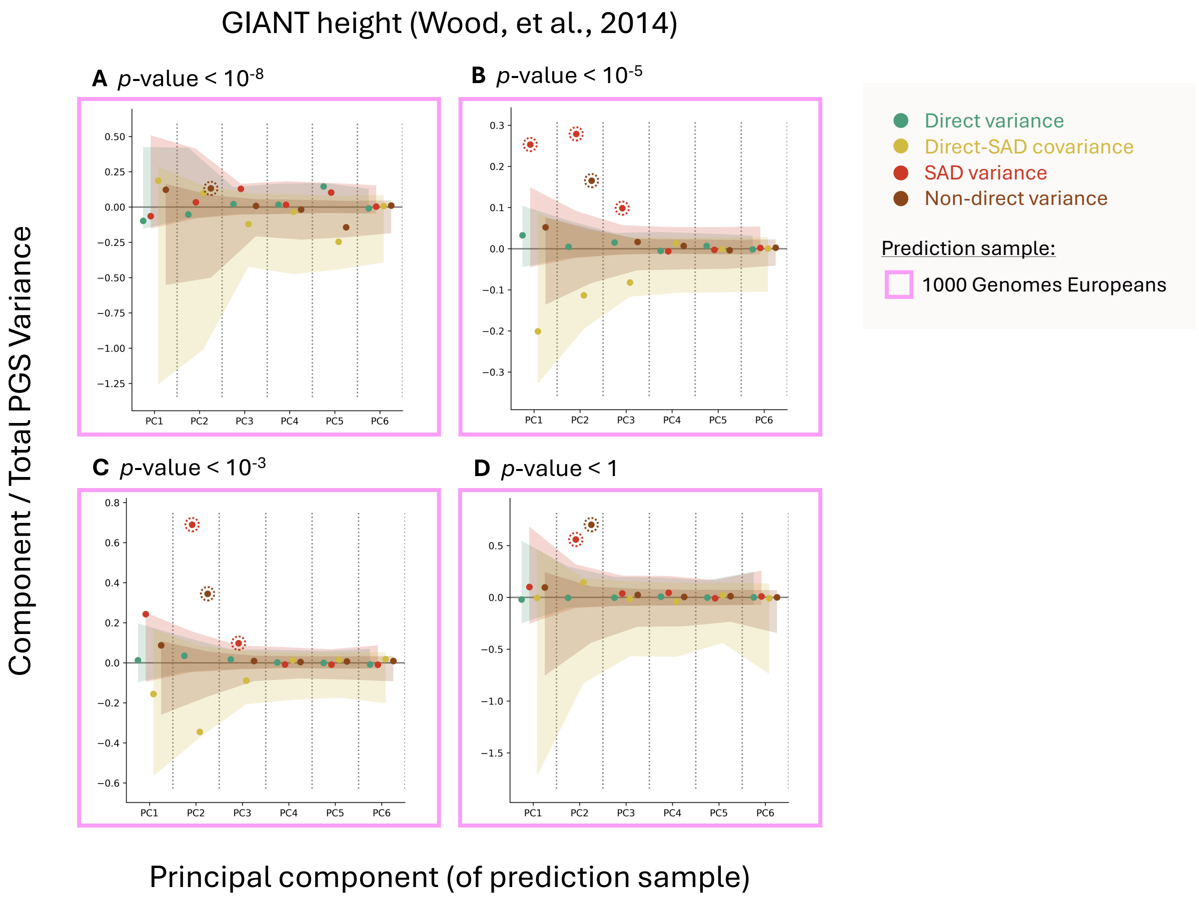

**Figure S9. Variance partitionings for PGSs for height using the GIANT consortium 2014 summary statistics.** PGSUS partitions the variance of a PGS among individuals in a prediction sample. Variance components are attributable to an effect type—direct effects, SAD effects and their covariance—and a principal component of the genotype matrix of the prediction sample. Shown are variance components (divided by the total variance in the PGS) along the first six PCs for four PGSs constructed for the GIANT 2014 standard-GWAS summary statistics for height using different ascertainment thresholds. Shaded regions show empirical permutation-based null non-rejection regions. Components deviating from this expectation are highlighted with dashed circles.

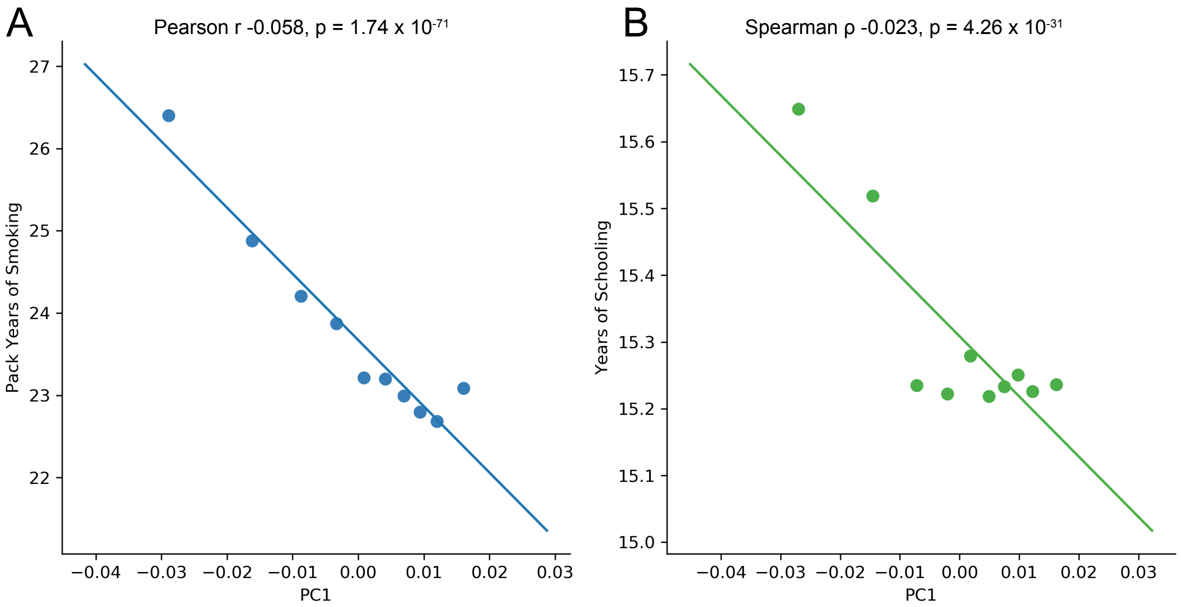

**Figure S10. Correlations between individual PC coordinates and two measures related to forced vital capacity.** X-axis values show means of PC1 deciles. **(A)** Y-axis values show mean pack years of smoking per bin. **(B)** Y-axis values show mean years of schooling (calculated using the ordinal scale of Okbay et al. [61]) per bin. Linear fits (solid lines) and correlations are based on the raw, un-binned data.

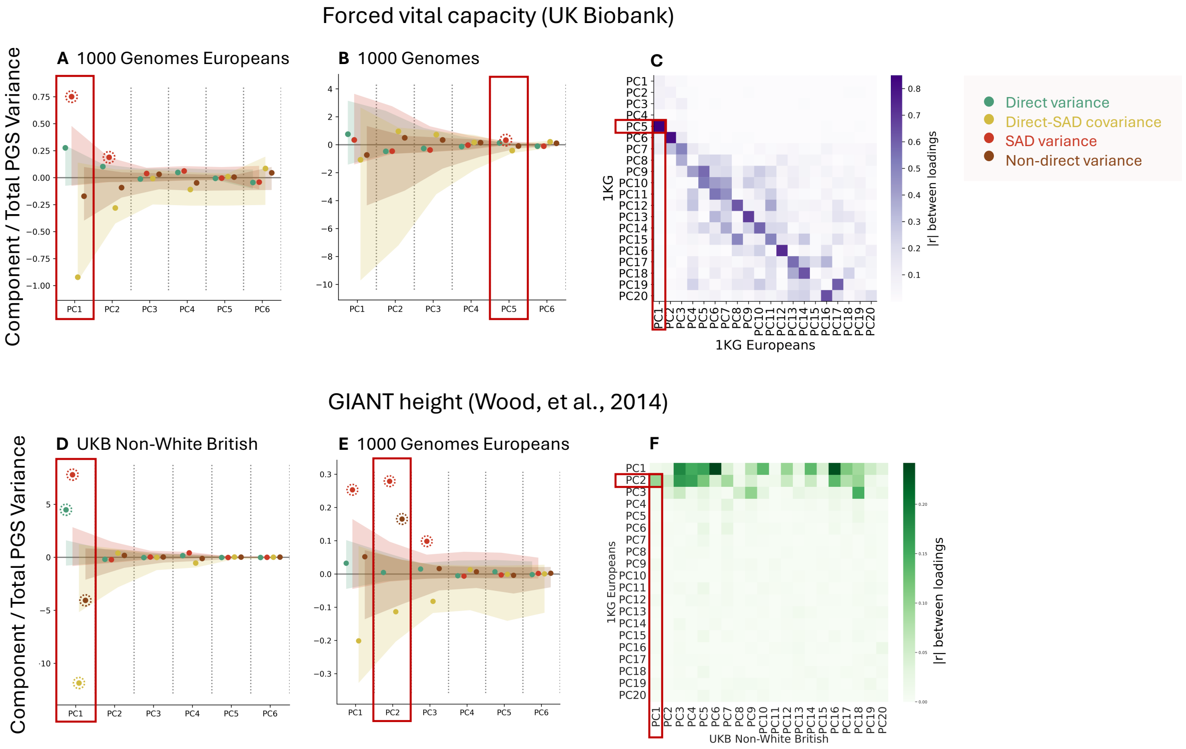

**Figure S11. Signals of stratification observed in distinct prediction samples along correlated axes of ancestry. (A)** Partitioning of variance in a UK Biobank (UKB)-based PGS of forced vital capacity applied to 1KG Europeans. A significant SAD variance component is present along PC1 and PC2. **(B)** Partitioning of variance in a PGS of forced vital capacity applied to 1KG individuals. A significant SAD variance component is present along PC5. **(C)** Correlation of the SNP loadings for the first 20 PCs of the 1KG Europeans and 1KG cohorts, highlighting the high correlation between 1KG Europeans PC1 and 1KG PC5 (Pearson $|r| = 0.933$, $p < 1 \times 10^{-250}$). This correlation suggests that these two PCs in these distinct samples tag a correlated axis of ancestry. (This panel is the same as **Fig. S31C**.) **(D)** Partitioning of variance in a GIANT height PGS applied to the UKB Non-White British (UKB NWB) prediction sample. A significant SAD variance component is present along PC1. **(E)** Partitioning of variance in a GIANT height PGS applied to 1KG Europeans, showing a significant SAD variance component along PC2. **(F)** Correlation of the SNP loadings for the first 20 PCs of the UKB NWB and 1KG European cohorts, highlighting the significant correlation between UKB NWB PC1 and 1KG Europeans PC2 (Pearson $|r| = 0.096$, $p < 1 \times 10^{-258}$). This correlation suggests that these two PCs in these distinct samples tag a correlated axis of ancestry. All PGSs shown were ascertained at GWAS $p$-value $< 10^{-5}$.

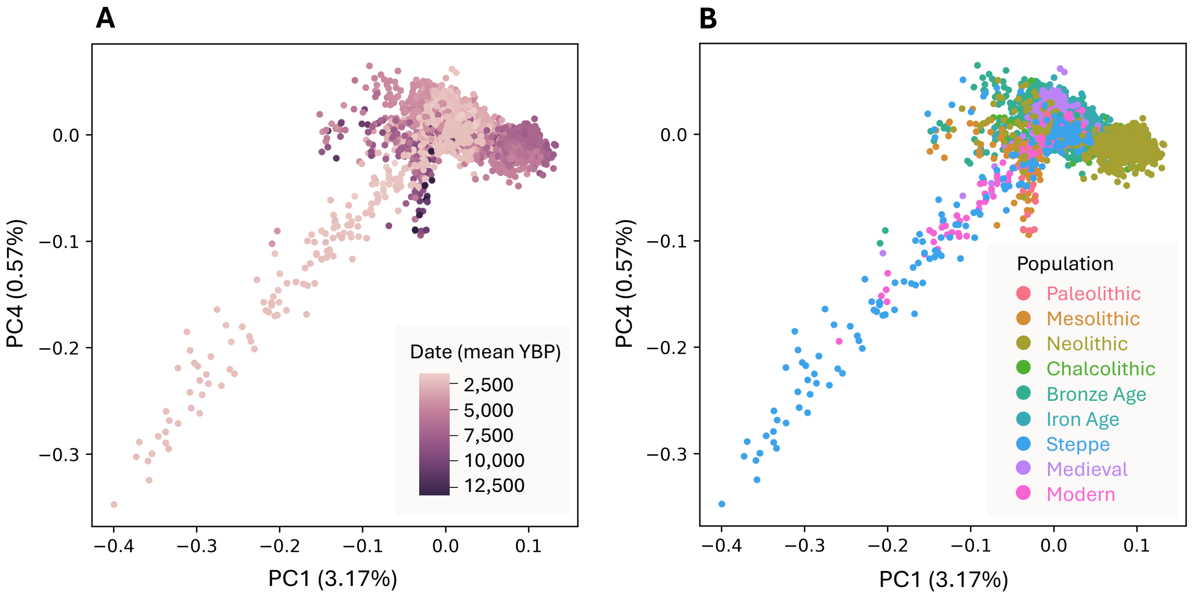

**Figure S12. Principal components analysis of 4,588 ancient Eurasians. (A)** Individuals shown in PC1-by-PC4 space colored by sample date in mean years before present (YBP). **(B)** Individuals shown in PC1-by-PC4 space colored by population identifiers. The population identifiers are Paleolithic (n=16), Mesolithic (n=125), Neolithic (n=791), Chalcolithic (n=240), Bronze Age (n=1,457), Iron Age (n=746), Steppe Pastoralist (n=301), Medieval (n=654), or Modern (n=221). 37 individuals did not have a population assigned. Individuals were assigned to a population using a combination of $f_4$-statistics, sample date (based on either radiocarbon dating or archaeological context), and geographic location[15,16]. PCs were computed from ancient genotypes themselves (as opposed to conventional projection into a PC space computed from modern populations).

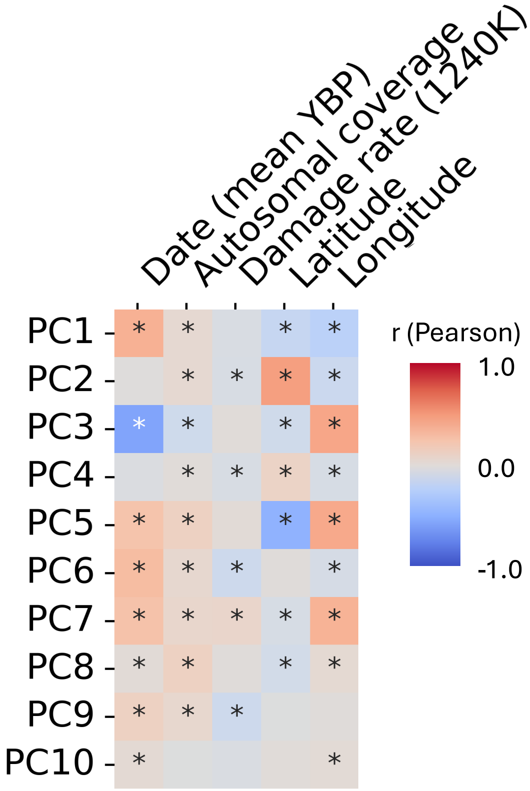

**Figure S13. Correlation between top 10 PCs and sequencing and sample metadata for AADR Eurasian subsample.** Pearson correlations were computed between sequence and sample metadata and the first 10 PCs ($*$ denotes $p$-value < 0.05). 1240K refers to the 1240K target array which allows for in-solution enrichment of low-volume or contaminated samples[62–65]. Date is the mean sample date of the date range determined by radiocarbon dating or well-understood archaeological context in years before present (YBP). Coverage refers to the fraction of reference genome that ancient sequences successfully mapped to. Damage rate (1240K) refers to the fraction of the 1240K array sequence positions with errors due to DNA damage[66]. We find significant correlations between individual-level PC coordinates and quality metrics, as well as with the location and date of the sample. Sample and sequencing metadata was sourced from Allen Ancient DNA Resource (AADR, v62.0)[15].

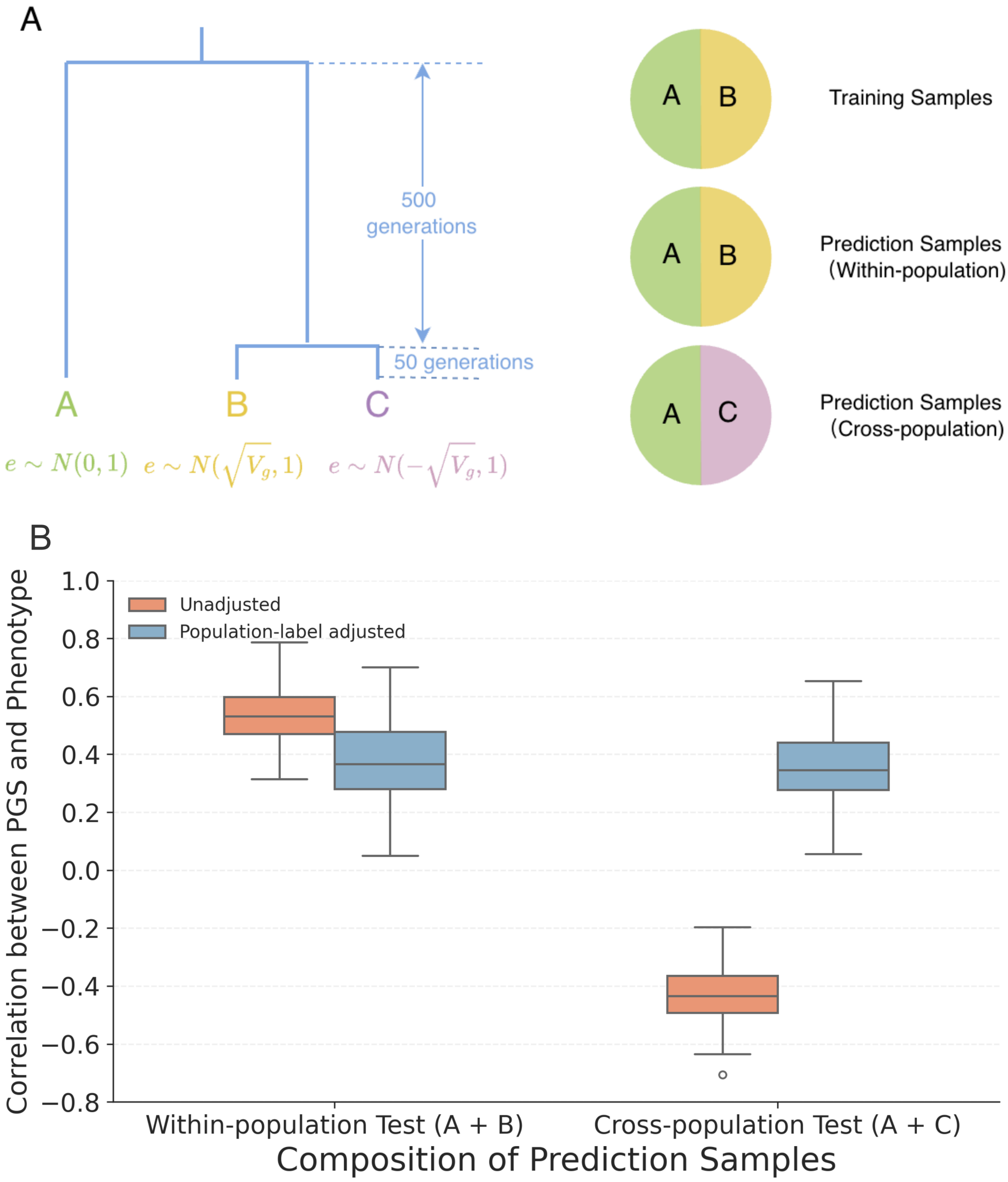

**Figure S14. PGS portability across simulated populations.** We tested the correlation between phenotype and PGS in a scenario where the GWAS and prediction sample either share or have different subpopulation compositions. **(A)** Simulation framework for our three-population split model used to generate a GWAS training sample and two prediction samples (**Text S14.10**). **(B)** Distributions of Pearson correlation coefficients between PGS and phenotype across test samples, comparing allelic effects derived from unadjusted GWAS (blue) and GWAS adjusted for population label as a covariate (orange).

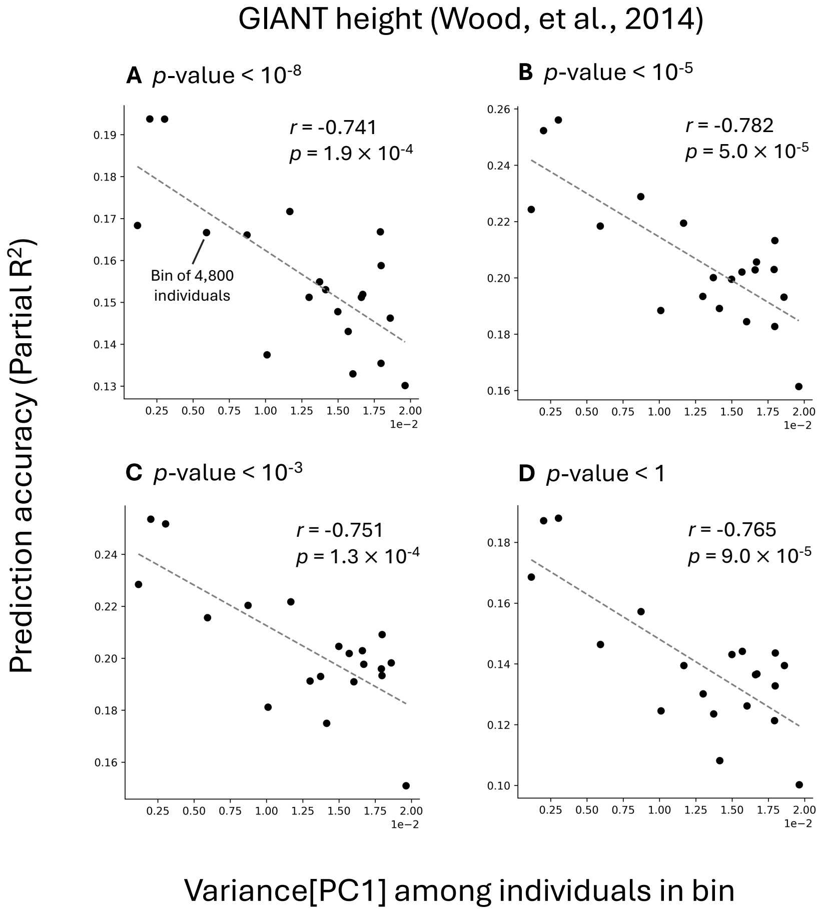

**Figure S15. PGS prediction accuracy for GIANT height as a function of cohort variance along a stratified axis at different GWAS ascertainment thresholds.** This figure relates to the same analysis as in **Fig. 5B** in the main text, but with variable index SNP ascertainment stringency for the PGS. Using the Non-White British cohort from the UK Biobank, we generated 20 bins of 4,800 individuals with increasing variance along PC1 (which showed evidence of stratification in **Fig. 5A**). In each bin, we regressed height to a linear combination of the PGS, age, age$^2$, sex, age × sex, and age$^2$ × sex and measured PGS prediction accuracy for the bin (measured as partial $R^2$ between the PGS and the trait). $p$-values in panel titles refer to GWAS ascertainment thresholds for SNP inclusion in the PGS.

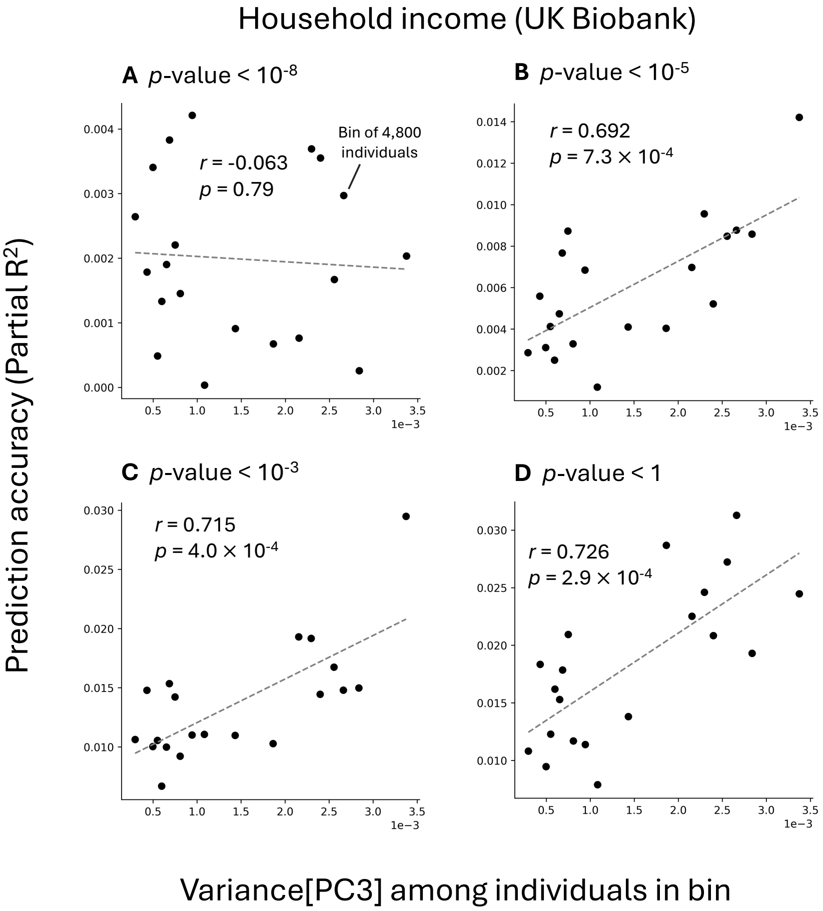

**Figure S16. PGS prediction accuracy for household income as a function of cohort variance along a stratified axis at different GWAS ascertainment thresholds.** This figure relates to the same analysis as in **Figs. 5D** in the main text, but with variable index SNP ascertainment stringency for the PGS. Using the Non-White British cohort from the UK Biobank, we generated 20 bins of 4,800 individuals with increasing variance along PC3 (which showed evidence of stratification in **Fig. 5C**). In each bin, we regressed household income to a linear combination of the PGS, age, $age^2$, sex, age × sex, and $age^2$ × sex and measured PGS prediction accuracy for the bin (measured as partial $R^2$ between the PGS and the trait). $p$-values in panel titles refer to GWAS ascertainment thresholds for SNP inclusion in the PGS.

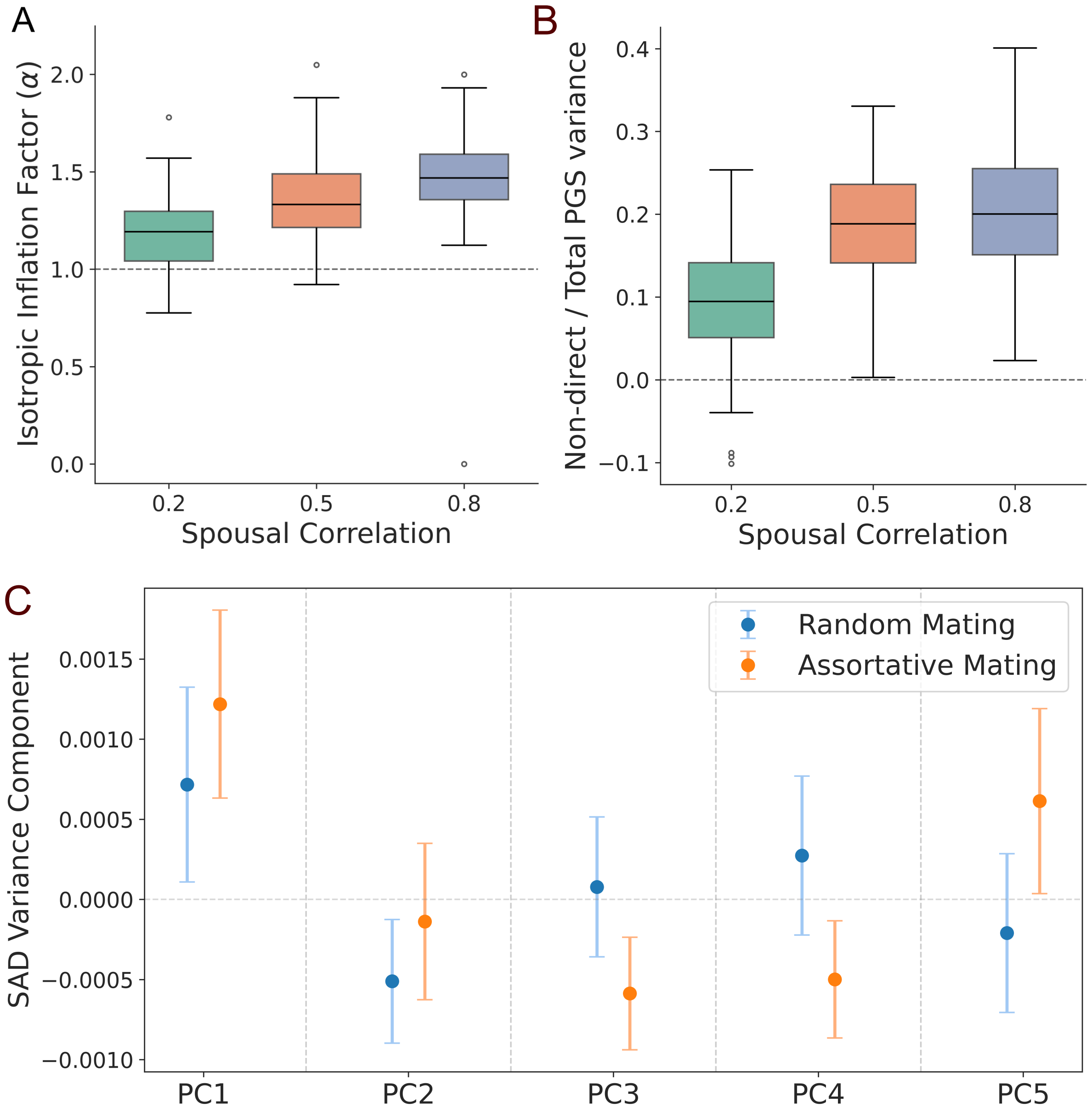

**Figure S17. Simulation of assortative mating in a single population.** We performed variance decomposition on PGSs constructed with GWASs performed on a simulated, assortatively mating population (as indicated by spousal correlation, see **Text S14.5**). **(A)** Distribution of isotropic inflation factor estimates under varying spousal correlation. **(B)** Distribution of the non-direct variance component under varying spousal correlation. **(C)** Average (points) and standard deviation (vertical error bars) of the SAD variance components along the first six PCs under random mating (blue) and assortative mating (orange, spousal correlation $\rho = 0.2$) across 100 iterations.

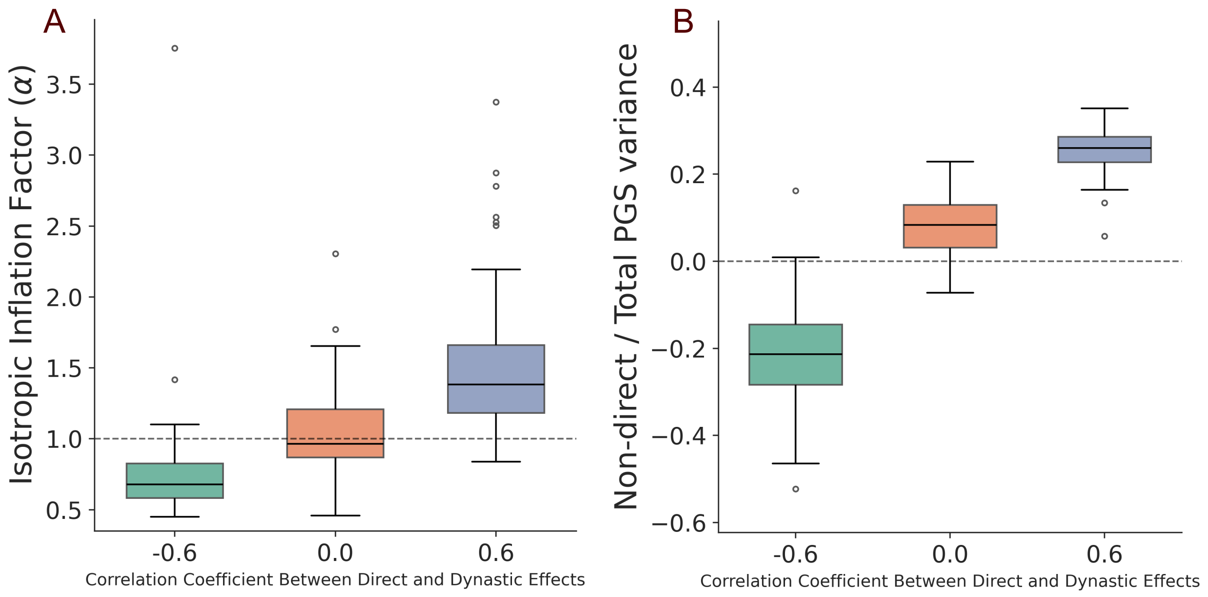

**Figure S18. Distribution of isotropic inflation factor estimates and non-direct variance components across simulations with dynastic effects.** We performed variance decomposition on PGSs for a simulated trait affected by dynastic effects (**Text S14.6**). **(A)** Distribution of isotropic inflation factor estimates across simulations in which dynastic effects are negatively correlated ($\rho = -0.6$), uncorrelated ($\rho = 0$), or positively correlated ($\rho = 0.6$) with direct genetic effects. Positive correlation between dynastic and direct effects leads to an average isotropic inflation factor above one, while negative correlation leads to it falling below one. Gray dashed line represents no isotropic inflation ($\alpha = 1$). **(B)** Distribution of non-direct variance components under varying correlation coefficients between direct and dynastic effects.

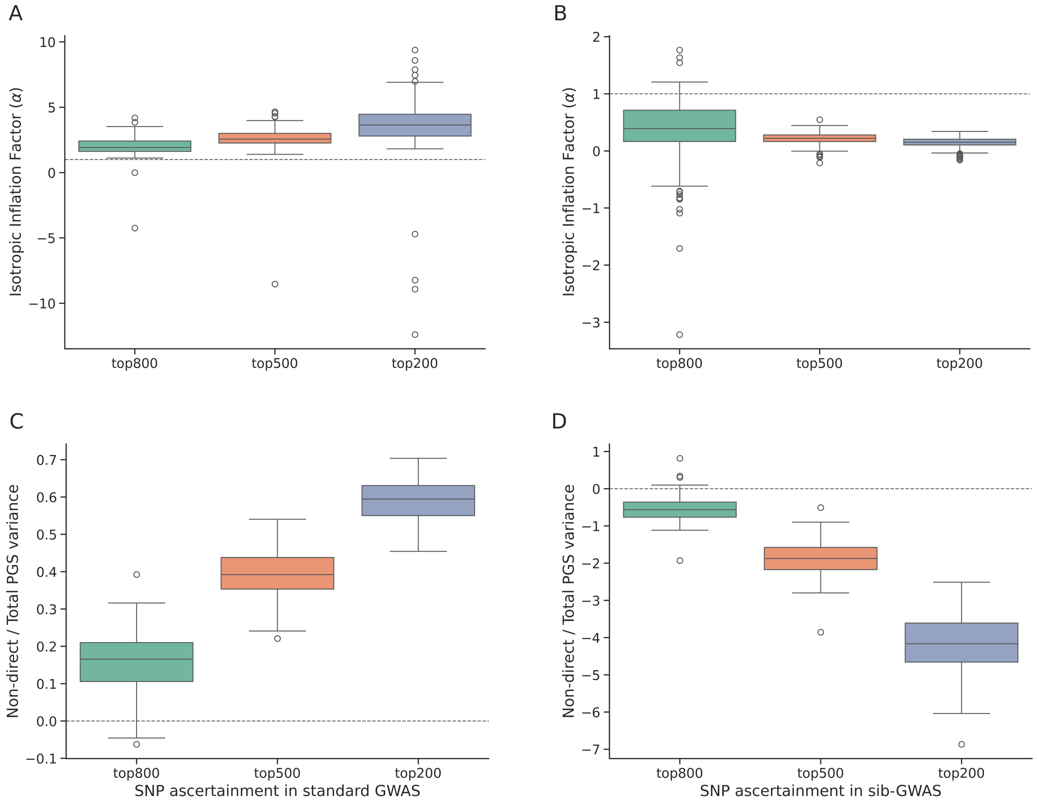

**Figure S19. The effect of GWAS ascertainment on estimated isotropic inflation factors and non-direct variance components in simulated genetic data. (A)** The distribution of estimated isotropic inflation factors when the significant SNPs are ascertained on the basis of standard-GWAS-estimated $p$-values. **(B)** The distribution of estimated isotropic inflation factors when the significant SNPs are ascertained on the basis of sib-GWAS-estimated $p$-values. The horizontal dashed line in **(A,B)** indicates an isotropic inflation factor equal to one. **(C)** The distribution of estimated non-direct variance component when the significant SNPs are ascertained on the basis of standard-GWAS-estimated $p$-values. **(D)** The distribution of estimated non-direct variance component when the significant SNPs are ascertained on the basis of sib-GWAS-estimated $p$-values. The horizontal dashed lines in **(C,D)** indicate a non-direct variance component of zero. (Because the non-direct variance component contains a covariance term, it can be negative.)

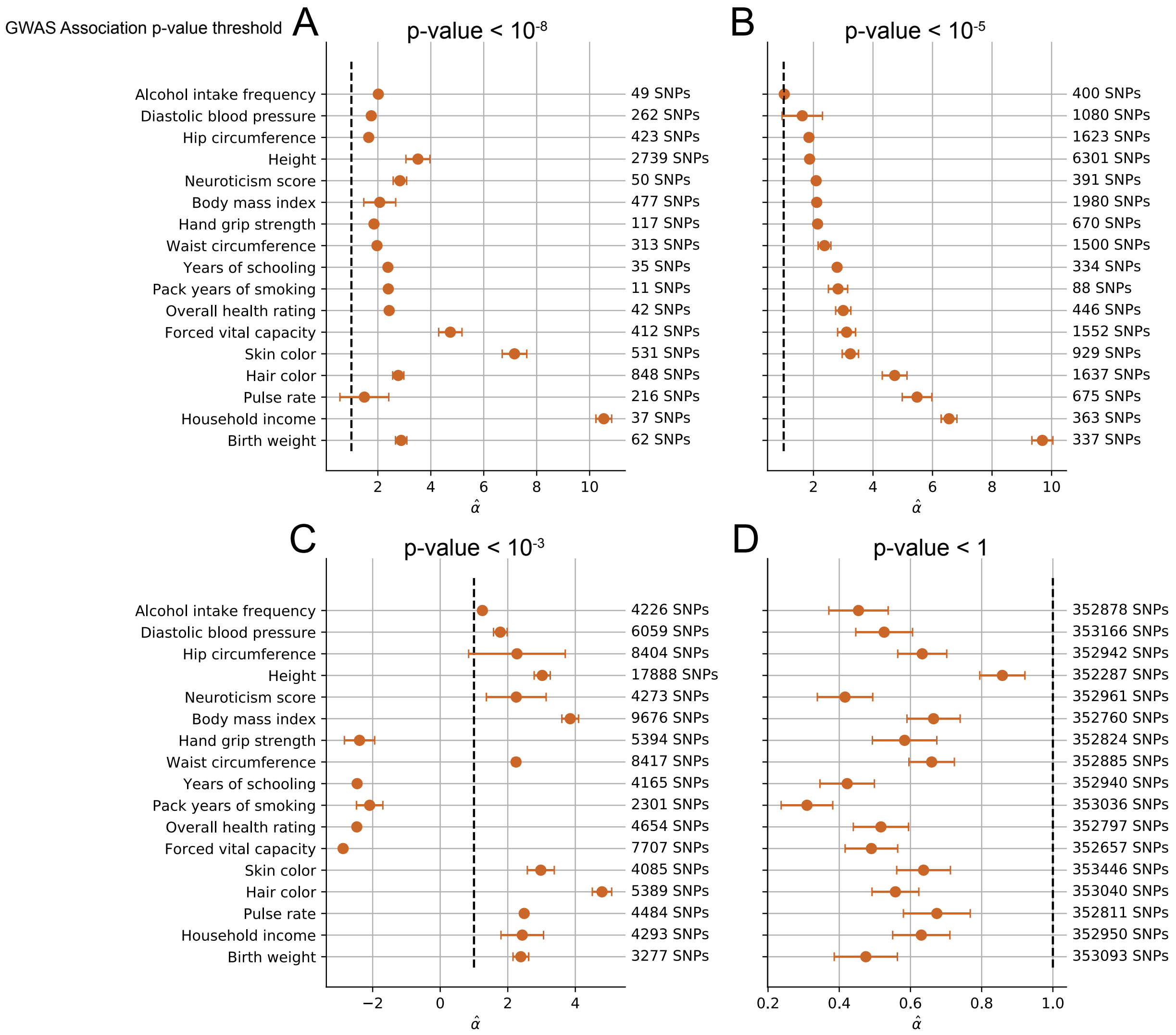

**Figure S20. The isotropic inflation factor at four ascertainment thresholds in the 1KG prediction sample.** Panels **(A)** through **(D)** show isotropic inflation estimates for each of 17 traits using ascertainment thresholds $p$-value $< 10^{-8}$, $p$-value $< 10^{-5}$, $p$-value $< 10^{-3}$, and $p$-value $< 1$, respectively. Isotropic inflation refers to SAD variance that is not PC-specific, and manifests as systematic differences in the magnitude of allelic effects estimated by standard GWAS and sib-GWAS. A factor of 1 indicates no isotropic inflation (dashed vertical line). Counts for the index SNPs for each PGS are provided on the right.

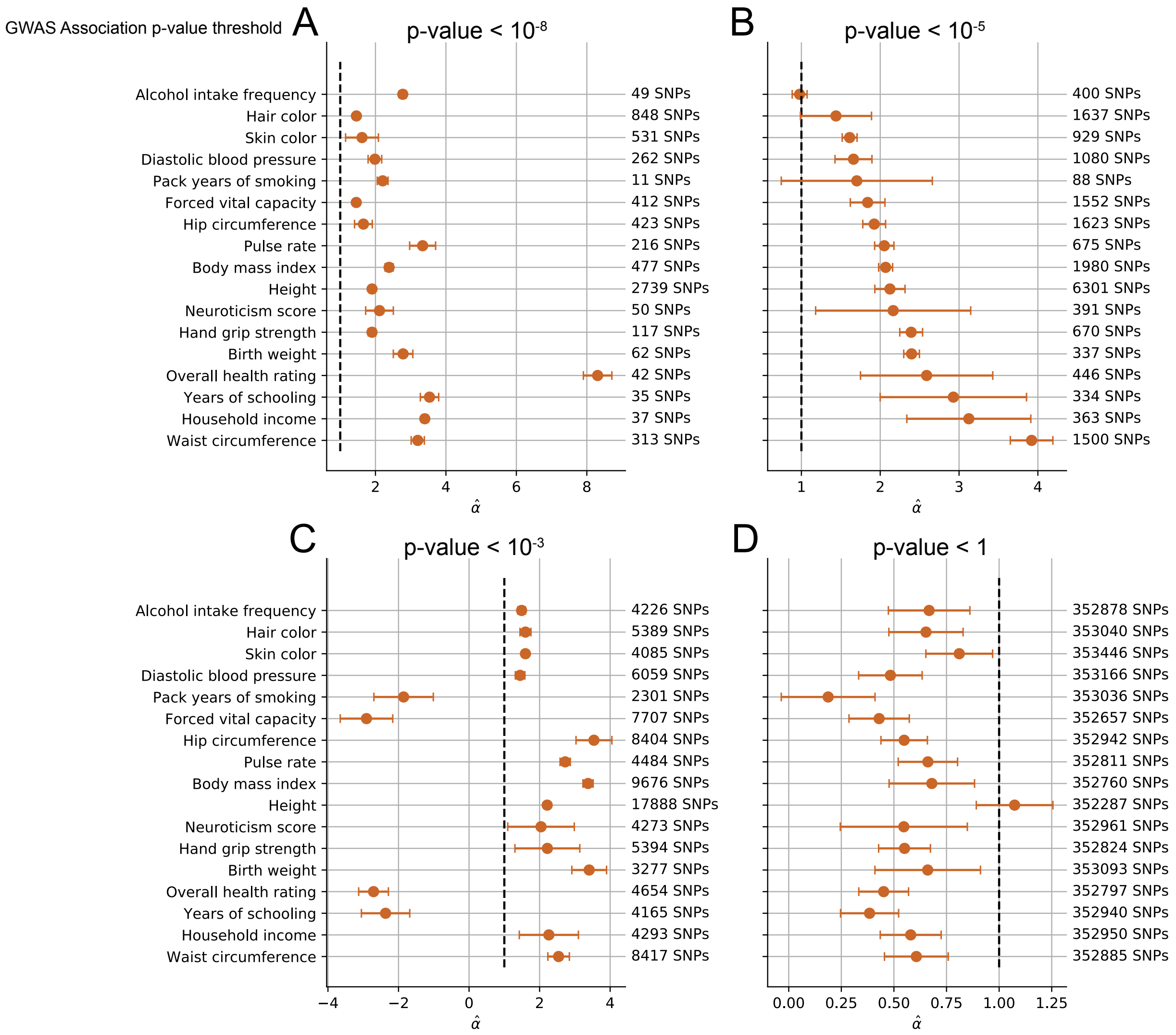

**Figure S21. The isotropic inflation factor at four ascertainment thresholds in the 1KG Europeans prediction sample.** Panels **(A)** through **(D)** show isotropic inflation estimates for each of 17 traits using ascertainment thresholds $p$-value < $10^{-8}$, $p$-value < $10^{-5}$, $p$-value < $10^{-3}$, and $p$-value < 1, respectively. Isotropic inflation refers to SAD variance that is not PC-specific and that manifests as systematic differences in the magnitude of allelic effects estimated by standard GWAS and sib-GWAS. A factor of 1 indicates no isotropic inflation (dashed vertical line). Counts for the index SNPs for each PGS are provided on the right.

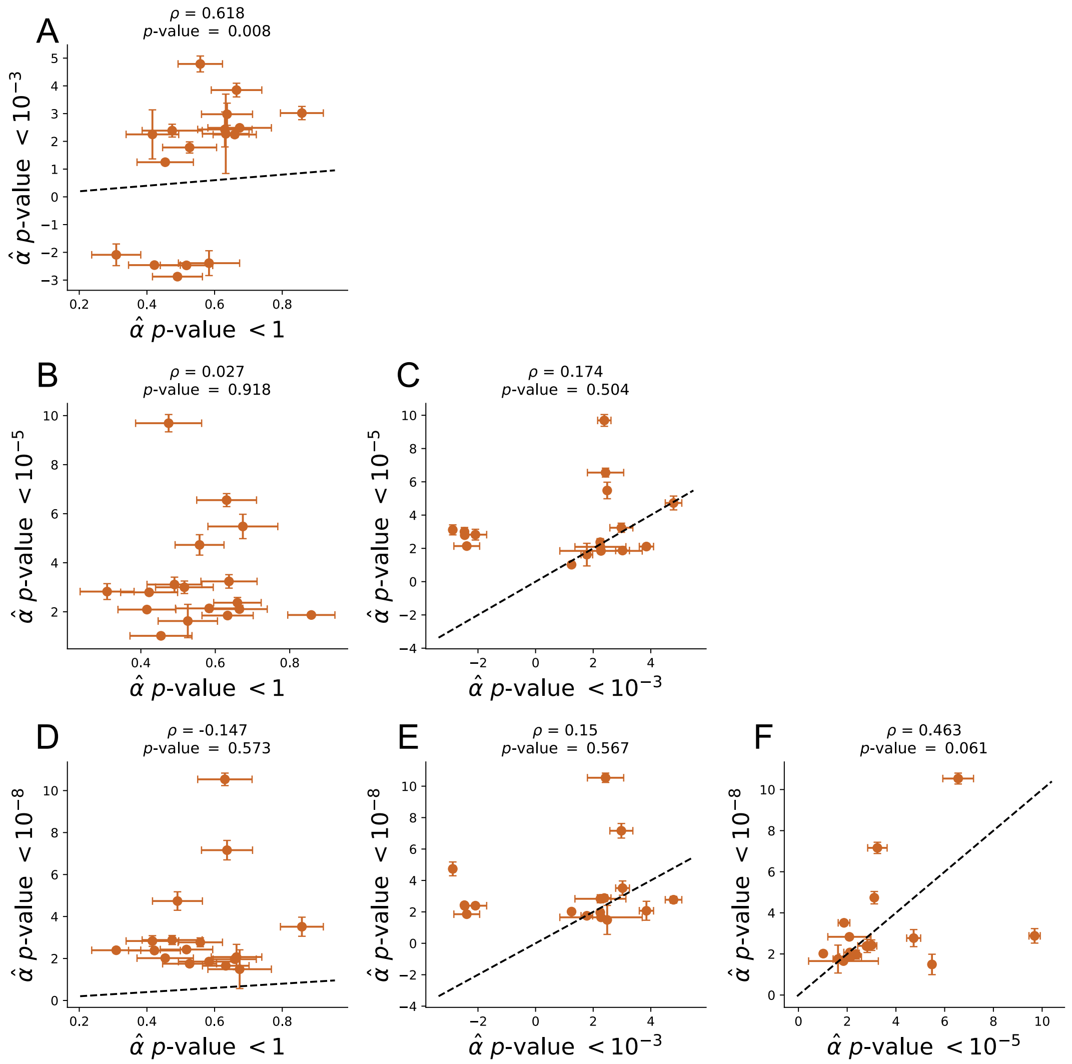

**Figure S22. Correlation between isotropic inflation factor estimates for PGSs constructed using different ascertainment thresholds.** Panels **(A)** through **(F)** show the correlation between isotropic inflation estimates for PGSs for 17 traits constructed using different ascertainment thresholds ($p$-value $< 10^{-8}$, $p$-value $< 10^{-5}$, $p$-value $< 10^{-3}$, or $p$-value $< 1$) applied to the 1KG prediction sample. Isotropic inflation refers to SAD variance that is not PC-specific and that manifests as systematic differences in the magnitude of allelic effects estimated by standard GWAS and sib-GWAS. The dashed line represents a 1:1 relationship.

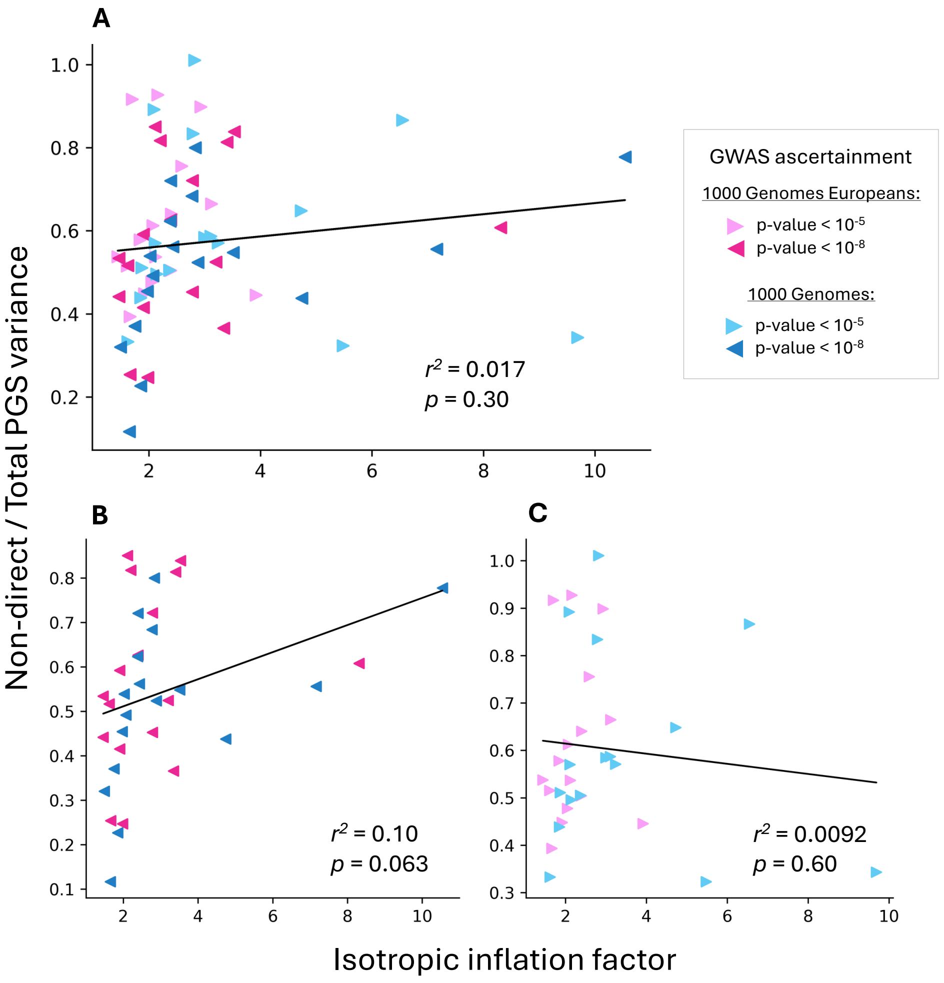

**Figure S23. The correlation between isotropic inflation and the non-direct variance component.** We evaluated the correlation between the isotropic inflation factor and the non-direct variance component for 34 PGSs based on GWAS in the UKB White British cohort for the 17 traits in **Table S1**, applied to both the 1KG and 1KG Europeans cohorts at two GWAS ascertainment thresholds. **(A)** shows all PGSs together, and **(B)**,**(C)** show the fits for each of the two GWAS ascertainment thresholds separately.

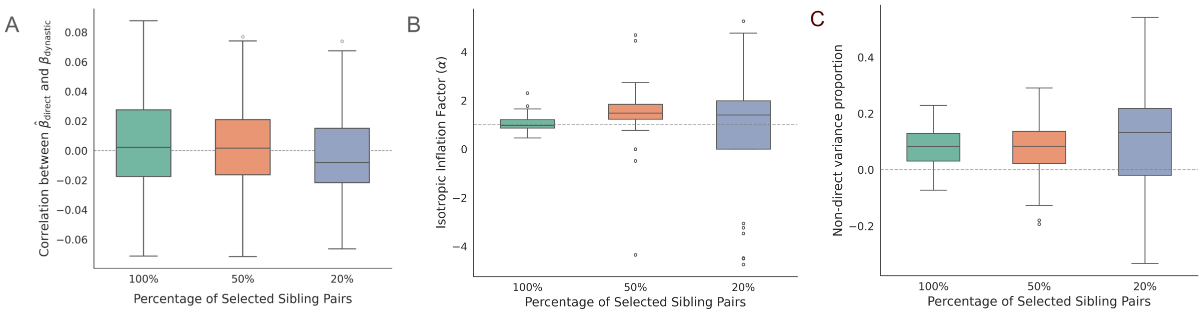

**Figure S24. Simulation of trait-biased sampling based on both direct and dynastic effects.** We simulated trait-biased sampling by ascertaining sibling pairs for the sib-GWAS under three schemes: random sampling (100%), inclusion of sibling pairs with mean phenotypes in the top 50% phenotype, and inclusion of sibling pairs with mean phenotypes in the top 20%. **(A)** Distribution of Pearson correlations between sib-GWAS-estimated direct effects and simulated dynastic effects under the three ascertainment levels. **(B)** Distribution of isotropic inflation factor estimates under the same three ascertainment levels. **(C)** Distribution of the non-direct variance component under the same three ascertainment levels.

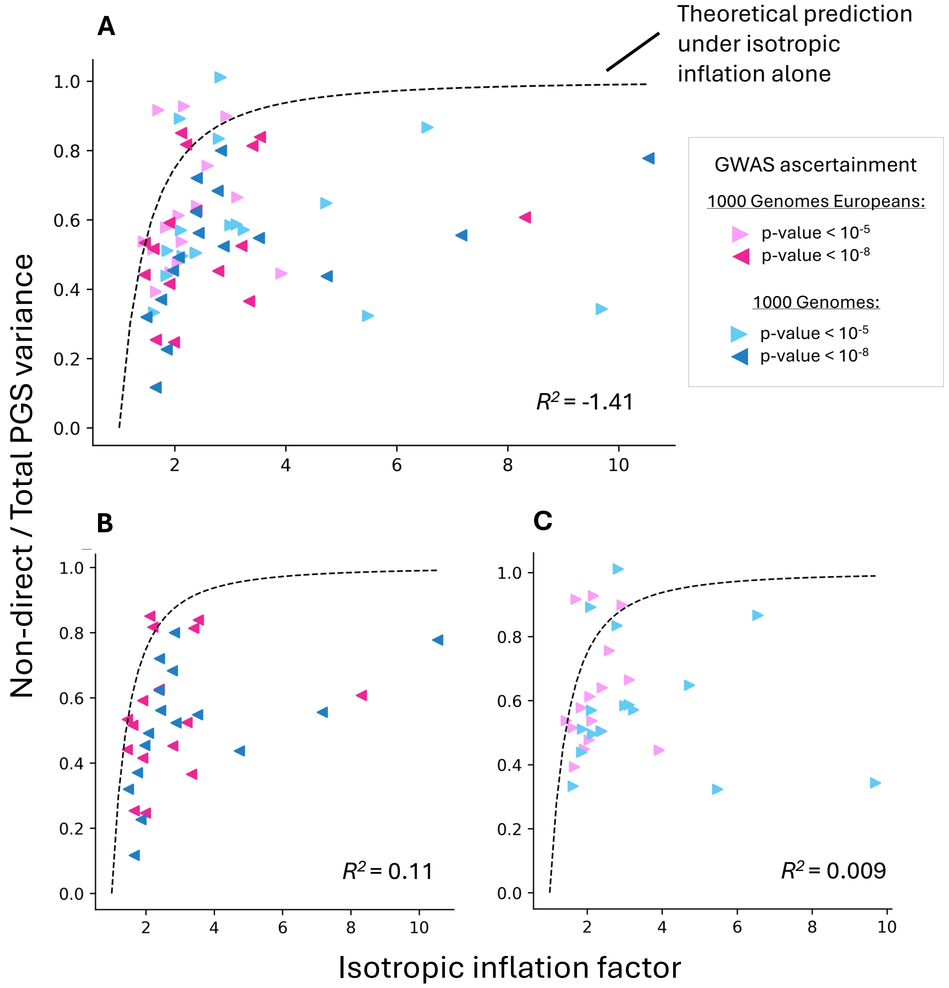

**Figure S25. Isotropic inflation alone does not fully explain non-direct variance.** We considered a theoretical model (represented by the dashed line) wherein no PC-specific SAD effects are at play, and all non-direct variance is due to isotropic inflation. We evaluated the fit to this model for 34 PGSs based on GWAS in the UKB WB for the 17 traits in **Table S1**, applied to both the 1KG and 1KG Europeans cohorts at two GWAS ascertainment thresholds. **(A)** shows all PGSs together, and **(B)**,**(C)** show the fits for each of the two GWAS ascertainment thresholds separately.

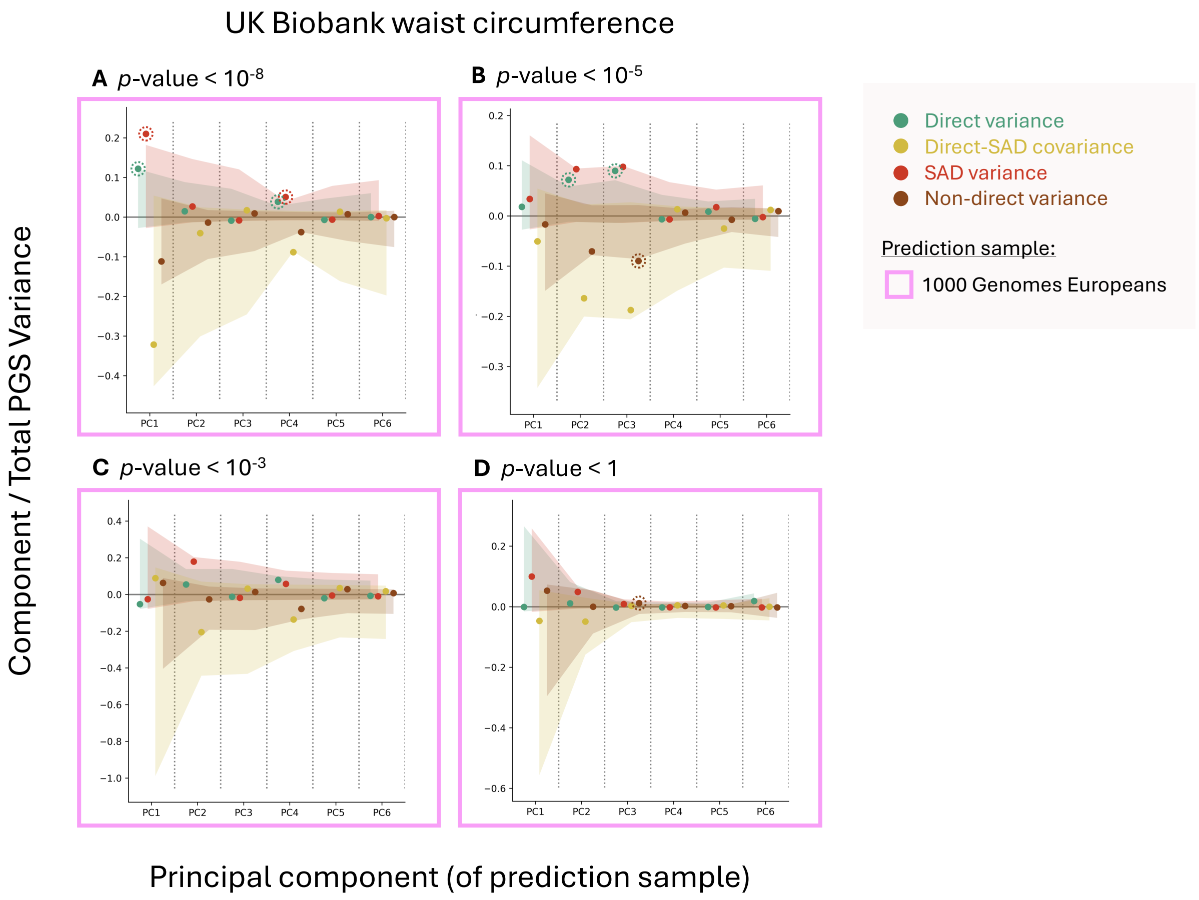

**Figure S26. Variance partitionings for waist circumference PGSs in the 1KG Europeans prediction sample.** PGSUS partitions the variance of a PGS among individuals in a prediction sample. Variance components are attributable to an effect type—direct effects, SAD effects and their covariance—and a principal component of the genotype matrix of the prediction sample. Shown are variance components (divided by the total variance in the PGS) along the first six PCs for four PGSs constructed for the UK Biobank standard-GWAS summary statistics for waist circumference using different ascertainment thresholds applied to the 1KG Europeans prediction sample. Shaded regions show empirical permutation-based null non-rejection regions. Components deviating from this expectation are highlighted with dashed circles.

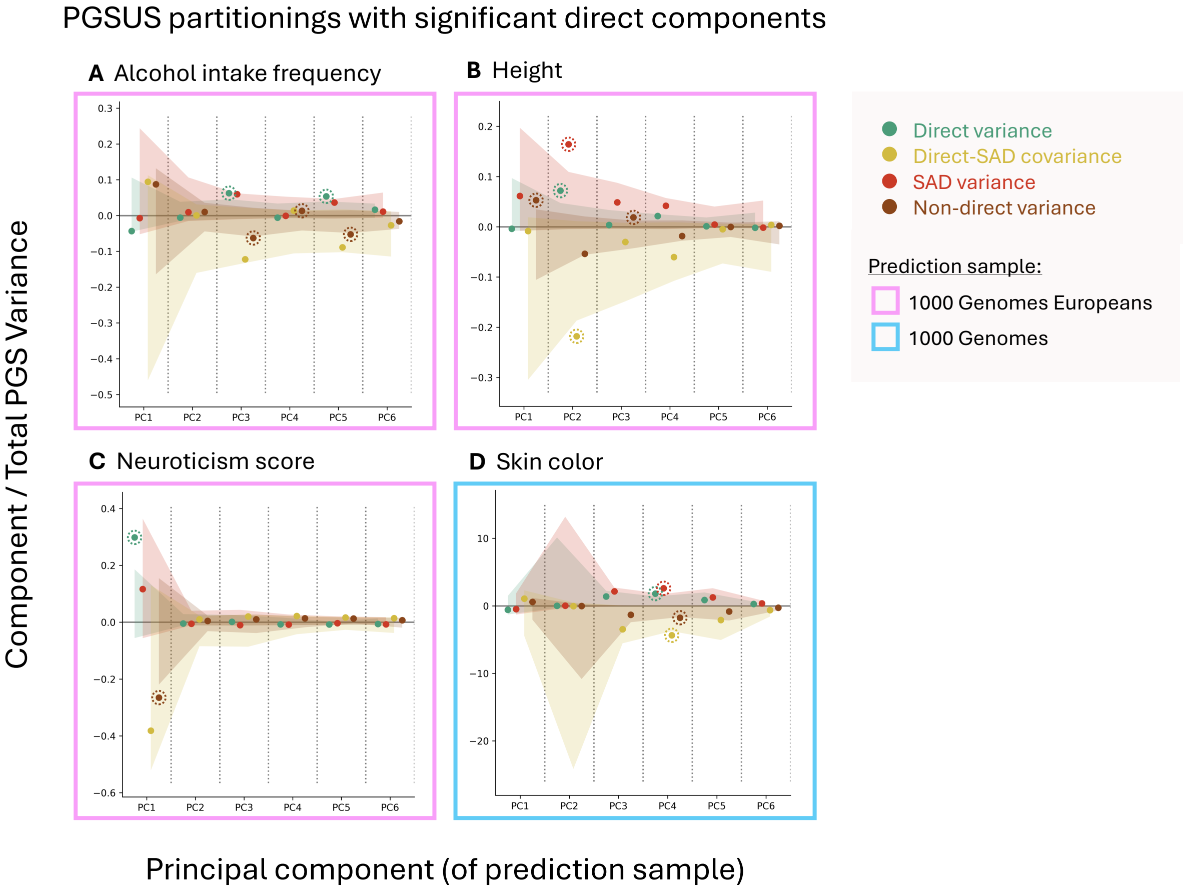

**Figure S27. Examples of significant direct variance components.** PGSUS partitions the variance of a PGS among individuals in a prediction sample. Variance components are attributable to an effect type—direct effects, SAD effects and their covariance—and a principal component of the genotype matrix of the prediction sample. Shown are variance components (divided by the total variance in the PGS) along the first six PCs for four PGSs constructed for the UK Biobank standard-GWAS summary statistics for **(A)** alcohol intake frequency ascertained at $p$-value $< 10^{-8}$, **(B)** height ascertained at $p$-value $< 10^{-5}$, **(C)** neuroticism score ascertained at $p$-value $< 10^{-8}$, and **(D)** skin color ascertained at $p$-value $< 10^{-8}$. Shaded regions show empirical permutation-based null non-rejection regions. Components deviating from this expectation are highlighted with dashed circles.

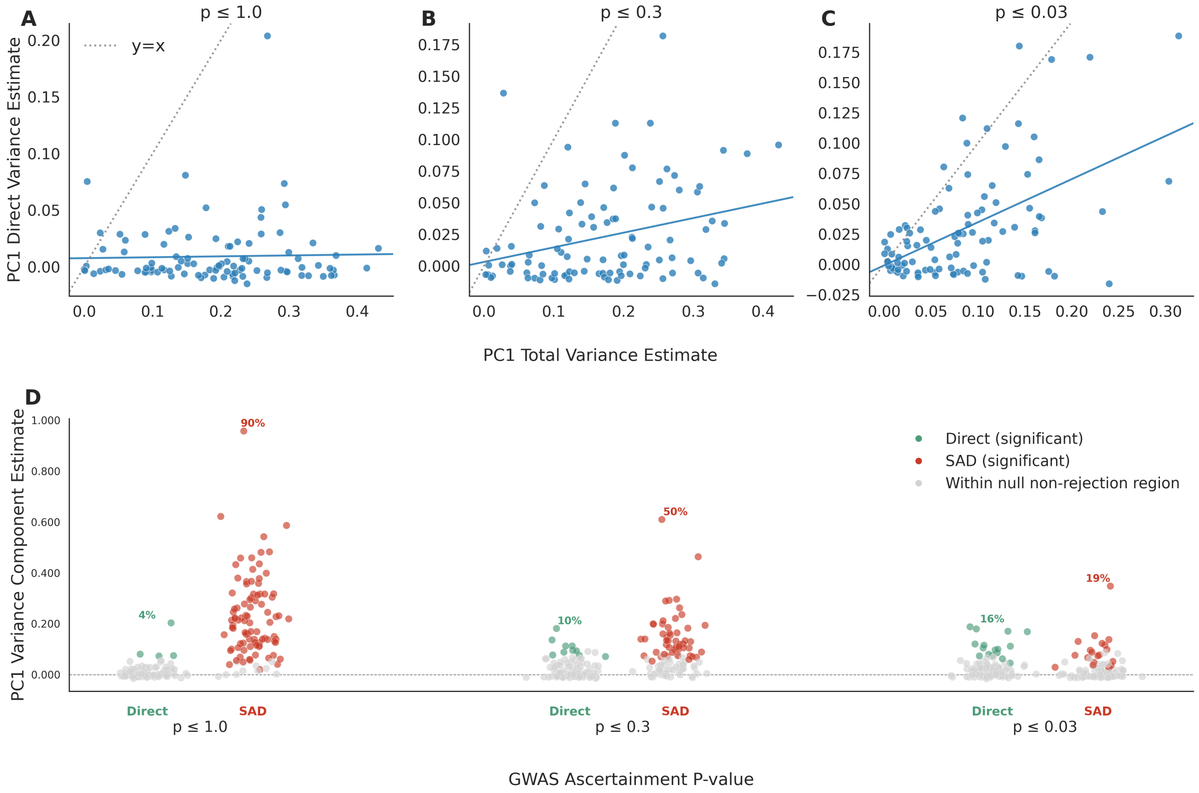

**Figure S28. Collider bias from GWAS ascertainment induces elevated PC1-direct variance components in the presence of stratification.** We simulated a population split ($t = 100$ generations) into two subpopulations with an environmental difference between them ($\delta = 0.6$), predominantly differentiated along PC1 (see **Text S14.4**). **(A-C)** For three different ascertainment thresholds, the x-axis shows the estimated total variance projected onto PC1 (which combines both direct variance and SAD variance) and the y-axis shows the estimated direct variance projected onto PC1. Each data point represents a simulation iteration. **(D)** As standard-GWAS-based ascertainment becomes more stringent, the rejection rate becomes higher for the direct variance component along PC1 and lower for the SAD variance component along PC1. Each data point corresponds to one simulation iteration. Rejection rates are shown as percentages, printed on top of direct (green) and SAD (red) components.

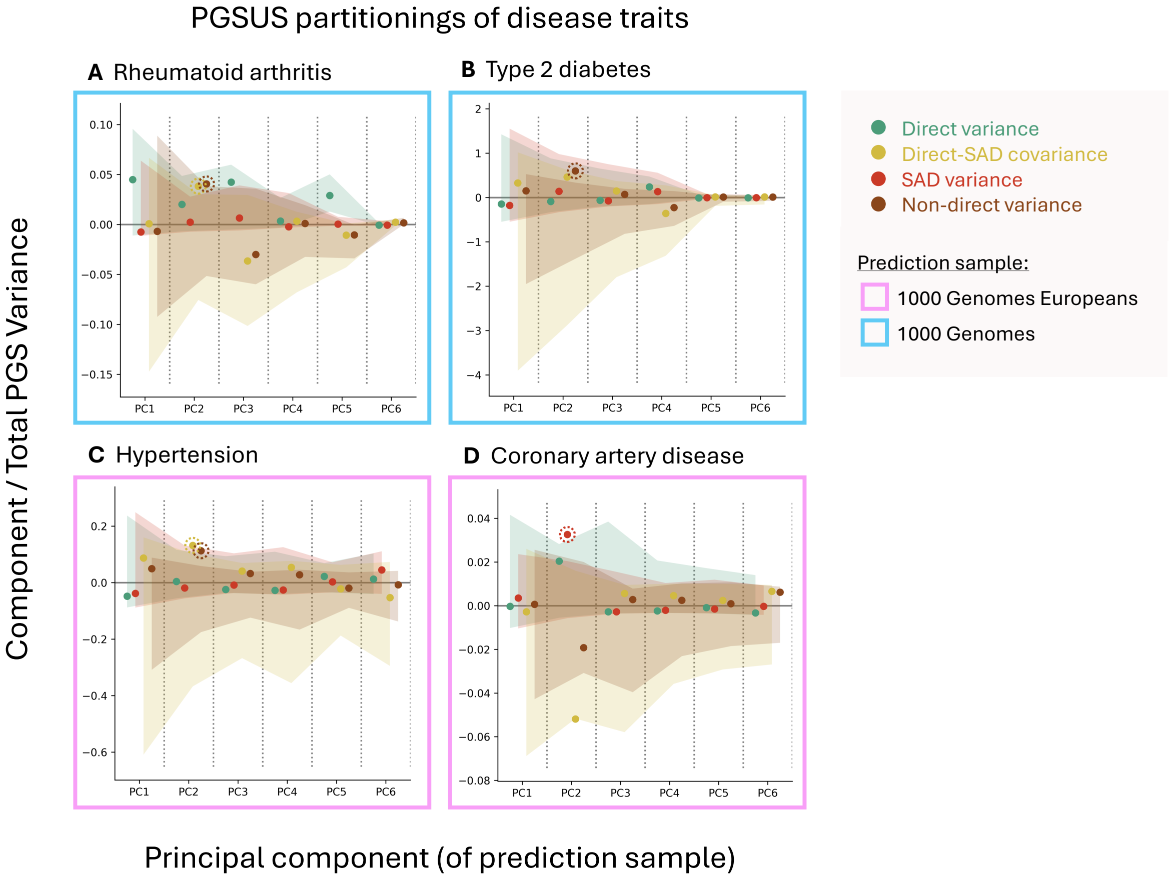

**Figure S29. Variance partitionings for various PGSs for binary health conditions in the UK Biobank.** PGSUS partitions the variance of a PGS among individuals in a prediction sample. Variance components are attributable to an effect type—direct effects, SAD effects and their covariance—and a principal component of the genotype matrix of the prediction sample. Shown are variance components (divided by the total variance in the PGS) along the first six PCs for PGSs constructed for the UK Biobank standard-GWAS summary statistics for four different binary health conditions. Shaded regions show empirical permutation-based null non-rejection regions. Components deviating from this expectation are highlighted with dashed circles. Selected examples show PGSs for binary health conditions with a significant SAD or non-direct component at various ascertainment thresholds: **(A)** rheumatoid arthritis at $p$-value $< 10^{-5}$ and **(B)** type 2 diabetes at $p$-value $< 10^{-3}$ applied to the 1KG prediction sample, and **(C)** hypertension at $p$-value $< 10^{-3}$ and **(D)** coronary artery disease at $p$-value $< 10^{-5}$ applied to the 1KG Europeans prediction sample.

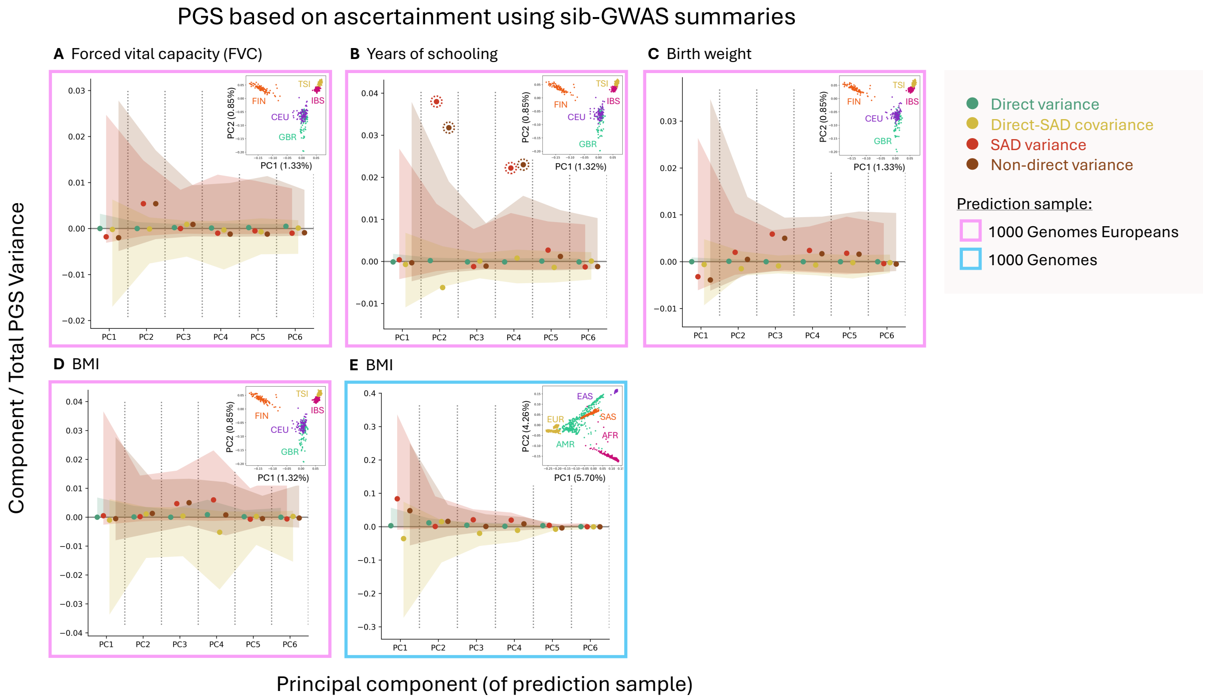

**Figure S30. Variance partitionings of PGSs constructed with ascertainment on allelic effect estimates from sib-GWASs.** PGSUS partitions the variance of a PGS among individuals in a prediction sample. Variance components are attributable to an effect type—direct effects, SAD effects and their covariance—and a principal component of the genotype matrix of the prediction sample. Shown are variance components (divided by the total variance in the PGS) along the first six PCs for five example PGSs corresponding to those in **Fig. 4D-H** but ascertained using sib-GWAS summaries. Shaded regions show empirical permutation-based null non-rejection regions. Components deviating from this expectation are highlighted with dashed circles. Insets show the prediction-sample individual coordinates along the top 2 PCs. Insets vary slightly between the same prediction cohort as they are constructed using the set of SNPs selected through clumping using trait-specific summary statistics. The 1KG subsample labels in panels **(A-D)** correspond to the five European subpopulations: Finnish in Finland (FIN), Utah residents with northern and western European ancestry (CEU), Iberian populations in Spain (IBS), British from England and Scotland (GBR), and Tuscan in Italy (TSI). The 1KG subsample labels in panel **(E)** correspond to the five 1KG superpopulations: European (EUR), admixed Americans (AMR), East Asian (EAS), South Asian (SAS), and African (AFR). Percentages of variance explained by each PC are given in the parentheses of each axis label in the insets. PGS were constructed per the process described in **Text S6**. Due to differences in power between standard and sib-GWASs, we ascertained the $n$ most associated SNPs per PGS, where $n$ is the number of SNPs used to construct the corresponding PGSs from **Fig. 4C-G** (see **Table S2**).

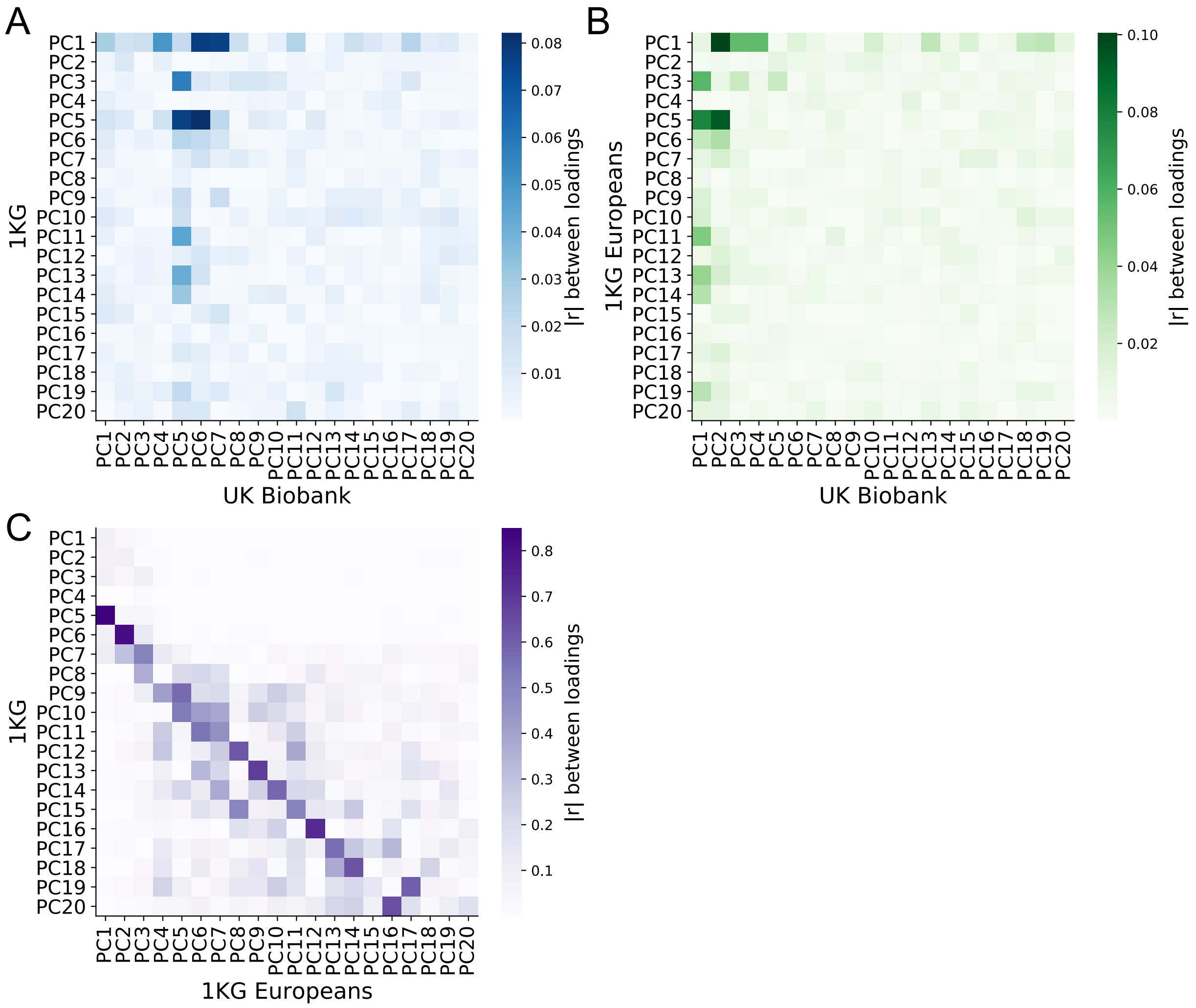

**Figure S31. Correlation among the loadings of individual PCs across considered datasets. (A)** Pearson correlation among PC loadings for the 20 PCs used as covariates in the UKB White British (WB) standard GWAS in the main text and top 1KG cohort PC loadings. **(B)** Pearson correlation among UKB WB PC loadings and top 1KG Europeans PC loadings. **(C)** Pearson correlation among 1KG PC loadings and 1KG Europeans PC loadings.

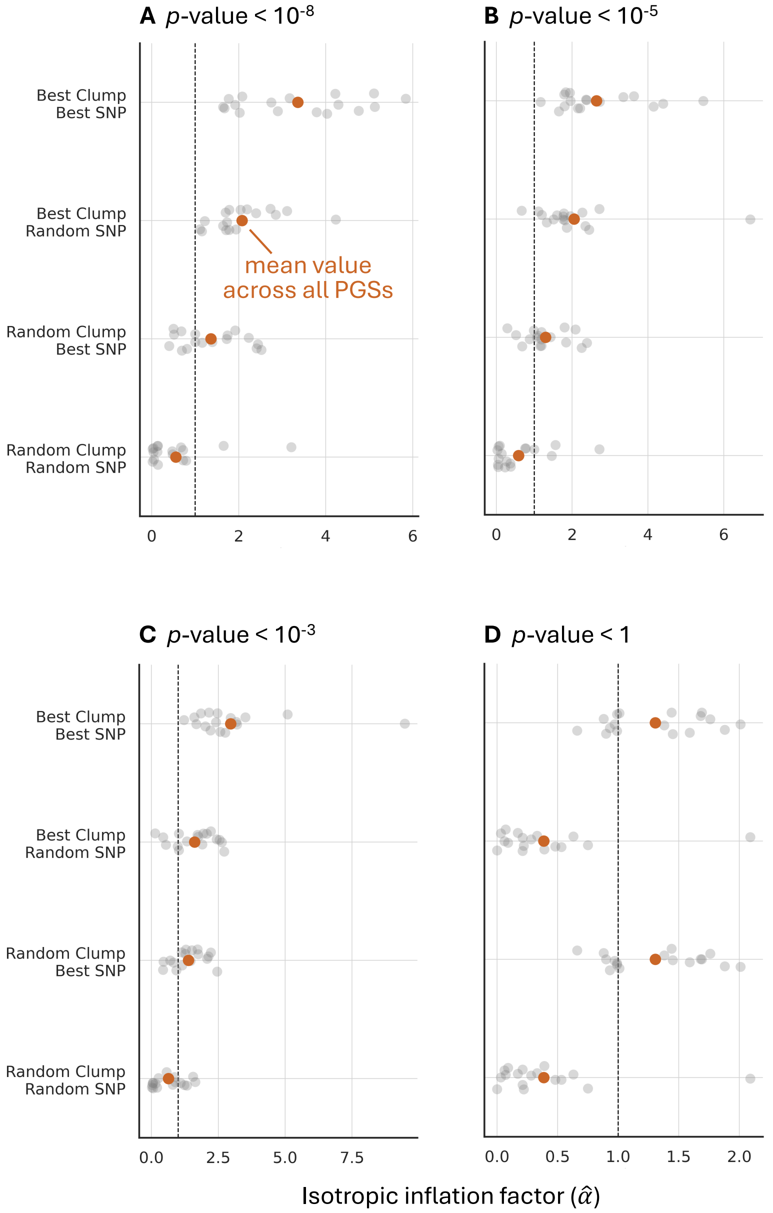

**Figure S32. Effect of SNP selection through clumping and significance thresholding on isotropic inflation in the UK Biobank.** Panels **(A)-(D)** show estimates for the isotropic inflation factor for PGSs for all 17 traits from **Table S1** with index SNPs ascertained at four different ascertainment thresholds using one of the following procedures: standard clumping and thresholding, i.e. choosing the best index SNP of each of the best clumps, i.e., clumps containing the most significantly associated SNPs ("best clump, best SNP"); taking index SNPs to be one random SNP from each of the best clumps ("best clump, random SNP"); taking the index SNPs to be the most strongly associated SNP from each of randomly chosen clumps ("random clump, best SNP"); or random choices of one index SNP per each of randomly chosen clump ("random clump, random SNP"). Each gray point represents one trait and the orange points represent the mean value for that procedure across all PGSs.

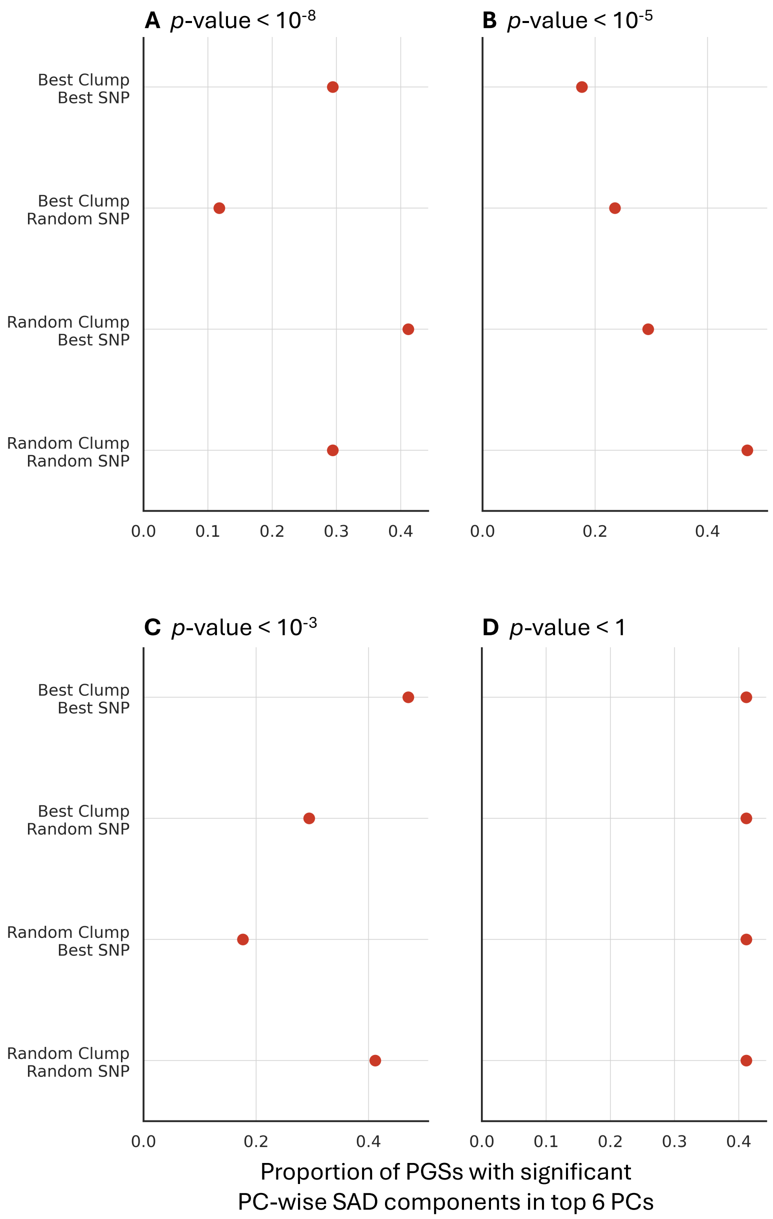

**Figure S33. No systemic effect of index SNP significance thresholding on rates of significant PC-specific SAD variance in the UK Biobank.** Panels **(A)-(D)** show, for PGSs for all 17 traits from **Table S1**, the proportion of PGSs with significant PC-wise SAD variance components. Index SNPs for each PGS were ascertained at four different ascertainment thresholds using one of the following procedures: standard clumping and thresholding, i.e. choosing the best index SNP of each of the best clumps, i.e., clumps containing the most significantly associated SNPs ("best clump, best SNP"); taking index SNPs to be one random SNP from each of the best clumps ("best clump, random SNP"); taking the index SNPs to be the most strongly associated SNP from each of randomly chosen clumps ("random clump, best SNP"); or random choices of one index SNP per each of randomly chosen clump ("random clump, random SNP").

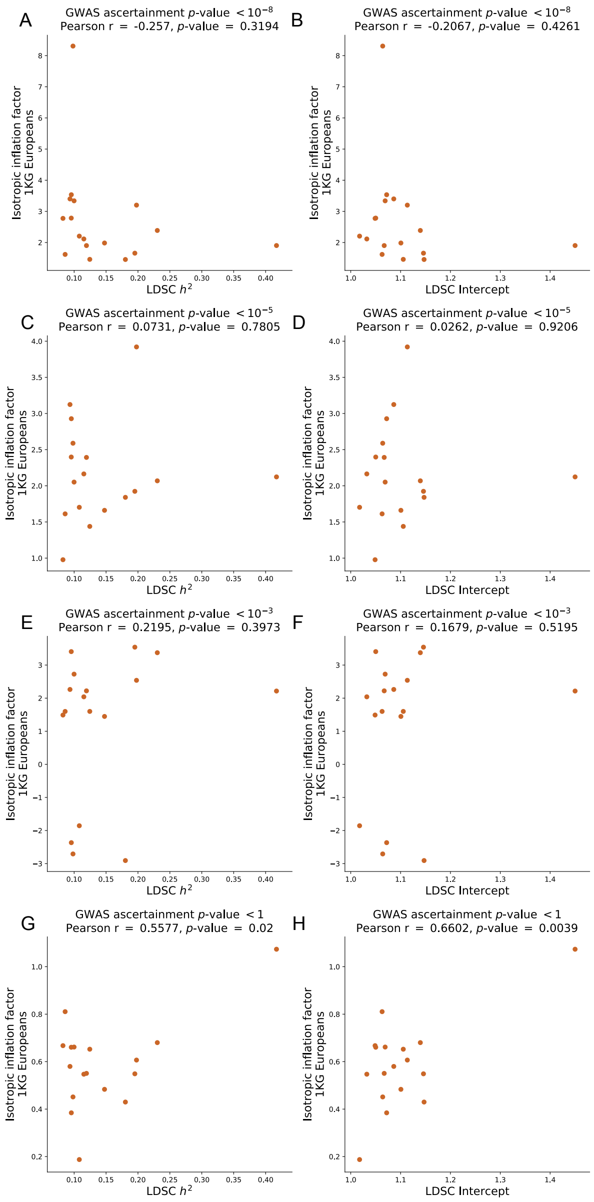

**Figure S34. Correlation between isotropic inflation factor estimates and LDSC-based $h^2$ estimates and intercepts in the 1KG Europeans.** Panels **(A,C,E,G)** show the correlation between LD score regression (LDSC) heritability estimates and estimates of the isotropic inflation factor and panels **(B,D,F,H)** show the correlation between LDSC intercepts and estimates of the isotropic inflation factor for each trait at four ascertainment thresholds. The x-axis in each panel within a column is the same, representing the LDSC additive SNP heritability estimate (left) or the LDSC intercept (right) for each of the 17 traits in **Table S1**.

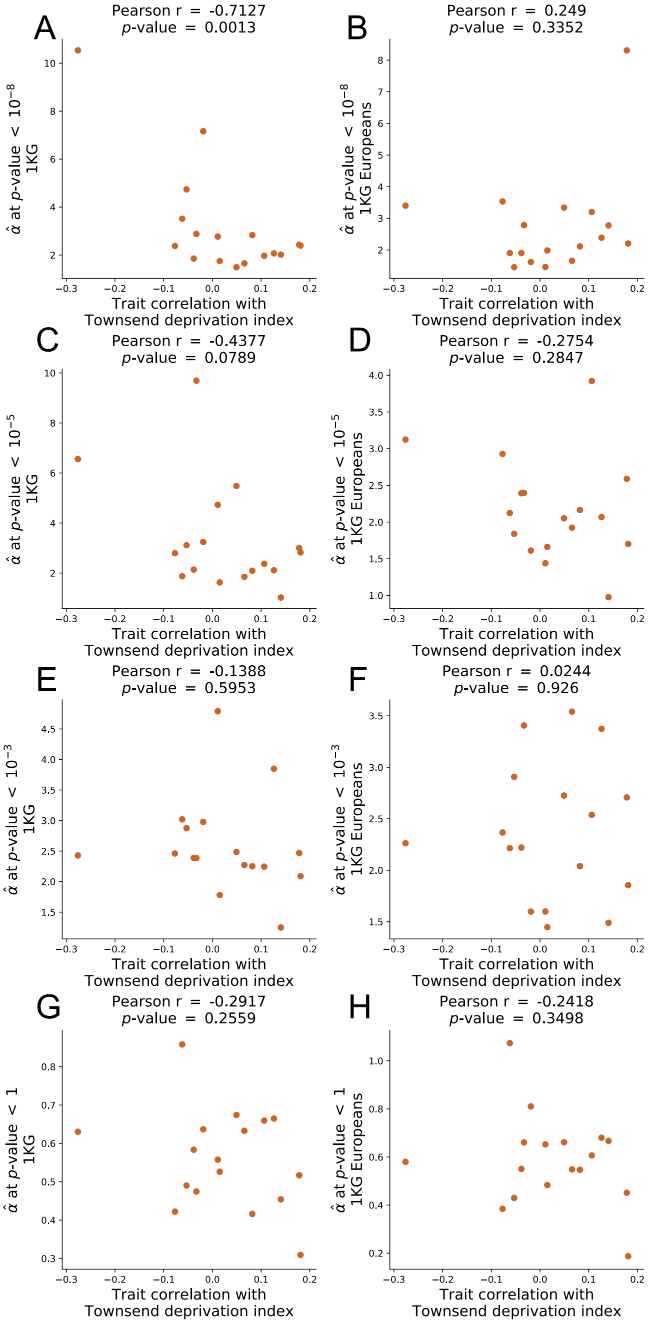

**Figure S35. Correlation between isotropic inflation factor estimates and trait correlation with Townsend Deprivation Index.** For 17 traits, we calculated the correlation between trait values and the Townsend Deprivation Index (TDI) for individuals in the UKB White British cohort, which is shown on the x-axis. For PGSs at four $p$-value ascertainment thresholds for each trait, we show the correlation between these trait-specific correlations with TDI and estimates of the isotropic inflation factor when using either the entire 1KG cohort as the prediction sample (panels **(A,C,E,G)**) or when using the 1KG Europeans as the prediction sample (panels **(B,D,F,H)**).

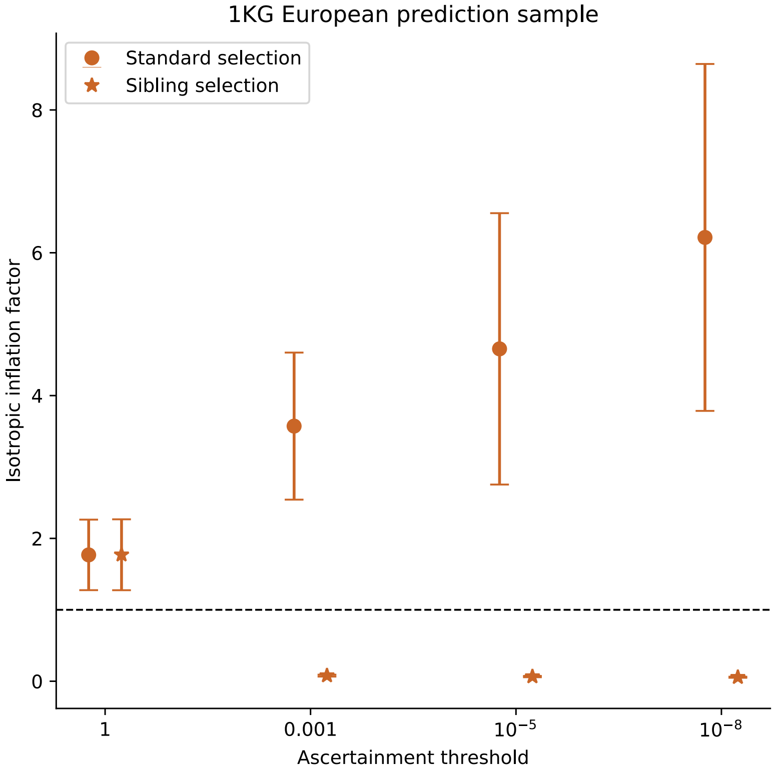

**Figure S36. Effect of winner's curse on isotropic inflation estimates in simulations based on real UK Biobank genotype data.** 100 simulations of trait architecture were performed using genotypes of individuals from the UK Biobank White British cohort and then analyzed using ascertainment based on standard and sib-GWAS $p$-values. 1% of all variants in the UK Biobank data were selected at random to be causal and explain 50% of total trait variance. The remaining trait variance was set to be explained by random environmental effects.

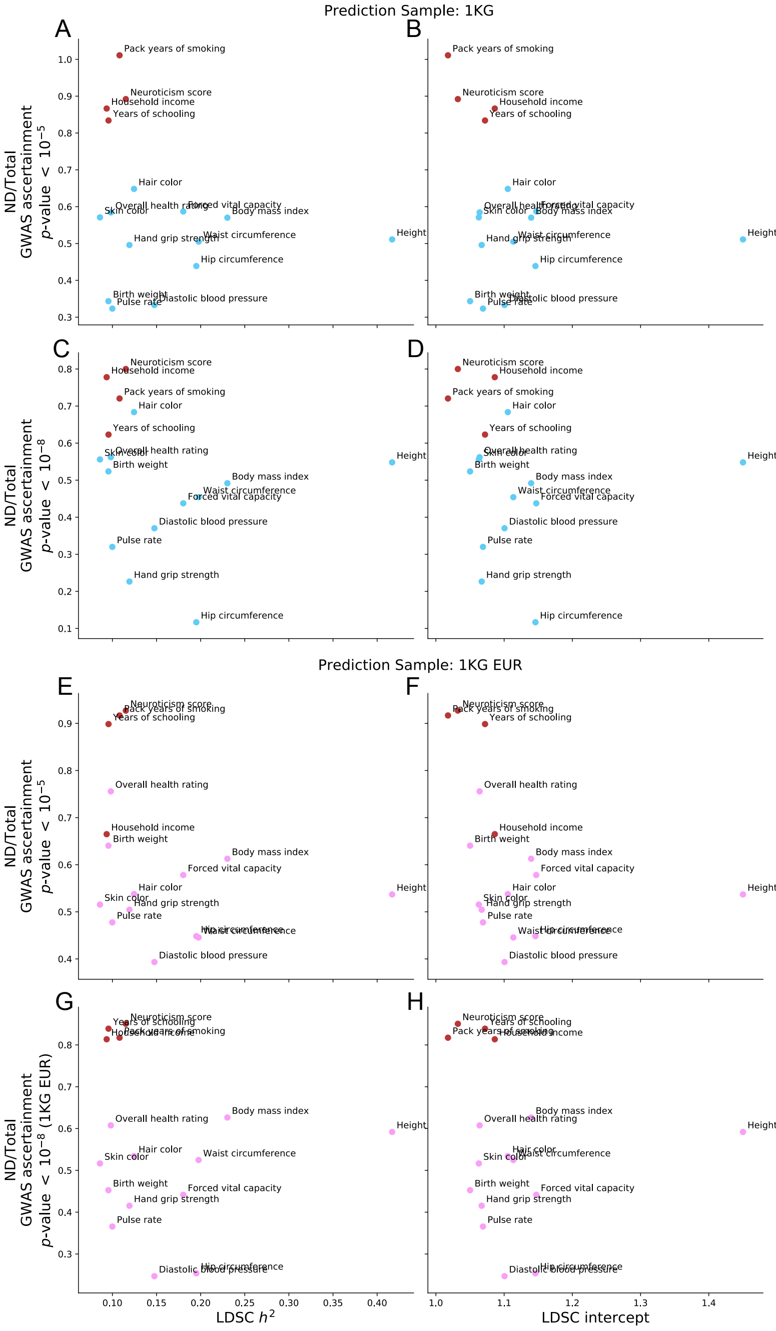

**Figure S37. Comparison of LDSC parameters with the non-direct variance component.** Shown are correlations between the non-direct variance component and the LD score regression (LDSC) SNP heritability estimate (panels **(A,C,E,G)**) or the LDSC intercept (panels **(B,D,F,H)**) for each of the 17 traits estimated based on the UKB WB GWAS we performed at two different ascertainment thresholds. Estimates in panels **(A-D)** are for PGSs applied to the 1KG prediction sample and those in panels **(E-H)** are for PGSs applied to the 1KG Europeans prediction sample. Brown points indicate PGSs for social or behavioral traits.

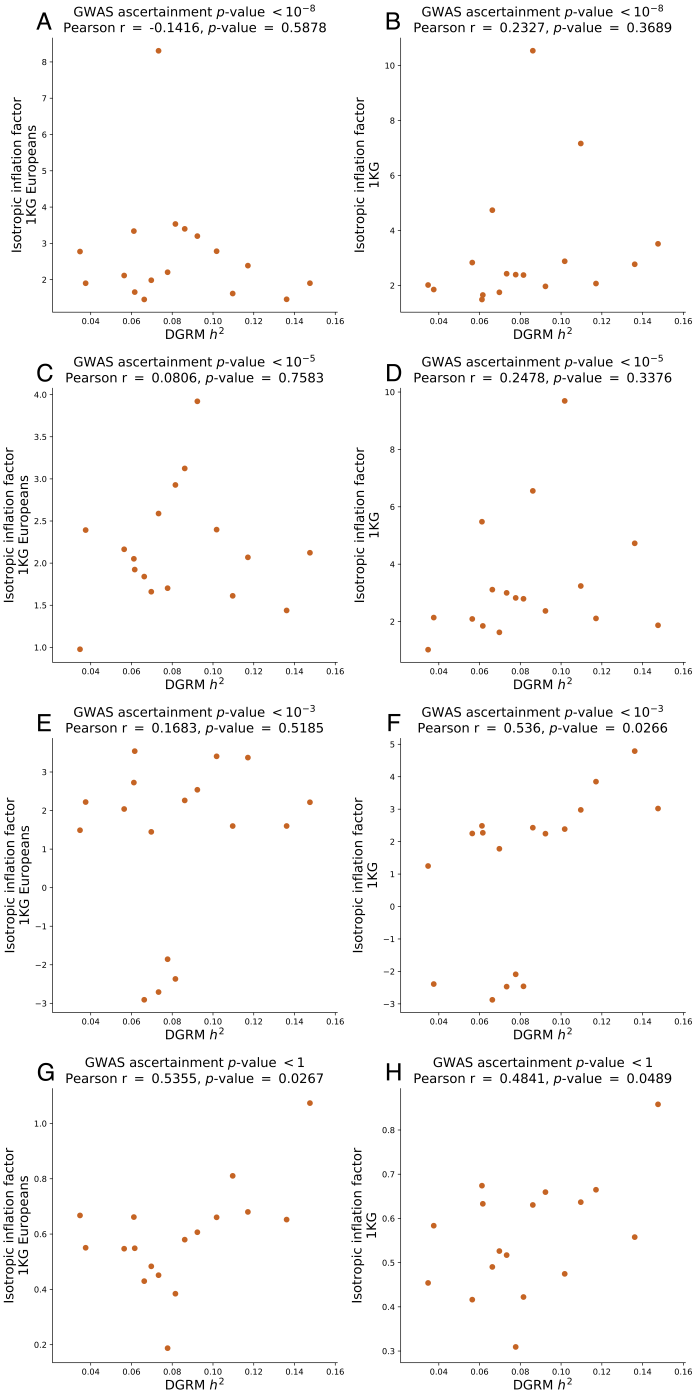

**Figure S38. Comparison of DGREML heritability estimates and isotropic inflation factors for 17 traits in the UK Biobank.** Panels show the relationship between the estimated heritability due to gametic phase disequilibrium estimated by DGREML[50] and estimates of the isotropic inflation factor at four $p$-value ascertainment thresholds when using either **(A,C,E,G)** the 1KG European cohort as the prediction sample or **(B,D,F,H)** the entire 1KG cohort as the prediction sample.

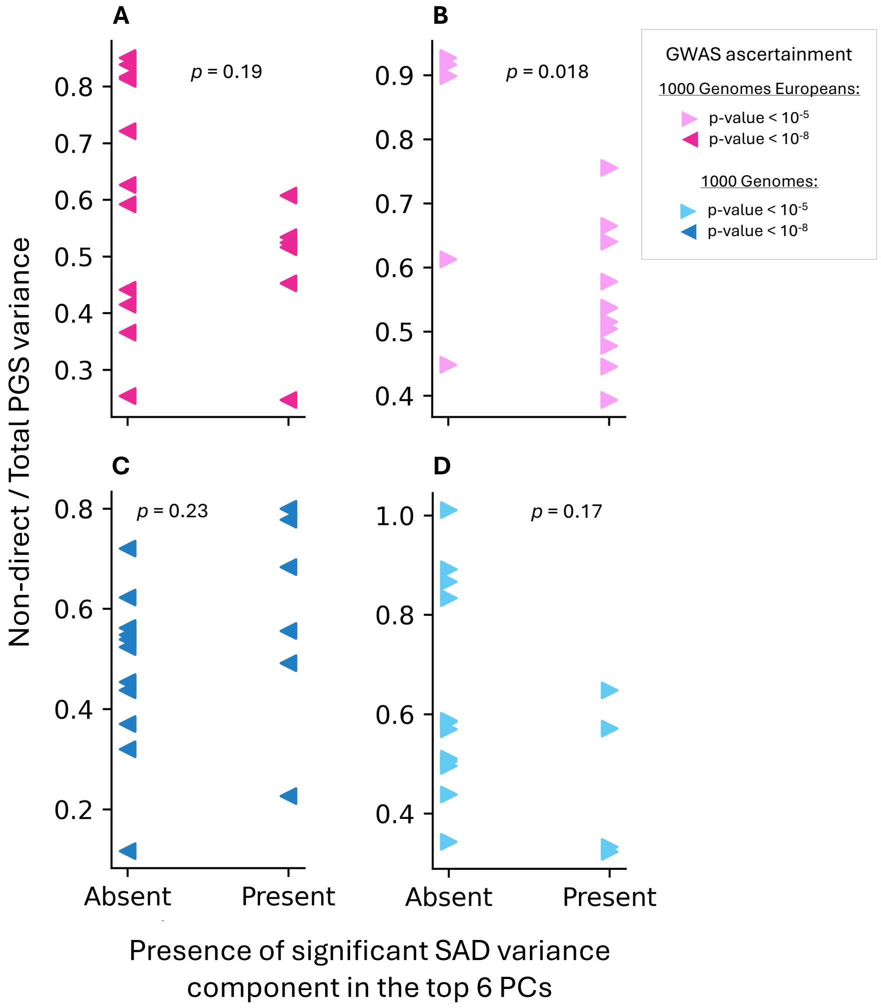

**Figure S39. The correlation between the presence of PC-specific confounding and the non-direct variance component.** We evaluated the correlation between the presence of a significant SAD variance component in the top six PCs and the non-direct variance component for 34 PGSs based on GWAS in the UKB WB for the 17 traits in **Table S1**, applied to both the 1KG Europeans **(A,B)** and 1KG **(C,D)** cohorts at two GWAS ascertainment thresholds. $p$-values are for t-tests of the mean difference in estimates of the non-direct variance components between PGSs with and without significant PC-specific SAD variance components among the top six PCs.

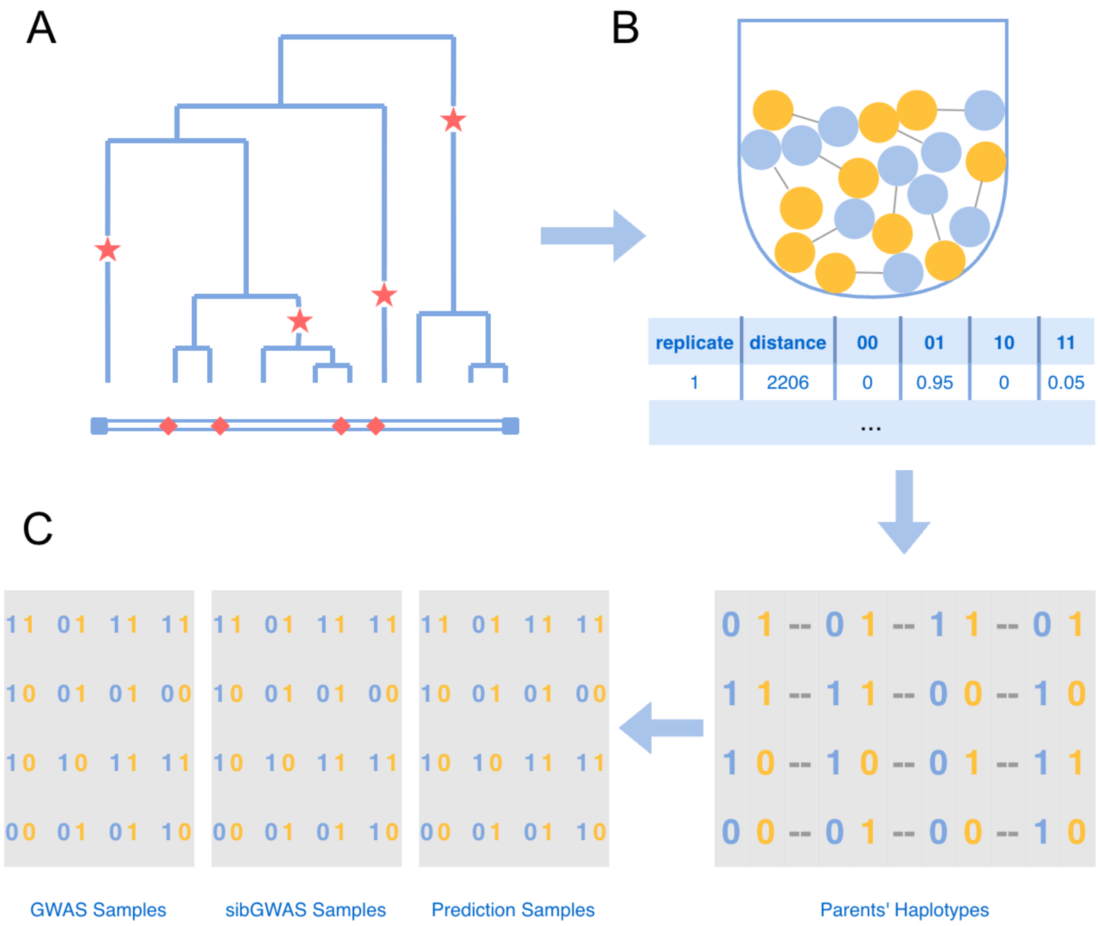

**Figure S40. Generation of genotype data in simulations. (A)** Marker-causal variant pairs were simulated across independent 500-kb genomic regions using *msprime*[51]. Mutations arising on distinct lineages of the coalescent tree are represented by stars; their corresponding physical positions on the genomic region are represented by diamonds. In this example, 4 mutations yields 6 unique pairs (i.e., $\binom{4}{2}=6$), which were then evaluated for allele frequencies and linkage disequilibrium. **(B)** A bucket serves as a schematic of the pool of possibilities from which marker-causal variant pairs are sampled, and an example entry from the marker-causal variants library is displayed below. Each row corresponds to a unique pair and records the physical distance (bp) between variants and the four observed haplotype frequencies $(f_{00}, f_{01}, f_{10}, f_{11})$. **(C)** The rightmost matrix shows the simulated parental genomic data, where each row represents a single haploid genome (with 1,000 marker-causal pairs) in an individual and each pair of columns represent a marker-causal variants pair. At each genomic position, a set of haplotype frequencies was randomly sampled without replacement from the library, and haplotypes were assigned through proportional sampling from the recorded frequencies. Parental individuals were mated to produce an offspring population. Each parent produced gametes by drawing from their own haplotypes at each locus, with recombination occurring with a probability determined by the physical distance between loci multiplied by a constant recombination rate of $1 \times 10^{-8}$ per base pair. The offspring data were used to construct non-overlapping samples for standard GWAS, sib-GWAS and prediction.

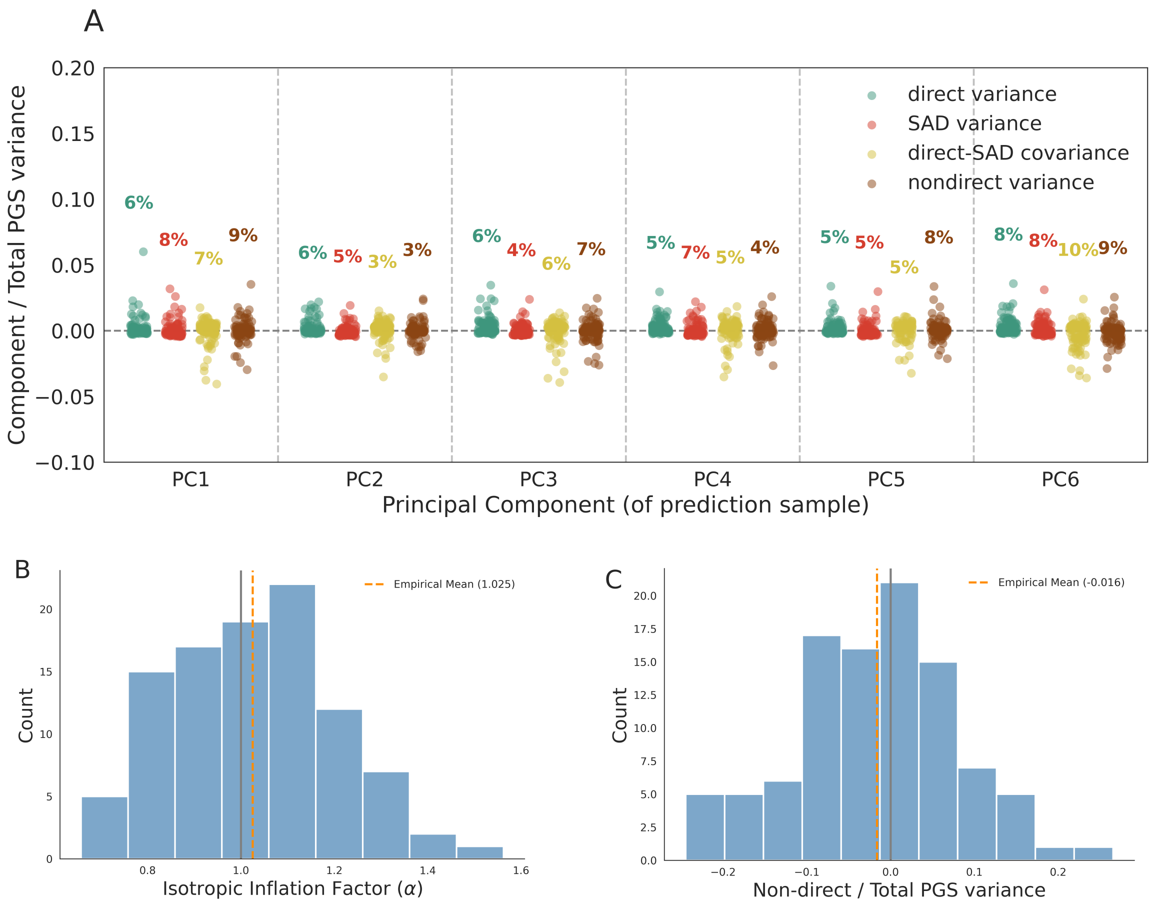

**Figure S41. PC-specific variance components, isotropic inflation factors, and non-direct variance components under the null model.** We ran PGSUS on simulated genetic data in a scenario with no SAD effects. **(A)** Within each column, four color-coded jitter groups represent the distribution of specific variance components observed across 100 iterations. Percentages above each group represent the proportion of iterations where the variance falling outside the permutation-based null non-rejection region, which is defined by the 0th and 95th percentile of empirical null samples for direct and SAD components, and the 2.5th and 97.5th percentiles for covariance and non-direct components. Across the first six PCs, the average rejection rate for each variance component is 0.06. **(B)** The mean value of the estimated isotropic inflation factor across 100 iterations under the null scenario is approximately 1. **(C)** The mean estimated non-direct variance component across 100 iterations under the null scenario is near zero.

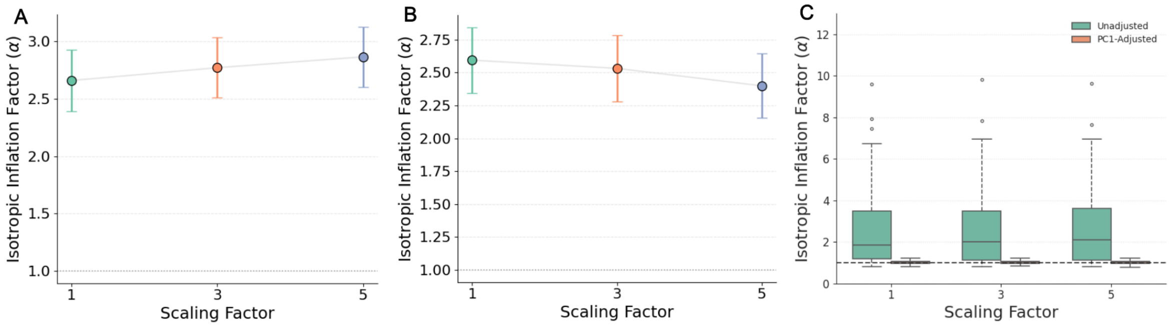

**Figure S42. Isotropic inflation in a demographic model where environment and genetic relatedness covary.** We estimated the isotropic inflation factor under a two-population demographic model in which the covariance of environmental components among individuals is proportional to the product of their genetic relatedness and a scaling factor (shown on the x-axis; **Text S14.7**). **(A)** Simulation iterations in which the environmental difference between the two populations has the same sign as the genetic difference between them. **(B)** Simulation iterations in which the environmental difference and genetic difference have opposite signs. **(C)** Including PC1 from standard-GWAS sample as an adjustment for population stratification led to the average isotropic inflation factor returning to approximately one.

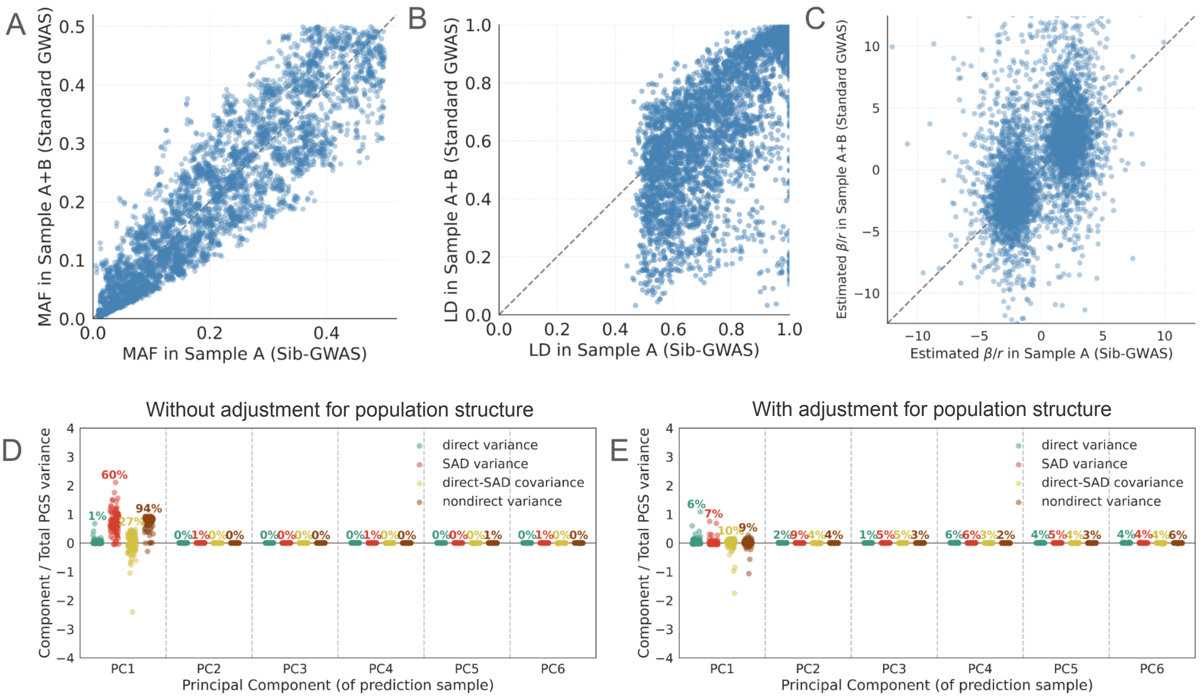

**Figure S43. Ancestry-biased sampling leads to incongruity between standard- and sib-GWAS estimates in simulated data.** **(A)** Comparison of the minor allele frequencies (MAF) for the top 50 SNPs with the largest allelic effects across 100 iterations in population A versus the mixed population comprised of equal numbers of individuals from populations A and B. **(B)** Comparison of local LD (measured by $r^2$) for the top 50 SNPs with the largest allelic effect across 100 iterations in population A and mixed population A+B. **(C)** Comparison of the LD-adjusted estimated causal allelic effect for the top 50 SNPs with the largest allelic effects across 100 iterations in population A versus the mixed population A+B. **(D)** Variance components for the top six PCs without adjustment to the standard GWAS for population structure. PC1-SAD variance and the frequency with which it falls outside the null non-rejection region are markedly elevated because standard-GWAS estimates capture underlying population structure. **(E)** Adjusting for population structure by including population labels as a covariate in the standard GWAS reduces PC1 SAD variance and rejection rate, but they remain elevated compared with the baseline case.

### Supplemental Tables

| Trait | Standard-GWAS Sample Size (Individuals) | Sib-GWAS Sample Size (Pairs) | UK Biobank Field ID |
|---|---|---|---|
| Alcohol intake frequency | 321,718 | 17,328 | 1558 |
| Birth weight | 184,380 | 6,569 | 20022 |
| BMI | 320,898 | 17,257 | 21001 |
| Diastolic blood pressure | 293,726 | 14,717 | 4079, 6153, 6177 |
| Forced vital capacity | 293,657 | 14,522 | 3062 |
| Hair color | 321,309 | 17,292 | 1747 |
| Hand grip strength | 320,373 | 17,178 | 46, 47 |
| Height | 321,251 | 17,284 | 50 |
| Hip circumference | 321,371 | 17,297 | 49 |
| Household income | 277,310 | 13,125 | 738 |
| Neuroticism score | 221,552 | 8,358 | 20127 |
| Overall health rating | 320,823 | 17,234 | 2178 |
| Pack years of smoking | 96,942 | 2,163 | 20161 |
| Pulse rate | 300,744 | 15,415 | 102 |
| Skin color | 318,056 | 16,924 | 1717 |
| Waist circumference | 321,407 | 17,301 | 48 |
| Years of schooling | 264,131 | 12,076 | 6138 |
| Townsend deprivation index | 275,796 | NA | 22189 |

**Table S1. Data field IDs and sample sizes for standard and sib-GWAS in the UK Biobank.** Sample sizes for the 17 traits described in the **Methods** section. Summary statistics were obtained using a standard regression in the standard-GWAS analyses and a sibling regression in pairs of full siblings in the sib-GWAS. Additionally included are the pertinent numbers for the Townsend Deprivation Index analyzed in **Fig. S35**.

| **Trait** | **p-value < 1** | **p-value < $10^{-3}$** | **p-value < $10^{-5}$** | **p-value< $10^{-8}$** |
|---|---|---|---|---|
| Alcohol intake frequency | 352,878 | 4,226 | 400 | 49 |
| Birth weight | 353,093 | 3,277 | 337 | 62 |
| BMI | 352,760 | 9,676 | 1,980 | 477 |
| Diastolic blood pressure | 353,166 | 6,059 | 1,080 | 262 |
| Forced vital capacity | 352,657 | 7,707 | 1,552 | 412 |
| Hair color | 353,040 | 5,389 | 1,637 | 848 |
| Hand grip strength | 352,824 | 5,394 | 670 | 117 |
| Height | 352,287 | 17,888 | 6,301 | 2,739 |
| Hip circumference | 352,942 | 8,404 | 1,623 | 423 |
| Household income | 352,950 | 4,293 | 363 | 37 |
| Neuroticism score | 352,961 | 4,273 | 391 | 50 |
| Overall health rating | 352,797 | 4,654 | 446 | 42 |
| Pack years of smoking | 353,036 | 2,301 | 88 | 11 |
| Pulse rate | 352,811 | 4,484 | 675 | 216 |
| Skin color | 353,446 | 4,085 | 929 | 531 |
| Waist circumference | 352,885 | 8,417 | 1,500 | 313 |
| Years of schooling | 352,940 | 4,165 | 334 | 35 |

**Table S2. Number of PGS index SNPs at four different ascertainment $p$-value thresholds.** Index SNP counts for PGSs at four different ascertainment thresholds for the traits in **Table S1**.

| Trait | Standard-GWAS Cases | Standard-GWAS Controls | Sib-GWAS Both Cases (Pairs) | Sib-GWAS One Case (Pairs) | Sib-GWAS No Cases (Pairs) | ICD10 code(s) |
|---|---|---|---|---|---|---|
| Asthma | 2,373 | 317,835 | 158 | 263 | 16,578 | J45 |
| Coronary artery disease | 17,738 | 302,470 | 237 | 1,609 | 15,132 | I25 |
| Cataract | 25,083 | 295,125 | 308 | 2,211 | 14,459 | H26 |
| Hypertension | 1,870 | 318,338 | 155 | 205 | 16,639 | I10 |
| Obesity (BMI $\geq$ 30) | 77,553 | 243,345 | 1,447 | 5,081 | 10,729 | – |
| Rheumatoid arthritis | 1,713 | 318,495 | 148 | 184 | 16,664 | M06 |
| Type 1 diabetes | 628 | 319,580 | 151 | 73 | 16,774 | E10 |
| Type 2 diabetes | 1,342 | 318,866 | 152 | 122 | 16,724 | E11 |

**Table S3. ICD10 codes and sample sizes for standard and sib-GWAS of binary health conditions in the UK Biobank.** Sample sizes for the 8 binary health conditions as described in **Text S6**. Individuals were categorized as obese using the definition of BMI $\geq$ 30 (UKB data field 21001).

| Trait | p-value < 1 | p-value < $10^{-3}$ | p-value < $10^{-5}$ | p-value< $10^{-8}$ |
|---|---|---|---|---|
| Asthma | 375,092 | 1,667 | 51 | 3 |
| Coronary artery disease | 374,770 | 2,663 | 186 | 50 |
| Cataract | 374,864 | 1,989 | 73 | 5 |
| Hypertension | 375,403 | 1,654 | 37 | 1 |
| Obesity | 374,248 | 6,378 | 928 | 192 |
| Rheumatoid arthritis | 374,957 | 1,850 | 94 | 23 |
| Type 1 diabetes | 375,169 | 1,990 | 119 | 33 |
| Type 2 diabetes | 375,438 | 1,806 | 41 | 1 |

**Table S4. Number of PGS index SNPs for binary health conditions.** Index SNP counts for PGSs at four different ascertainment thresholds for the binary health traits in **Table S3**.

### Supplemental File Descriptions

**File S1.** Contains the significant direct (d), SAD (s), and covariance (c) components at four different ascertainment thresholds for the standard *PLINK* GWASs with age, sex, and the first 20 GWAS cohort PCs as covariates in the entire 1KG ("1KG All") and 1KG Europeans prediction samples.

**File S2.** Contains the significant direct (d), SAD (s), covariance (c), and non-direct (n) components at four different ascertainment thresholds for the standard *PLINK* GWASs with age and sex alone as covariates in the entire 1KG ("1KG All") and 1KG Europeans prediction samples.

**File S3.** Contains the significant direct (d), SAD (s), covariance (c), and non-direct (n) components at four different ascertainment thresholds for the standard *PLINK* GWASs with age, sex, and the first 20 PCs from the *prediction* sample as covariates in the entire 1KG ("1KG All") cohort.

**File S4.** Contains the significant direct (d), SAD (s), covariance (c), and non-direct (n) components at four different ascertainment thresholds for the standard *PLINK* GWASs with age, sex, and the first 20 PCs from the *prediction* sample as covariates in the 1KG Europeans cohort.

**File S5.** Contains the significant direct (d), SAD (s), covariance (c), and non-direct (n) components at four different ascertainment thresholds for the standard *PLINK* GWASs with age, sex, the first 20 GWAS cohort PCs, and the first 20 PCs from the *prediction* sample as covariates in the entire 1KG ("1KG All") cohort.

**File S6.** Contains the significant direct (d), SAD (s), covariance (c), and non-direct (n) components at four different ascertainment thresholds for the standard *PLINK* GWASs with age, sex, the first 20 GWAS cohort PCs, and the first 20 PCs from the *prediction* sample as covariates in the 1KG Europeans cohort.

**File S7.** Contains the significant direct (d), SAD (s), covariance (c), and non-direct (n) components at four different ascertainment thresholds for the standard *BOLT-LMM* GWASs in the entire 1KG ("1KG All") and 1KG Europeans prediction samples using the GRM alone to correct for population structure.

**File S8.** Contains the significant direct (d), SAD (s), covariance (c), and non-direct (n) components at four different ascertainment thresholds for the standard *BOLT-LMM* GWASs in the entire 1KG ("1KG All") and 1KG Europeans prediction samples using the GRM and the first 20 GWAS cohort PCs as fixed effects to control for population structure.

**File S9.** Contains statistics and $p$-values from paired t-tests (isotropic inflation factor, non-direct variance component) and McNemar's tests (PC-specific SAD variance components) evaluating the effect of GWAS adjustment methods to adjust for confounding on PGSUS-derived estimates with a random subsample of the UKB as the GWAS sample.

**File S10.** Contains statistics and $p$-values from paired t-tests (isotropic inflation factor, non-direct variance component) and McNemar's tests (PC-specific SAD variance components) evaluating the effect of GWAS adjustment methods to adjust for confounding on PGSUS-derived estimates with the UKB White British cohort as the GWAS sample.

8. Po-Ru Loh, George Tucker, Brendan K Bulik-Sullivan, Bjarni J Vilhjálmsson, Hilary K Finucane, Rany M Salem, Daniel I Chasman, Paul M Ridker, Benjamin M Neale, Bonnie

Berger, et al. Efficient Bayesian mixed-model analysis increases association power in large cohorts. *Nature Genetics*, 47(3):284–290, 2015. doi: 10.1038/ng.3190.

9. Andries T Marees, Hilde De Kluiver, Sven Stringer, Florence Vorspan, Emmanuel Curis, Cynthia Marie-Claire, and Eske M Derks. A tutorial on conducting genome-wide association studies: Quality control and statistical analysis. *International Journal of Methods in Psychiatric Research*, 27(2):e1608, 2018.

10. Shing Wan Choi, Timothy Shin-Heng Mak, and Paul F O'Reilly. Tutorial: a guide to performing polygenic risk score analyses. *Nature Protocols*, 15(9):2759–2772, 2020. doi: 10.1038/s41596-020-0353-1.

11. Scott Hazelhurst, Noah Zaitlen, Bogdan Pasaniuc, Bjarni Vilhjálmsson, and Shaun Aron. Medical population genetics and GWAS for complex diseases 19th april - 24th april 2015 training material archive, March 2022.

30. Tammy Tan, Hariharan Jayashankar, Junming Guan, Seyed Moeen Nehzati, Mahdi Mir, Michael Bennett, Esben Agerbo, Rafael Ahlskog, Ville Pinto de Andrade Anapaz, Bjørn Olav Åsvold, Stefania Benonisdottir, Laxmi Bhatta, Dorret I. Boomsma, Ben Brumpton, Archie Campbell, Christopher F. Chabris, Rosa Cheesman, Zhengming Chen, China Kadoorie Biobank Collaborative Group, Eco de Geus, Erik A. Ehli, Abdelrahman G. Elnahas, Estonian Biobank Research Team, Finngen, Andrea Ganna, Alexandros Giannelis, Liisa Hakaste, Ailin Falkmo Hansen, Alexandra Havdahl, Caroline Hayward, Jouke-Jan Hottenga, Mikkel Aagaard Houmark, Kristian Hveem, Jaakko Kaprio, Arnulf Langhammer, Antti Latvala, James J. Lee, Mikko Lehtovirta, Liming Li, LifeLines Cohort Study, Kuang Lin, Richard Karlsson Linnér, Stefano Lombardi, Nicholas G. Martin, Matt McGue, Sarah E. Medland, Andres Metspalu, Brittany L. Mitchell, Guiyan Ni, Ilja M. Nolte, Matthew T. Oetjens, Sven Oskarsson, Teemu Palviainen, Rashmi B. Prasad, Anu Reigo, Kadri Reis, Julia Sidorenko, Karri Silventoinen, Harold Snieder, Tiinamaija Tuomi, Bjarni J. Vilhjálmsson, Robin G. Walters, Emily A. Willoughby, Bendik S. Winsvold, Eivind Ystrom, Jonathan Flint, Loic Yengo, Peter M. Visscher, Augustine Kong, Elliot M. Tucker-Drob, Richard Border, David Cesarini, Patrick Turley, Aysu Okbay, Daniel J. Benjamin, and Alexander Strudwick Young. Family-gwas reveals effects of environment and mating on genetic associations. 2024. doi: 10.1101/2024.10.01.24314703.

31. Yuchang Wu, Xiaoyuan Zhong, Yunong Lin, Zijie Zhao, Jiawen Chen, Boyan Zheng, James J Li, Jason M Fletcher, and Qiongshi Lu. Estimating genetic nurture with summary statistics of multigenerational genome-wide association studies. *Proceedings of the National Academy of Sciences*, 118(25):e2023184118, 2021. doi: 10.1073/pnas.2023184118.

32. Laurence J Howe, Michel G Nivard, Tim T Morris, Ailin F Hansen, Humaira Rasheed,

Yoonsu Cho, Geetha Chittoor, Rafael Ahlskog, Penelope A Lind, Teemu Palviainen, et al. Within-sibship genome-wide association analyses decrease bias in estimates of direct genetic effects. *Nature Genetics*, 54(5):581–592, 2022.

60. Jennifer Blanc and Jeremy J Berg. Testing for differences in polygenic scores in the presence of confounding. *bioRxiv*, 2023.

61. Aysu Okbay, Jonathan P Beauchamp, Mark Alan Fontana, James J Lee, Tune H Pers, Cornelius A Rietveld, Patrick Turley, Guo-Bo Chen, Valur Emilsson, S Fleur W Meddens, et al. Genome-wide association study identifies 74 loci associated with educational attainment. *Nature*, 533(7604):539–542, 2016. doi: 10.1038/nature17671.

62. Qiaomei Fu, Mateja Hajdinjak, Oana Teodora Moldovan, Silviu Constantin, Swapan Mallick, Pontus Skoglund, Nick Patterson, Nadin Rohland, Iosif Lazaridis, Birgit Nickel, Bence Viola, Kay Prüfer, Matthias Meyer, Janet Kelso, David Reich, and Svante Pääbo.

An early modern human from Romania with a recent Neanderthal ancestor. *Nature*, 524: 216–219, 2015. doi: 10.1038/nature14558.

63. Wolfgang Haak, Iosif Lazaridis, Nick Patterson, Nadin Rohland, Swapan Mallick, Bastien Llamas, Guido Brandt, Susanne Nordenfelt, Eadaoin Harney, Kristin Stewardson, Qiaomei Fu, Alissa Mittnik, Eszter Bánffy, Christos Economou, Michael Francken, Susanne Friederich, Rafael Garrido Pena, Fredrik Hallgren, Valery Khartanovich, Aleksandr Khokhlov, Michael Kunst, Pavel Kuznetsov, Harald Meller, Oleg Mochalov, Vayacheslav Moiseyev, Nicole Nicklisch, Sandra L. Pichler, Roberto Risch, Manuel A. Rojo Guerra, Christina Roth, Anna Szécsényi-Nagy, Joachim Wahl, Matthias Meyer, Johannes Krause, Dorcas Brown, David Anthony, Alan Cooper, Kurt Werner Alt, and David Reich. Massive migration from the steppe was a source for Indo-European languages in Europe. *Nature*, 522:207–211, 2015. doi: 10.1038/nature14317.

64. Iain Mathieson, Iosif Lazaridis, Nadin Rohland, Swapan Mallick, Nick Patterson, Songül Alpaslan Roodenberg, Eadaoin Harney, Kristin Stewardson, Daniel Fernandes, Mario Novak, Kendra Sirak, Cristina Gamba, Eppie R. Jones, Bastien Llamas, Stanislav Dryomov, Joseph Pickrell, Juan Luís Arsuaga, José María Bermúdez de Castro, Eudald Carbonell, Fokke Gerritsen, Aleksandr Khokhlov, Pavel Kuznetsov, Marina Lozano, Harald Meller, Oleg Mochalov, Vyacheslav Moiseyev, Manuel A. Rojo Guerra, Jacob Roodenberg, Josep Maria Vergès, Johannes Krause, Alan Cooper, Kurt W. Alt, Dorcas Brown, David Anthony, Carles Lalueza-Fox, Wolfgang Haak, Ron Pinhasi, and David Reich. Genome-wide patterns of selection in 230 ancient Eurasians. *Nature*, 528:499–503, 2015. doi: 10.1038/nature16152.

65. Nadin Rohland, Swapan Mallick, Matthew Mah, Robert Maier, Nick Patterson, and David Reich. Three assays for in-solution enrichment of ancient human DNA at more than a million SNPs. *Genome Research*, 32:2068–2078, 2022. doi: 10.1101/gr.276728.122.

66. Ludovic Orlando, Robin Allaby, Pontus Skoglund, Clio der Sarkissian, Philipp W Stockhammer, María C Ávila Arcos, Qiaomei Fu, Johannes Krause, Eske Willerslev, Anne C Stone, and Christina Warinner. Ancient DNA analysis. *Nature Reviews Methods Primers*, 1, 2021. doi: 10.1038/s43586-020-00011-0.
